# Engineered yeast cells simulating CD19+ cancers to control CAR T cell activation

**DOI:** 10.1101/2023.10.25.563929

**Authors:** Marcus Deichmann, Giovanni Schiesaro, Keerthana Ramanathan, Katrine Zeeberg, Nanna M. T. Koefoed, Maria Ormhøj, Rasmus U. W. Friis, Ryan T. Gill, Sine R. Hadrup, Emil D. Jensen, Michael K. Jensen

## Abstract

Chimeric antigen receptor (CAR) T cells have become an established immunotherapy and show promising results for the treatment of hematological cancers. However, modulation of surface levels of the targeted antigen in cancer cells affects the quality and safety of CAR T cell therapy. Here we present the <u>S</u>ynthetic <u>C</u>ellular <u>A</u>dvanced <u>S</u>ignal <u>A</u>dapter (SCASA) system, based on successful engineering of yeast to simulate cancer cells with tunable surface-antigen densities, as a tool for controlled activation of CAR T cell responses and assessment of antigen density effects. Specifically, we demonstrate I) controllable antigen-densities of CD19 on yeast using G protein-coupled receptors (GPCRs), II) a customizable system allowing choice of signal input and modular pathway engineering for precise fine-tuning of the output, III) synthetic cell-cell communication with CAR T cells and the application of CD19-displaying yeast in the characterization of CAR designs, and IV) more efficient and robust activational control of clinically-derived CAR T cells in comparison to the NALM6 cancer cell line. Based on this yeast-based antigen-presenting cell system, we envision efficient assessment of how varying antigen densities in cancer cells affect CAR T cell responses and ultimately support development of safer and better quality of personalized cancer therapies.

## Introduction

Over the past two decades, synthetic biology has enabled new generations of cell therapies through novel genetic engineering strategies and programmable gene circuits to achieve controlled and therapeutically-balanced functions in both microbes and human cells^1^. Cellular immunotherapies show promising clinical results for the treatment of cancers, notably with chimeric antigen receptor (CAR) technology, which commonly relies on autologous T cells engineered to heterologously express a synthetic CAR construct enabling T cell anticancer responses targeted towards cancer-associated antigens (e.g. CD19), upon patient re-infusion^2,3^. Since 2017, six FDA approvals on CAR products for hematological cancers have been registered^3^, including anti-CD19 CAR T cells for relapsed or refractory (r/r) B cell acute lymphoblastic leukemia (B-ALL), showing complete remission (CR) in 70-96% of patients^4–13^.

However, while remission rates are encouraging, 14-57% of r/r B-ALL patients obtaining CR from CAR T cell therapy eventually experience relapse^4–13^. Some relapses are caused by immunosuppression, exhaustion, or poorly persistent and ineffectual CAR T cells^13–16^, and others are attributed directly to escape via antigen modulation by cancer cells, where CAR T cells may remain functional, but simply lack sufficient targetable antigen^17^. CD19 modulation occurs through a variety of mechanisms^17–25^ and is thought to be the primary mechanism of tumor escape in r/r B-ALL, occurring in 7-28% of r/r B-ALL CRs, is a major determinant for response durability, and is associated with poor prognosis^3–9,12,13,26–29^.

Even gradual modulation that causes decrease in the cancer-cell-surface antigen density can increase resistance to therapy, as CAR T cell responses are graded to antigen density^25,30–39^. CAR T cell efficiency and behavior are directly influenced, as distinct anticancer responses are triggered at different antigen-density thresholds and have density-dependent intensities. For example, cytolytic activity requires a lower antigen density than CAR T cell proliferation and cytokine release^30–33,35,36^. These antigen density-dependent effects have differential impact across CAR designs and CAR expression levels^25,30,31,34,35,37,38^, emphasizing the importance of strategic CAR design to mitigate antigen modulation, as well as proper characterization of the hundreds of novel CARs for new antigens and cancers undergoing development and clinical trials^40,41^. Antigen-density model systems continue to improve the understanding of these dynamics, such as cell lines with varying antigen densities that closely mimic the complexity of *in vivo* cell-to-cell interactions^25,30–37,42–44^, as well as non-cellular materials embedded with antigens that enable greater antigen-density control and orthogonality^31,38,39,45–49^. Complementing these approaches, engineered yeast cells offer a highly versatile alternative.

The concept of using engineered yeast cells (*Saccharomyces cerevisiae*) as T-cell response screening platforms through co-cultivation emerged in literature in 1998 with the advent of yeast-surface display (YSD) technology^50,51^. Since then, yeast has become an applied tool in deciphering T-cell receptor (TCR) specificities and peptide-loaded major histocompatibility complexes (pMHC)^52–54^, and co-cultivation has been employed to verify pMHC-display functionality and directly assay TCR-specific T-cell activation^55–60^. In this context, yeast platforms benefit from a well-developed genetic engineering toolbox and an extensive gene-circuit parts repertoire for efficient and versatile customizability. Furthermore, DNA-encoding in yeast supports high-throughput interaction screening of libraries and directed evolution of human proteins. More practically, the availability and simplicity of implementation, as well as robustness and fast growth as a self-replicating system, makes yeast a powerful and cost-effective platform for synthetic interspecies cell-cell communication^52,56,59,61,62^.

In this study, we leverage these traits to develop a controllable antigen-density model system, and build cancer-simulating CD19+ yeast to study the activation dynamics of human CAR T cells *ex vivo*. To achieve this goal, we create a fully genome-encoded cellular platform, named the <u>S</u>ynthetic <u>C</u>ellular <u>A</u>dvanced <u>S</u>ignal <u>A</u>dapter (SCASA) system, which can control surface-displayed CD19 at levels spanning >3 orders of magnitude and is compatible with T cell co-cultivation with no inherent effect on viability, proliferation, or activation of donor-derived T cells or cell lines. This allows dynamic control of CAR-specific human immune cell activation at levels on par with the standard NALM6 cancer cell line. Further, we apply SCASA yeast cells to characterize and discriminate antigen-dependent effects on signaling pathways in different CAR designs, and lastly, to functionally assess a novel donor-derived CAR T cell product, with an activation efficiency and robustness in antigen density and target cell numbers beyond what cancer cell lines currently support.

## Results

### Designing the Synthetic Cellular Advanced Signal Adapter (SCASA) system

In the interest of establishing a versatile platform technology for evaluating antigen-density effects in immune cell responses, we envisioned the Synthetic Cellular Advanced Signal Adapter (SCASA) system allowing yeast to surface-display proteins (*the effector module*) at levels regulated by an engineered version of the well-characterized and highly modular yeast pheromone response pathway (PRP)^63–67^ (*the processing module)*, from a sensed amount of a chosen extracellular G protein-coupled receptor (GPCR) ligand presented to the cells (*the sensory module*) (**Fig. 1a**). Ideally, such a design comprises controllability of display density, tunability of signal processing dynamics, orthogonality to T cell activation, and customizability in the types and intensity of sense-response functions, which would be favorable for adapting the SCASA system to user-defined cell assays and health applications^1,52^ (**Fig. 1b**).

**Figure 1.**
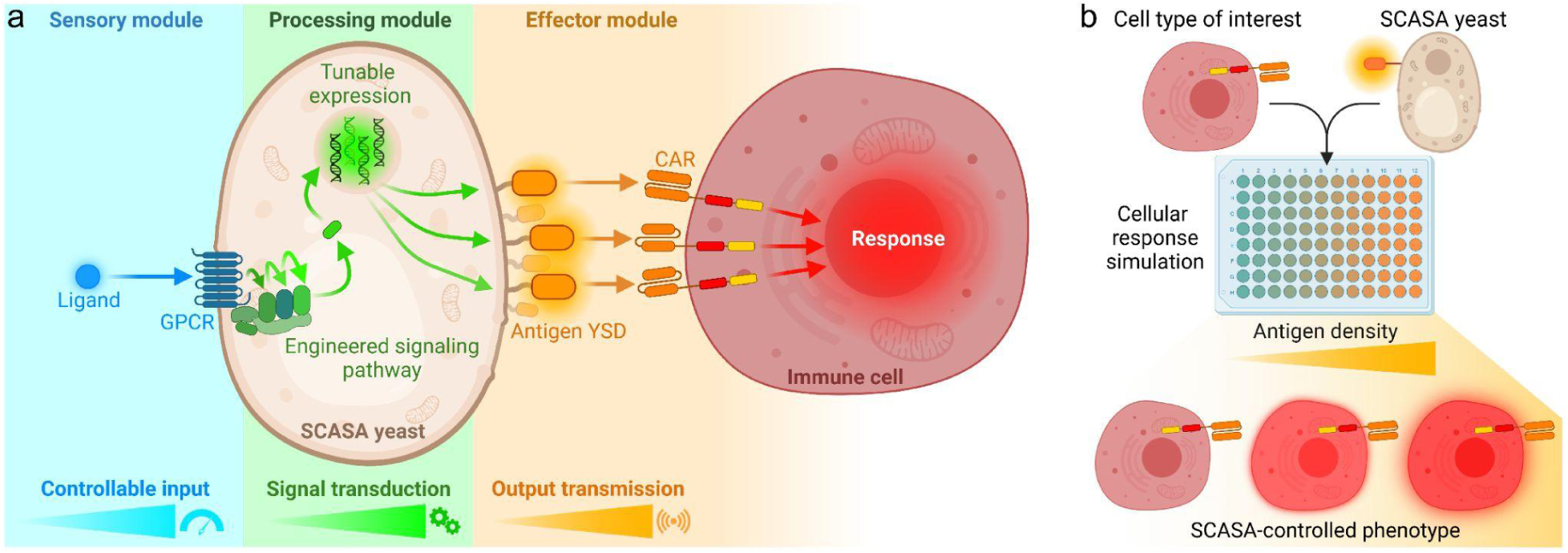
Outline of the Synthetic Cellular <u>A</u>dvanced <u>S</u>ignal <u>A</u>dapter (SCASA) system. **a.** SCASA yeast cells engineered to present tunable dosages of antigens via yeast surface display (YSD) on the cell surface (*effector module*), proportional to the intracellular signal transduced via an engineered pheromone response pathway (*processing module*) with controlled activation and intensity by G protein-coupled receptors (GPCRs) based on cognate ligands presented to the cells (*sensory module*). Each module is customizable to adapt to the specific application of the SCASA yeast cells. **b.** Example of a co-cultivation layout in which engineered SCASA yeast cells are used to simulate different antigen densities of a CAR T cell target to obtain gradual intensities of CAR T cell responses.

### CD19 yeast surface display for cancer cell simulation

In our first design for antigen display, we employed a highly mutated CD19 extracellular domain (CD19.1 ECD) in the Aga2-fusion display construct for correct folding and expression in yeast^68,69^, hereafter merely denoted CD19 (**Fig. 2**). Functional CD19 display was confirmed via staining with FMC63-clone anti-CD19 antibody containing the binding domain of current FDA-approved CAR T cell designs^3^. YSD was optimized by overexpression of the Aga1 anchor and deletion of native Aga2 (*aga2Δ0*) to avoid competitive binding to Aga1. To both increase antigen-display efficiency and limit inconsistent behavior in antigen-displaying populations, genetic display cassettes were genomically integrated rather than expressed from a plasmid^59,70–72^. Ultimately, this yielded 99.7±0.3% CD19+ yeast cells, clearly exemplified by the P_TDH3_ promoter, thus closely resembling the phenotype of the cancer benchmark CD19+ NALM6 B cell leukemia cell line (100±0% CD19+) (**Fig. 2, Suppl. Fig. 1**). We additionally confirmed the increased efficiency via galactose-induced CD19 expression from P_GAL1_ (**Suppl. Fig. 2**), though this traditional system was not further pursued due to its limitation to a single input and limited customizability^51,73–75^.

**Figure 2.**
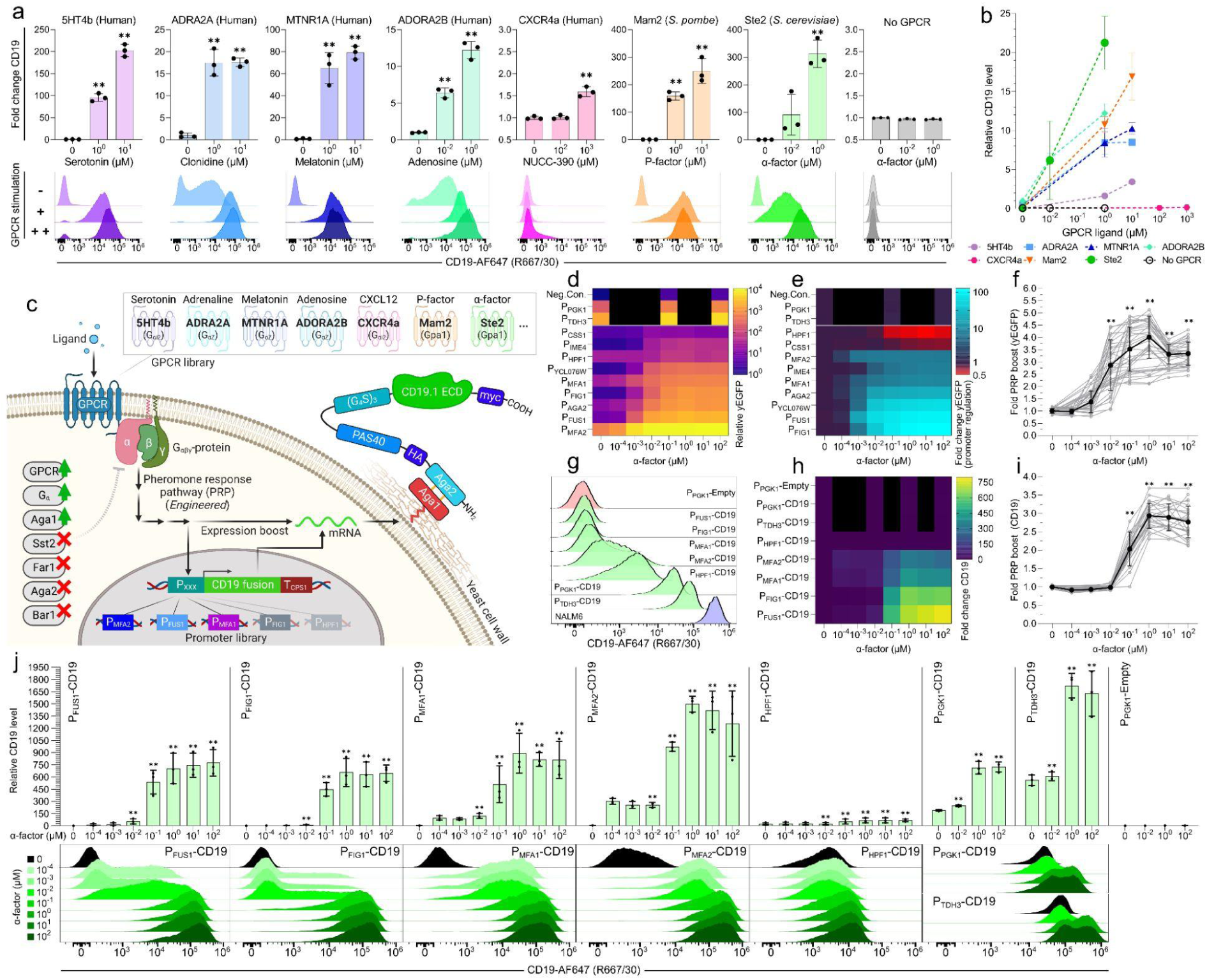
Characterization of SCASA system tunable CD19 antigen presentation by yeast cells. **a.** Heterologous GPCRs as sensory modules for controlling CD19 display with cognate agonists: fold change CD19 levels (*top row*) and absolute CD19 histograms (*lower row*). All strains employ P_FUS1_ and vary between GPCRs and PRP-coupling G_ɑ_-subunits. **b.** Comparison of CD19 levels for all GPCRs. **c.** CD19 SCASA yeast cell design; CD19 expression is controllable by GPCR-dependent ligands (*sensory module*). System regulation depends on GPCR signaling through G_ɑβγ_-protein to an engineered PRP, choice of promoter, and expression boost effects (*processing module*). CD19 output is a fusion protein composed of HA- and myc-tags, PAS40-linker, (G_4_S)_3_-linker, and CD19.1 ECD, fused to Aga2 (*effector module*). Strains were optimized by; G_ɑ_-subunit, GPCR, and Aga1 overexpression, as well as native gene knock-outs; *ste2Δ0, ste3Δ0, gpa1Δ0, sst2Δ0, bar1Δ0, far1Δ0, aga2Δ0*. **d**. Relative yEGFP levels of individual promoters with ɑ-factor stimulation (0-100 µM), including PRP boost effects, sorted; low to high. **e**. Fold change promoter induction (yEGFP), excluding PRP boost effects (SSC-normalization), sorted; low to high. **f**. Approximated PRP boost of yEGFP expression from quantified PRP activation (*black*) across all strains and designs (*grey*) (n=27). **g**. CD19 histograms of SCASA yeast library with different promoters without GPCR stimulation (*green*), a P*_PGK1_*-Empty control lacking CD19 (*red*), and NALM6 (*blue*). **h.** Fold change CD19 of SCASA yeast library during GPCR stimulation with ɑ-factor (0-100 µM). **i.** Approximated PRP boost of CD19 display from quantified PRP activation (*black*), across all strains and designs (*grey*) (n=15). **j.** Comparison of CD19 levels during GPCR stimulation of SCASA yeast library, relative to lowest detected CD19 level (*top row*), and CD19 histograms (*lower row*). Quantification by flow cytometry for CD19 (20 hrs. post-induction, R667/30) or yEGFP (5 hrs. post-induction, B525/45). Unless otherwise noted, data is means of cell counts or median fluorescence intensities (mMFI) for biological replicates (n=3) and standard deviations hereof. Histograms are representative replicates normalized to the mode. Statistical tests: One- and Two-way ANOVA with multiple comparisons statistical tests. Significance levels: ✱: P≤0.05, ✱✱: P≤0.001. Not all pairwise comparisons are shown. Statistics and extended analyses in: **Suppl. Fig. 1-12** and **Suppl. Table 1-6**.

### Sensory modules using heterologous GPCRs allow for customizable input

To empower SCASA yeast cells with versatile and customizable input-sensing capabilities, we next assessed if the sensory module could comprise GPCRs coupled to an engineered yeast pheromone response pathway (PRP)^64,65,67^ (**Fig. 2a-c**). The extracellular sensing of GPCRs displays exceptional diversity in the types of specific ligands, ranging from light rays and small molecules to peptides and large complex proteins^63–67,76,77^. Accordingly, the constructed strain library encoded individual heterologously expressed GPCRs for sensing a diverse set of ligands, with CD19 expression controlled by the PRP-regulated promoter P_FUS1_ ^78^ (**Fig. 2c**). Upon administration of GPCR agonists to the SCASA yeast cells, all designs showed controlled functional up-regulation of CD19 levels (p<0.0001) (**Fig. 2a, Suppl. Fig. 3**). In the tested concentration ranges, the highest absolute CD19 levels and fold-changes were provided by fungal peptide-sensing GPCRs Ste2 (ɑ-factor) at 312.5±50.0-fold and Mam2^64^ (P-factor) at 249.4±44.9-fold (**Fig. 2a-b**), while human GPCRs sensing small-molecule neurotransmitters showed lower absolute CD19 levels (**Fig. 2b**) and fold changes with 5HT4b^79^ (serotonin) at 202.6±13.8-fold, MTNR1A^64^ (melatonin) at 79.2±5.9-fold, ADRA2A^65^ (adrenaline) at 17.6±0.9-fold, and ADORA2B^64^ (adenosine) at 12.2±1.2-fold (**Fig. 2a**). Lastly, the human chemokine-sensing CXCR4a^80^ (CXCL12/SDF1) showed a modest 1.6±0.1-fold up-regulation of CD19 levels (**Fig. 2a**). The GPCRs caused significantly different unstimulated background CD19 levels, notably with ADORA2B causing a 14.0±0.8-fold background increase (p≤0.0013) and 5HT4b lowering the background to 0.24±0.02-fold the level of cells lacking GPCRs (p<0.0001) (**Fig. 2a**).

In conclusion, the sensory module was customizable, as exemplified by the individual expression of six heterologous GPCRs enabling the adoption of small molecule-, peptide-, and protein-based input signals, and providing a 1,249-fold span in surface-displayed CD19 levels across the examined range of GPCR stimulation.

### An engineered pheromone response pathway enables a customizable processing module

As CAR T cell products show different sensitivities to changes in antigen densities^25,30,31,34,35,37,38^, a high degree of adaptability of SCASA cell designs is advantageous to enable simulation of the relevant ranges of antigen densities for a specific CAR design. In this regard, PRP-coupled GPCR signaling in yeast has several valuable attributes, such as highly sensitive concentration-dependent analog signal transduction and a large dynamic output range, combined with its engineerability characterized by high modularity and tunability^63–65,67^ (**Fig. 2**). To generate well-characterized processing module parts for tuning surface display levels, we characterized 11 candidate promoters through yEGFP-expression (**Fig. 2d-e**). The promoters were selected from our recent transcriptome analysis of GPCR-mediated PRP activation^63^, showing expression ranging from undetectable mRNA levels to the most highly expressed gene in *S. cerevisiae*, and spanning 6 orders of magnitude in mRNA abundance upon GPCR stimulation (**Suppl. Fig. 4, Suppl. Table 1**). The processing module promoter library spanned a 1,025-fold range in basal yEGFP output intensity (**Fig. 2d, Suppl. Fig. 5**), and following ligand-mediated GPCR stimulation, the individual promoters displayed further up- or downregulated dynamics (**Fig. 2e**, **Suppl. Fig. 6**), providing a 6,866-fold span in total library output range (**Fig. 2d**). Characterization of yEGFP revealed that output dynamics were highly dependent on the choice of promoter, emphasized by response variation being attributed mainly to promoter choice (SV=75.2%), rather than GPCR stimulation level (SV=13.4%) (**Fig. 2d-e**, **Suppl. Table 2**). Output was boosted by an overall post-transcriptional promoter-independent multiplicative amplification of expression, also observed in similar systems^64^, at an intensity dependent on the degree of yeast mating phenotype induction (p<0.0001) (**Fig. 2f**). This amplification coincided with shmooing, a morphological key indicator of PRP activation^63^ (**Suppl. Fig. 7**), and was dependent on the main yeast mating response transcription factor Ste12^81^ (p<0.0001)(**Suppl. Fig. 8**). In this case, yEGFP expression was generally boosted by up to 4.0±0.9-fold (**Fig. 2f**), exemplified by the regulated output for P_PGK1_ and P_TDH3_ in response to GPCR stimulation (**Fig. 2d, Suppl. Fig. 9**), despite these genes showing no up-regulation of mRNA levels (**Suppl. Fig. 4**, **Suppl. Table 1**), and commonly being regarded as constitutive promoters^63,82^.

Based on these learnings, we generated a yeast strain library to simulate the antigen heterogeneity of CD19+ cancer cells through YSD, with antigen variations enabled by promoter choice and GPCR stimulation (**Fig. 2g-j**). Baseline constitutive expression levels of the seven chosen promoters yielded a 562-fold span of CD19 levels across the strain library, with the promoter P_TDH3_ reaching a CD19 level of 20.3±2.9% of NALM6 (**Fig. 2g**, **Suppl. Fig. 10**). For all strains, CD19 levels could be significantly diversified from their baseline through GPCR stimulation (Ste2) by ɑ-factor (0-100 µM) resulting in a multitude of different CD19+ profiles (p<0.0001)(**Fig. 2j**). Specifically, the highest CD19 levels were obtained for P_TDH3_ at 1,724±151-fold higher than the lowest basal CD19+ level (p≤0.0001)(**Fig. 2j**). However, P_MFA2_ showed the largest span in absolute CD19 levels upon stimulation (p≤0.0361) (**Suppl. Fig. 11**), by increasing 322.7±20.7-fold (**Fig. 2h, Suppl. Fig. 12**), to a level that was 2.7±0.4-fold higher than baseline P_TDH3_ expression, commonly regarded as the strongest constitutive yeast promoter^82^ (**Fig. 2j**). P_FUS1_, P_FIG1_, P_MFA1_, and P_MFA2_ could all provide drastic changes in CD19 levels relative to the unstimulated condition (p<0.0001) (**Fig. 2j**), with P_FUS1_ (774±165-fold) and P_FIG1_ (630±99-fold) providing up-regulation from undetectable CD19 to levels comparable to P_TDH3_, highlighting that these individual strains were capable of spanning the entire CD19 range of the unstimulated strain library (**Fig. 2g-j**). Surface display levels were also boosted by PRP activation, providing an estimated maximum PRP boost effect of 2.9±0.3-fold increase in CD19 (p<0.0001) (**Fig. 2i**), which was attributed to GPCR stimulation (SV=94.7%) and was equivalent across strain design differences (SV=0.9%) (**Suppl. Table 5**). Remarkably, the change in absolute output from the 3.1±0.3-fold up-regulation of P_TDH3_, resulting from the PRP boost, was greater than the 774±165-fold up-regulation of P_FUS1_ (150.2±37.5%, p=0.012) (**Suppl. Fig. 11**), hence the PRP boost allows high-intensity antigen presentation and larger diversity in the library.

In summary, CD19 levels were controllable on SCASA yeast cells, enabling CD19+ profiles spanning up to 1,724-fold in intensity. Importantly, the control of CD19 levels was I) continuous (non-binary), II) dependent on the GPCR ligand concentration, III) showing no residual non-displaying populations, IV) specific for the chosen processing module promoter, and V) relying on two separate effects, namely promoter regulation and a post-transcriptional PRP boost caused by phenotypic changes to the yeast cell (**Fig. 2**).

### Co-cultivation of yeast and human T cells

Before proceeding to yeast-mediated CAR T cell activation assays, we first assessed physiological properties of the co-cultivation of yeast with human immune cells. To confirm that undesired yeast growth would not occur, we first established that yeast showed negligible growth in T cell media, RPMI+10%FBS and ImmunoCult™-XF T Cell Expansion Medium, not initiating growth for at least 22 hrs, providing stable yeast cell numbers during co-cultivation (**Suppl. Fig. 13**). Secondly, to assess T cell viability and proliferation in the presence of yeast, we co-cultivated yeast with donor-derived naïve T cells at cellular ratios of 0.5x, 1.0x, and 10x yeast cells with compliance to T cell growth requirements for 96 hrs. Here we found that T cell viability and proliferation were not affected by the presence of yeast at any ratio (**Suppl. Fig. 13**). Lastly, we verified that the SCASA system of the engineered yeast cells remained functional during T cell growth conditions, despite the constrained growth in T cell media (**Suppl. Fig. 13**).

In summary, all experiments indicated that co-cultivation of yeast and T cells was possible under standard T cell growth requirements without affecting the viability or inducing proliferation in donor-derived naïve T cells, and with yeast cells showing constrained growth yet retaining SCASA system functionality.

### Controlled human immune cell activation by SCASA yeast cells

We proceeded to investigate the ability of SCASA yeast cell to confer communication with immune cells of relevance to immuno-oncological applications, specifically, CAR T cells. Here, a Jurkat cell line incorporating a luciferase reporter system linked to the T cell activation transcription factor NFAT (Jurkat NFAT-Luc) was transduced with lentiviral vectors to express the FDA-approved anti-CD19 CAR, FMC63-CD8ɑ-4-1BB-CD3ζ (4-1BB CAR) used in tisagenlecleucel^3,41,83^. Consequently, CAR T cells provide a bioluminescent signal of proportonal intensity to activation imposed by target CD19+ yeast cells (**Fig. 3**).

**Figure 3.**
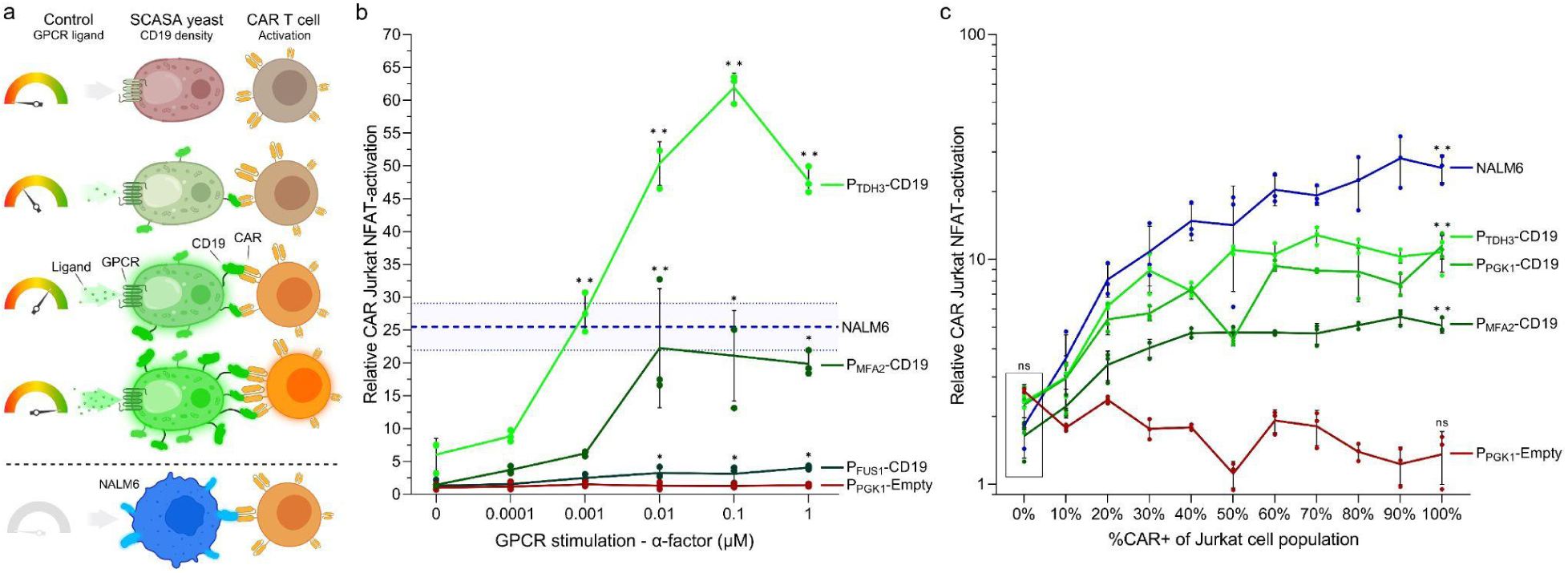
Activation of 4-1BB CAR Jurkat NFAT-Luc cells using SCASA yeast cells. **a.** Illustration of Jurkat cell co-culture with SCASA yeast cells. Jurkat cells that express anti-CD19 CARs (*CAR+*) bind to CD19+ SCASA yeast cells, analogously to as for the CD19+ NALM6 human cancer cell line. Here, Jurkat cells contain a NFAT-Luc reporter system, which upon activation expresses luciferase. SCASA yeast cells can be stimulated through their GPCRs to induce differential CD19 antigen density (e.g. Ste2 with ligand ɑ-factor). **b.** Relative CAR activation in co-cultures of 100% CAR+ Jurkat NFAT-Luc cells with SCASA yeast designs; P_FUS1_-CD19, P_MFA2_-CD19, and P_TDH3_-CD19 (*green lines*) at a target-to-effector (T/E) cell ratio of 0.2x for 18 hrs., with increasing SCASA yeast GPCR stimulation for CD19 display control (0-1 µM ɑ-factor). Negative control: P_PGK1_-Empty lacking the CD19 CDS in the display construct (*red line*). Positive benchmark control: NALM6 (*blue line*). One-way ANOVA with Dunnett’s multiple comparisons statistical test of NFAT-activation compared to 0 µM is shown for each strain. **c**. Relative CAR activation (log_10_-scale) in co-cultures of Jurkat NFAT-Luc cells with SCASA yeast designs; P_MFA2_-CD19, P_PGK1_-CD19, and P_TDH3_-CD19, as well as a negative control P_PGK1_-Empty, and NALM6 as benchmark control. The percentage of Jurkat NFAT-Luc cells in the co-culture expressing CARs was graded from 0-100% CAR+. Two-way ANOVAs are shown for a comparison of background activation at 0% CAR+ (Tukey’s multiple comparisons test) (*box*), and significant activation within the 0-100% CAR+ range for each SCASA yeast strain and NALM6 relative to 0% CAR+ (Dunnett’s multiple comparisons test). Data is based on means of three biological replicates (n=3) and standard deviations hereof. Significance levels: ✱: P≤0.05, ✱✱: P≤0.001. Not all pairwise comparisons are shown. All statistics and extended analyses in: **Suppl. Fig. 14** and **Suppl. Table 8.**

To demonstrate controllable synthetic cell-to-cell communication, we diversified the CD19 levels on target SCASA yeast cells by GPCR stimulation, to simulate cancer cells with varying CD19 antigen densities, and performed co-cultivations with CAR Jurkat NFAT-Luc cells at a low target-to-effector cell ratio of 0.2x (**Fig. 3a**). Upon GPCR stimulation, all CD19+ yeast designs significantly induced activation of 4-1BB CAR Jurkat cells (p≤0.0045), with graded NFAT responses dependent on both the GPCR stimulation level and processing module promoter (p<0.0001) (**Fig. 3b**). Hence, SCASA yeast activated CAR Jurkat cells in an antigen density-dependent manner. The most intense CAR Jurkat cell activation was enabled by the P_TDH3_-CD19 design yielding 61.9±2.2-fold increased NFAT activity relative to a control mono-culture (p<0.0001), which exceeded the activation elicited by NALM6 by 2.4±0.4-fold (p<0.0001) (**Fig. 3b**).

To confirm the specificity of SCASA yeast-induced CAR activation, we made co-cultivations with a constant number of Jurkat cells that varied in the proportion of CAR+ cells, spanning from 0-100% CAR+ (**Fig. 3c**). Yeast containing CD19 in the display design could all significantly activate CAR Jurkat cells (p<0.0001) (**Fig. 3b-c**), while a CD19^neg^ control yeast (P_PGK1_-Empty) never induced activation under any conditions of 0-100% CAR+ levels (**Fig. 3c**) or GPCR stimulation levels (**Fig. 3b**). Likewise, no activated Jurkat cells were detected in the range of 0-10% CAR+ for any co-cultivation, nevertheless, any >10% CAR+ increase provided significant responses for NALM6 and CD19+ yeast designs only (p≤0.0331). Hence, CAR Jurkat cell activation relied entirely on CD19 presentation by yeast cells and CAR expression in Jurkat cells (**Fig. 3b-c**). Given the lack of indications that yeast itself can activate T cells, at least via NFAT, these results support the idea of yeast cells as an orthogonal platform for assaying CAR T cell activation.

Each CD19+ yeast design provided significantly different activation dynamics in the 0-100% CAR+ range (p<0.0001) (**Fig. 3c**) and across GPCR stimulation (p<0.0001) (**Fig. 3b**), corresponding to the individual yeast CD19 levels (**Fig. 2j**). Responses showed that 4-1BB CAR-mediated NFAT activation was differentially sensitive to antigen-densities. Specifically, high basal CD19 levels provided higher dynamic range and sensitivity to GPCR stimulation (i.e. P_MFA2_-CD19 and P_TDH3_-CD19), indicating that antigen densities satisfied the effective activation threshold (**Fig. 3b, Suppl. Fig. 14**). Accordingly, insufficient absolute CD19 to reach threshold levels for effective activation explains why the ∼775-fold CD19 range of P_FUS1_-CD19 (**Fig. 2h**) was not reflected in the CAR Jurkat cell response at this target-to-effector ratio (**Fig. 3b**), which only increased by 3.1±0.2-fold (**Suppl. Fig. 14**).

In summary, CAR T cells (Jurkat NFAT-Luc) were activated by CD19-presenting SCASA yeast cells in a manner that was I) CD19-specific, II) CAR-specific, III) sensitive to CD19 antigen densities in regards to the intensity of NFAT activation, IV) different for each SCASA yeast cell design, V) controllable through GPCR stimulation of yeast cells, VI) capable of surpassing activation levels of NALM6, and VII) unaffected by the presence of yeast itself (**Fig. 3b-c**).

### SCASA yeast applications in characterizing CAR designs and downstream signaling

We next sought to demonstrate the direct application of CD19+ SCASA yeast cells in characterizing responses of different CAR designs, employing 50 distinct yeast-based stimulatory conditions of antigen densities and target-to-effector cell ratios. We chose to compare the two most commonly used FDA-approved designs, namely the 4-1BB CAR (FMC63-CD8ɑ-4-1BB-CD3ζ) and the CAR used in axicabtagene ciloleucel and brexucabtagene autoleucel, FMC63-CD28-CD28-CD3ζ^3,41^, referred to as the CD28 CAR. These CARs contain different hinge, transmembrane, and co-stimulatory domains, which has previously been shown to cause significantly different responses to antigen-density variations^25,35,41,44^. To quantify activation, we used a triple-parameter-reporter Jurkat T-cell line (TPR Jurkat cells) providing fluorescent output for three crucial T-cell activation transcription factors; NF-κB, NFAT, and AP-1, involved in proliferation, differentiation, and effector functions^41,84^ (**Fig. 4a**).

**Figure 4.**
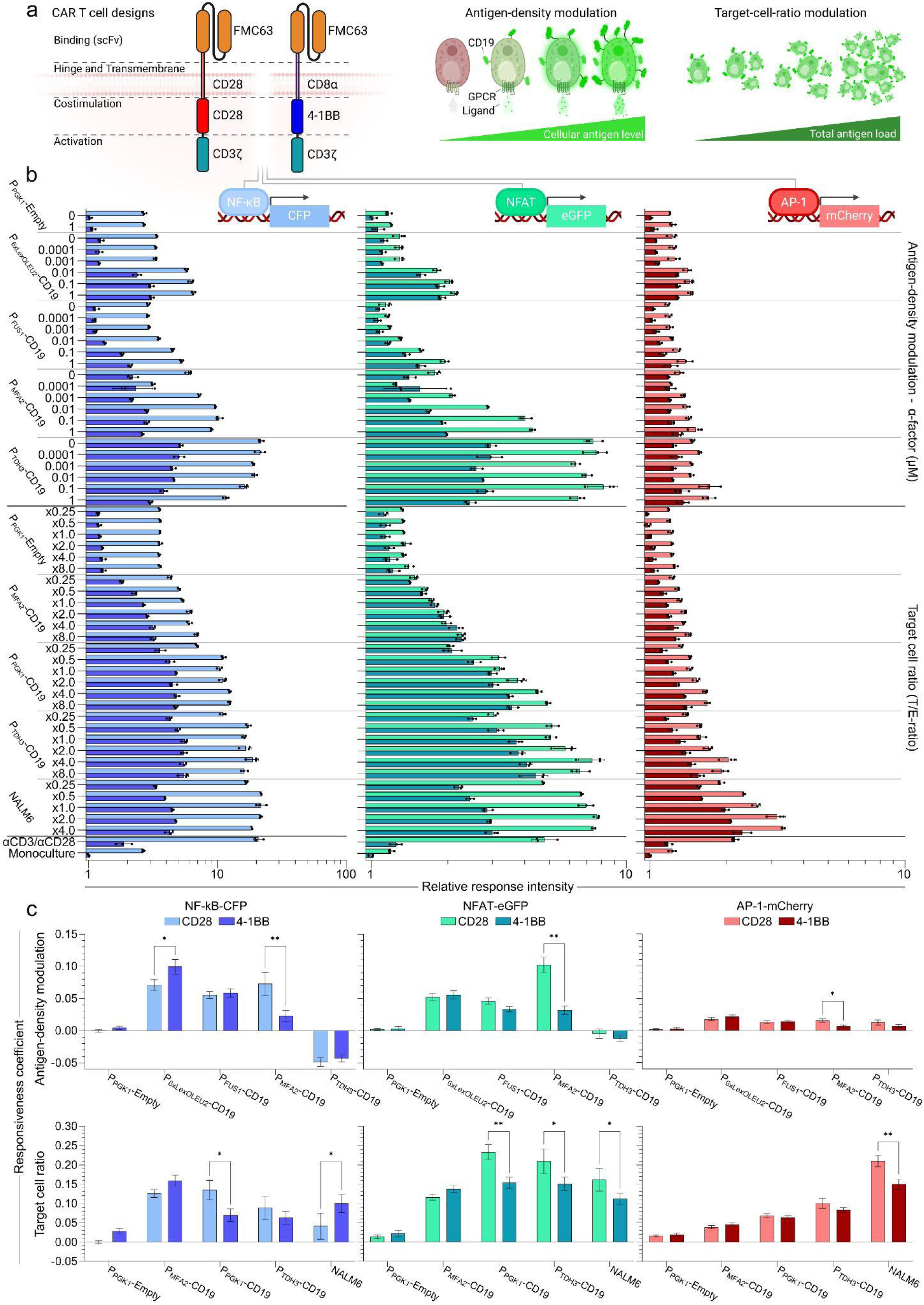
Characterization of FDA-approved CAR T cell designs and down-stream signaling using SCASA yeast cells. **a**. CAR T cells designs FMC63-CD28-CD28-CD3ζ (axicabtagene ciloleucel/brexucabtagene autoleucel) (‘CD28 CAR’) and FMC63-CD8ɑ-4-1BB-CD3ζ (tisagenlecleucel) (‘4–1BB CAR’), expressed in a triple-parameter-reporter (TPR) Jurkat T-cell line that couple T-cell activation transcription factors to fluorescent outputs; NF-κB-CFP, NFAT-eGFP, and AP-1-mCherry. CAR TPR Jurkats were co-cultivated for 24 hrs. with six CD19 SCASA yeast strains with different processing modules and hence different baseline CD19 levels (Fig. 2). Antigen-density modulation was examined by GPCR stimulation of individual strains (0-1 µM ɑ-factor) at a 1.0x target-to-effector (T/E) cell ratio. The effects of increasing target cells numbers were examined by modulating the T/E-ratio of SCASA yeast strains with fixed antigen densities and NALM6 from x0.25 to x8.0 relative to a fixed amount of CAR T cells. **b**. Relative response intensity of NF-κB-CFP (*blue*), NFAT-eGFP (*green*), and AP-1-mCherry (*red*) for the CD28 CAR (*light*) and 4-1BB CAR (*dark*) across all examined conditions, normalized to 4-1BB monocultures (log_10_-scale). ɑCD3/ɑCD28 dynabeads (1.0x) were employed as a positive control for reporter genes. **c**. Responsiveness coefficients for each reporter gene and CAR design, describing the degree of response change as a function of changes in antigen density and T/E-ratios for individual SCASA yeast strains and NALM6 - i.e. the response sensitivity to changes. Data represents means of median fluorescence intensities (mMFI) for three biological replicates (n=3) and standard deviations hereof. Selected comparisons from two-way ANOVAs with Šídák’s multiple comparisons tests are shown. Significance levels: ✱: P≤0.05, ✱✱: P≤0.001. All statistics and extended analyses in: **Suppl. Fig. 15-19** and **Suppl. Table 9-10**.

CAR signaling significantly increased in response to co-cultivation with CD19+ SCASA yeast of different antigen densities and target cell ratios for NF-κB (≤21.6±0.9-fold), NFAT (≤8.1±1.1-fold), and AP-1 (≤2.0±0.1-fold) (p<0.0001) (**Fig. 4b**). CD28 and 4-1BB CAR designs notably affected NF-κB and NFAT signaling intensities, whereas AP-1 was more consistent between the designs across all conditions (≤1.5±0.1-fold) (**Suppl. Fig. 15**). Specifically, the CD28 CAR caused higher baseline NF-κB activity than the 4-1BB CAR independently of CAR stimulation (2.6±0.1-fold, p<0.0001) and stronger NF-κB signaling upon CAR stimulation (≤5.6±0.1-fold) (**Suppl. Fig. 15**). NFAT signaling was affected differently by the CAR designs, as the CD28 and 4-1BB CAR had similar NFAT baselines, but upon antigen-density increase, NFAT signaling became preferentially activated by the CD28 CAR (≤2.9±0.4-fold, p<0.0001) (**Suppl. Fig. 15**).

Systematic variations in response dynamics revealed CAR-specific signaling (**Suppl. Fig. 16**) and distinct NF-κB, NFAT, and AP-1 signaling (**Suppl. Fig. 17**), which all were dependent on antigen densities (**Suppl. Fig. 15-17**). To quantify signaling dynamics, we performed regressions to obtain responsiveness coefficients that confirmed the differential sensitivity of responses to changes in antigen density and target cell ratios for both CARs (p<0.0001) (**Suppl Fig. 18**-**19**). The CD28 CAR responded with higher dynamicity than the 4-1BB CAR (≤1.72±0.03-fold, p<0.0001) (**Suppl. Fig. 15**) and had higher change in responsiveness to variations in high-antigen density targets (≤3.2±0.8-fold, p<0.0001) (**Fig. 4c**), exemplified by P_MFA2_-controlled CD19 levels (**Fig. 4c, Suppl. Fig. 15**). Conversely, the 4-1BB CAR showed a tendency for higher change in responsiveness against low-antigen targets, such as P_6xLexOLEU2_-CD19 (**Fig. 4c, Suppl. Fig. 15**). In parallel, NF-κB signaling displayed higher responsiveness for changes in low-antigen-density targets, such as P_6xLexOLEU2_-CD19 (p<0.0001) (**Suppl. Fig. 19**), most evident for the 4-1BB CAR, while NFAT displayed higher responsiveness for changes in high-antigen-density targets (e.g. P_MFA2_-CD19), most evident for the CD28 CAR (**Suppl. Fig. 19**). For the highest-antigen-density target, P_TDH3_-CD19, similar overstimulation was induced in both CARs by downregulation of NF-kB and NFAT (p<0.0001) (**Fig. 4c, Suppl. Fig. 15-17,19**). AP-1 activity retained upregulation (p<0.0001) (**Fig. 4c, Suppl. Fig. 15-17,19**), indicating that this signaling pathway was differentially regulated from those controlling NF-kB and NFAT. Moreover, AP-1 activity was more responsive to target variations in the CD28 CAR T cells (**Fig. 4c**), and relative to NF-κB and NFAT, more responsive to variations in target cell ratios than antigen-densities (**Fig. 4c**), and was preferably induced by NALM6 (p<0.0001) (**Suppl. Fig. 16,19**).

Lastly, of interest to SCASA yeast designs, both CAR designs could respond with equal or higher intensity (**Fig. 4b**), dynamic range (**Suppl. Fig. 16-17**), and responsiveness (**Suppl. Fig. 19**) to yeast compared to NALM6. Yeast itself (P_PGK1_-Empty) showed orthogonality to T cell activation, as it did not activate NFAT, NF-κB, or AP-1 for either the CD28- or 4-1BB-based CAR (**Fig. 4, Suppl. Fig. 15-19**). Notably, CD19+ SCASA yeast could cover a physiologically relevant range of CAR stimulation without increasing cell ratios, from the incipient responses to P_FUS1_-CD19, to overstimulation by increasing antigen density in P_TDH3_-CD19, for both NF-κB and NFAT (**Fig. 4b-c, Suppl. Fig. 17**). In particular, the P_MFA2_-CD19 SCASA strain alone provided high resolution and could significantly resolve the main response dynamics outlined above (**Fig. 4, Suppl. Fig. 15-19**), demonstrating the feasibility of a single-strain approach for general CAR design characterization. Additionally, the CARs showed similar response patterns to PRP-dependent P_FUS1_-CD19 and non-shmooing PRP-orthogonal P_6xLexoLEU2_-CD19 (**Fig. 4b-c, Suppl. Fig. 16-17,19** and **Suppl. Fig. 8**), confirming that responsiveness is a matter of antigen-density increase and not PRP activation.

In summary, using SCASA yeast cells, we determined that two CAR designs affect NF-κB, NFAT, and AP-1 activity in three different ways, with dynamics varying with CAR-stimulation intensity, of which antigen density and target cell ratios are individual parameters. The CD28 CAR was generally more responsive and exhibited stronger responses than the 4-1BB CAR, especially against high-antigen density targets, but both CAR types were sensitive to excessive stimulation leading to downregulated responses. Our conclusions regarding these CAR designs align with the current knowledge in the field^25,35,41,44,85^.

### Characterizing a donor-derived CAR T cell product using SCASA yeast cells

One important aspect of advancing cellular immunotherapies is the ability to robustly characterize responses of CAR T cell products resulting from novel designs or manufacturing methods. For this purpose, we proceeded to demonstrate the relevance of applying SCASA yeast cells by characterizing a CAR T cell product from a healthy donor made using a novel CRISPR-MAD7 method^86^ and based on the alternative Hu19-CD8ɑ-CD28-CD3ζ CAR that contains fully-human antigen-binding domains and has been associated with lower toxicity^87,88^.

Importantly, the SCASA yeast P_PGK1_-CD19 design activated the CAR T cells in a manner analogous to NALM6 across all examined target-to-effector cell ratios, hence verifying successful yeast-based simulation of CD19+ cancer cells towards the clinical Hu19-CD8ɑ-CD28-CD3ζ CAR T cell product (**Fig. 5a-c**). Specifically, CAR+ CD3+ T cells showed up-regulation of activation marker CD69 in co-cultures for all target cell ratios with P_PGK1_-CD19 yeast (p<0.0001) and NALM6 (p≤0.0001) (**Fig. 5a-b**). Meanwhile, no CAR-CD3+ T cells were activated for any target cell or condition, as also seen for the non-engineered control (CTRL) T cells (**Fig. 5a-b**). Furthermore, for the additional SCASA yeast cell designs P_MFA2_-CD19 and P_TDH3_-CD19, up-regulation of CD69 and CD25 was detected after 5 months of cryopreservation in liquid nitrogen (**Suppl. Fig. 20**).

**Figure 5.**
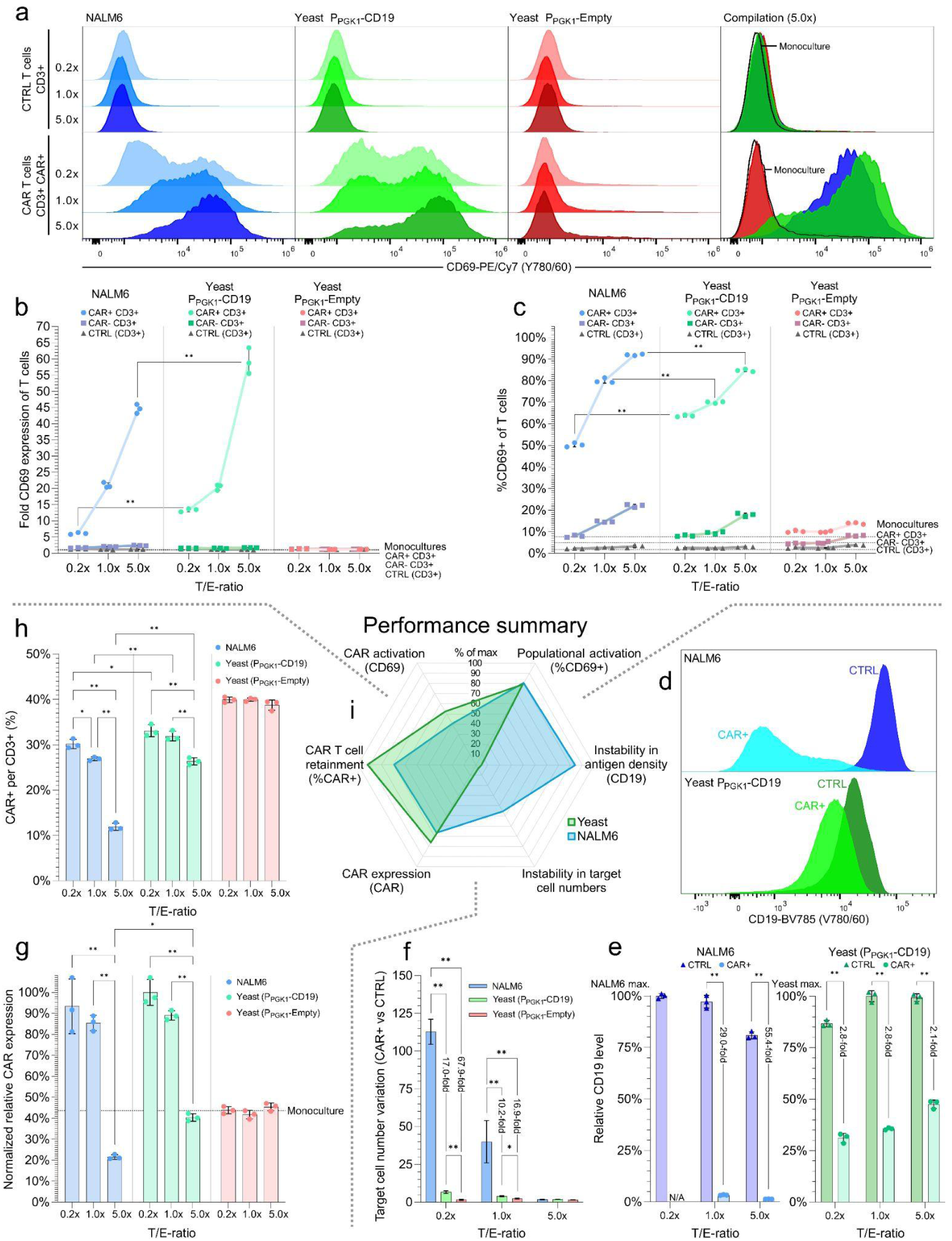
Activation of CAR T cells from a healthy donor using engineered SCASA yeast cells. **a.** CD69 expression in non-engineered control (CTRL) T cells (alive, CD3+) (*upper row*) and CAR T cells (alive, CD3+, CAR+) (*lower row*) after co-cultivation with NALM6 (*blue*), SCASA yeast cell P_PGK1_-CD19 (*green*), and negative control yeast P_PGK1_-Empty lacking CD19 (*red*), at three different target-to-effector (T/E) cell ratios (0.2x, 1.0x, 5.0x) for 20 hrs. An overlay of CD69 expression for 5.0x co-cultivations is shown (*far right*), including unstimulated T cell monocultures (*dotted line*). **b**. CD69 expression intensity of different T cell populations from co-cultivations: CAR+ (*circle*) and CAR- (*square*) T cells from the CAR-engineered population, CTRL T cells (*triangle*), and baseline CD69 of monocultures (*dotted lines*). Normalization: CTRL T cell monoculture. **c**. Percentage of T cell populations expressing CD69 (%CD69+) at any intensity for co-cultivations, and baseline %CD69+ of monocultures (*dotted lines*). **d**. CD19 levels on target cells after co-cultivations at 5.0x ratios for NALM6 (*blue*) and P_PGK1_-CD19 yeast (*green*) in CAR T cells (*light*) and CTRL T cells (*dark*). **e**. Relative CD19 levels to the measured maximum of NALM6 and P_PGK1_-CD19 yeast individually after co-cultivations with CTRL T cells (*triangle*) and CAR T cells (*circle*). **f.** Absolute fold target cell number variation per alive CD3+ T cell between CTRL and CAR T cell cultures for NALM6, P_PGK1_-CD19, and P_PGK1_-Empty, disregarding increased or diminished levels of target-to-effector cells, by acquiring the reciprocal value of fold changes<1. **g**. CAR expression intensity (CAR-DL488, B525/40) of the CAR+ population relative to the CAR-population (background), normalized to maximal CAR expression. **h**. Percentage of CAR+ T cells per CD3+ T cell detected after co-cultivations. **i**. Comparison of general performance of P_PGK1_-CD19 SCASA yeast (*green*) and NALM6 cancer cells (*blue*), quantified as percent intensity of most extreme behavior observed (% of max), averaging across T/E ratios. This plot summarizes all other plots (**a**-**h**), as indicated by dotted lines. Data represents means of cell counts or median fluorescence intensities (mMFI) for three biological replicates (n=3) and standard deviations hereof. Histograms are normalized to mode (CD69) or unit area (CD19) and are representative replicates. Selected comparisons from two-way ANOVAs with Tukey’s multiple comparisons tests are shown. Significance levels: ✱: P≤0.05, ✱✱: P≤0.001. All statistics and extended analyses in: **Suppl. Fig. 20-23** and **Suppl. Table 11**.

In summary, the CAR T cell product was functional towards CD19+ targets, as T cells activated solely upon CAR expression resulting from successful insertion via the CRISPR-MAD7 method. This was equally validated by the use of SCASA yeast cells or NALM6 cancer cells (**Fig. 5a-c**).

### SCASA yeast cells are efficient and robust CAR T cell activators

Having confirmed that CAR T cell products could be verified with SCASA yeast, we next sought to investigate the activation efficiency, i.e. target cell capacity to induce responses, and robustness, i.e. target cell maintenance of stimulatory conditions. Here, we found that increasing the ratio of SCASA yeast cells per CAR T cell intensified CAR activation, with significantly increased CD69 expression levels per CAR T cell, rising from 13.3±0.5-fold to 59.2±4.0-fold over the level of CTRL T cells (p<0.0001)(**Fig. 5b**), as well as enlargement of the population of activated CAR T cells (%CD69+) from 63.6±0.5% to 84.7±0.6% (p<0.0001)(**Fig. 5c**). A similar behavior was seen for the response to NALM6 at increasing ratios, however, with significant differences to SCASA yeast, as NALM6 provided lower or equal intensity of CAR activation, spanning from 6.1±0.3-fold to 44.6±1.4-fold CD69 expression (p<0.0001)(**Fig. 5b**), and had a larger spread in the number of activated CAR T cells, spanning from 50.3±0.9% to 91.9±0.3% (p<0.0001)(**Fig. 5c**). Overall, SCASA yeast and NALM6 co-cultivations shared the conclusion that CAR T cells are activated in a cell-to-cell ratio-dependent manner in relation to the amount of activated CAR T cells and their response intensity.

Interestingly, P_PGK1_-CD19 SCASA yeast cells could activate CAR T cells more intensely than NALM6 (**Fig. 5b**), yet had a CD19 level of merely 23.35±1.65% compared to that of NALM6 (**Suppl. Fig. 21**). This higher efficiency of SCASA yeast to activate the CAR T cells led us to investigate the target cells’ performance and robustness throughout co-cultivation. Here, we observed that NALM6 CD19 levels showed high instability throughout co-cultivation, evidenced by the occurrence of a CAR T cell-resistant CD19^low^ relapse-like phenotype of the NALM6 cancer cells during co-cultivation, as also observed by others^25,89–91^ (**Fig. 5d**). The appearance of the CD19^low^ NALM6 phenotype was fully dependent on CAR expression (SV=98.2%, p<0.0001) (**Suppl. Table 11**), and hence developed only in CAR T cell co-cultivations, while being undetectable in NALM6 monocultures and in co-cultivations with CTRL T cells (**Fig. 5d**). Specifically, NALM6 CD19 levels dropped 98.54±0.05% within 20 hrs of co-cultivation to a level that was 55.4±2.2-fold lower than in the CTRL co-cultivation (p<0.0001)(**Fig. 5e**), resulting in a large alive CD19^low^ population (19.8±0.9% of all cells detected) for the 5.0x co-cultivation with CAR T cells (**Fig.5d**, **Suppl. Fig. 22**). When equal amounts of NALM6 and CAR T cells were co-cultivated the effect was less drastic (29.0±1.9-fold lower CD19 levels) (**Fig. 5e**), resulting in killing of most NALM6 cells (0.5±0.1% of all cells detected) (**Suppl. Fig. 22**). When CAR T cells were in excess, all NALM6 cells were killed (**Suppl. Fig. 22**). Comparatively, SCASA yeast cells showed up to 26.5-fold more robust CD19 presentation than NALM6 throughout co-cultivations (**Fig. 5d-e**), with only 2.1±0.1-fold to 2.8±0.1-fold differences in CD19 levels between CTRL and CAR T cell co-cultivations (**Fig. 5e**). Additionally, CD19+ SCASA yeast cell numbers were up to 17.0-fold more robustly maintained during CAR T cell co-cultivations than observed for NALM6 cells (p<0.0001) (**Fig. 5f, Suppl. Fig. 23**), and there was no indication that SCASA yeast cells were killed by the CAR T cell cytotoxic response, contrary to NALM6 (**Suppl. Fig. 22**). Hence, the difference in CAR T cell activation patterns could be explained by a higher robustness in both antigen density and cell numbers of SCASA yeast cells compared to NALM6 (**Fig. 5d-f**), allowing yeast to be a more consistent target cell population to continuously provide activation signals to the CAR T cells, resulting in a higher efficiency of activation (**Fig. 5a-c**).

Further inspection of the CAR T cells showed that CAR expression patterns and the amount of CAR+ cells per CD3+ T cells alternated throughout co-cultivation in a CD19-dependent manner. CAR expression was elevated at low and equal target-to-effector cell ratios (p<0.0001), while target cell outnumbering of CAR T cells was associated with lower CAR expression intensities (p<0.0001)(**Fig. 5g**). This antigen-dependent up- and down-modulation of CAR surface levels has previously been shown to be dependent on CAR-stimulation levels, and reported to be caused by at least CAR expression regulation and CAR internalization^25,31,38,44,89,92^. The %CAR+ of CD3+ T cells dropped with an increase in target cell numbers (p<0.0001), indicating a loss of CAR+ T cells (**Fig. 5h**). Importantly, these effects were seen for both SCASA yeast cells and NALM6, but were in both cases more pronounced for NALM6 (**Fig. 5g-h**).

Orthogonality was confirmed by the CD19^neg^ yeast control (P_PGK1_-Empty), which did not activate CAR T cells or CTRL T cells under any circumstances, showing no increase in CD69 or CD25 (**Fig. 5a-b, Suppl. Fig. 20**), did not affect CAR expression intensity (**Fig. 5g**), or change the number %CAR+ cells per CD3+ cells (**Fig. 5h**), collectively showing that the activation of CAR T cells by SCASA yeast cells was specifically caused by the interaction between surface-displayed CD19 and successfully expressed CARs, and that yeast could not activate CD3+ T cells unspecifically, corroborating the CAR Jurkat observations (**Fig. 3-4**).

In summary, SCASA yeast cells provided an efficient method for activating and confirming functionality of a novel donor-derived CAR T cell product. SCASA yeast cells perform on par with cancer cells in terms of CAR T cell activation, yet with higher robustness and consistency in regards to antigen density and target cell numbers than cancer cells throughout co-cultivation (**Fig. 5i**).

## Discussion

Currently, CAR T cells are one of the most expensive types of therapy, leading to inequality in accessibility and rendering it prohibitively expensive for healthcare systems^93^. The CAR field is still emerging, and major challenges need to be mitigated to improve safety and efficacy, including mitigation of inhibitory immunosuppression, severe toxicity, CAR T cell trafficking, and target antigen escape^3,29,94^. Hence, to manifest the full potential of CAR T cells, it is essential to improve CAR product performance and develop safer and more cost-effective cellular designs, supported by innovative ways to enhance our understanding of the obstructive mechanisms.

Various experimental strategies have been employed to understand how antigen density and modulation impact CAR T cell responses, each with distinct advantages and limitations. Typically, these employ target cell panels with differential antigen densities enabled by either exploiting default antigen variation in cell lines^37,42^, engineering antigen-gene copy-numbers^25,30–33^, differential target-antigen mRNA electroporation^36,43^, or modular cell platforms for transiently attaching purified antigens^44^. These approaches have the great advantage of close imitation of *in vivo* cellular interactions. However, mammalian cell lines are not orthogonal and adapt to CAR T cell exposure. This is exemplified by their cytotoxic sensitivity resulting in target cell loss biased by antigen density or defensive responses to this selective pressure, such as rapid and extensive antigen modulation^17,25,89–91^. This leads to inconsistency in primary parameters of CAR-responses, antigen density and target cell numbers, even down to the hourly timescale in both *in vitro* and *in vivo* experimental models^25,89–91^, as seen for NALM6 (**Fig. 5d-f, Suppl. Fig. 22-23**). In addition, mammalian cell lines can require laborious multiplex engineering and have risks of genetic drift^44^. For these reasons, robust non-cellular material-based systems, such as plate-bound antigen, have been employed to ensure precise antigen control and to orthogonalize the activating molecular interactions from other cellular interactions and confounding effects^31,38,45–49^. Similarly, yeast cells are inert to these distorting effects and remain robust signal providers upon CAR T cell exposure (**Fig. 3,4,5, Suppl. Fig. 13,14,22,23**). Conversely, orthogonality is arguably also a limitation, as non-cellular systems and SCASA yeast do not simulate the full spectrum of cellular interactions between cancer and immune cells, which could explain the differences observed for AP-1 activity (**Fig. 4**). CAR-based T-cell activation depends on the physical dimensions of the intercellular space and supramolecular immunological synapse structures, favoring kinase activity and excluding phosphatases^49,95,96^. However, CAR-synapses are more disorganized than conventional T-cell synapses, differ in employment of co-stimulatory, adhesion, and accessory receptor molecules, and are additionally affected by CAR affinity and target cell antigen density^38,39,49,95,96^. SCASA yeast relies on the ability of antigen-density model systems to be greatly simplified, analagously to non-cellular platforms, as CAR T cells do not exploit accessory receptors as conventional T cells^38,44,49^. For example, while LFA-1-binding ICAM-1 and CD2-binding CD58 have been shown to greatly increase TCR sensitivity against non-cellular systems, the effect is only modest on CAR T cells^38,44^. Accordingly, yeast-based systems displaying pMHC-II for TCR activation were initially less successful^60^, but has since been functionalized by co-display of the ICAM-1^56^. This indicates that while oversimplification is a risk of orthogonality, yeast inherently provides a solution by being a flexible platform for bottom-up engineering of complexity to approximate native interactions, while retaining control^55,56^.

Hence, we consider yeast to be a cost-effective, robust, and powerful platform for investigating the dynamics of specific molecular interactions with the ability to combinatorially modify and control signal molecules in a high-throughput manner. Mammalian cell lines are arguably better suited for more holistic studies of the complex interplay between CAR T cells and cancer cells, such as in cytotoxicity assays^25,35^.

Lastly, absolute orthogonality could potentially be difficult to obtain when employing yeast, as T cells express pattern recognition receptors (PRRs) that can bind pathogen-associated molecular patterns (PAMPs), such as Toll-like receptor 2 (TLR2) and CD5 recognition of yeast cell wall-derived zymosan, which upon stimulation can modify T cell responses^97–100^. Interestingly, yeast-induced PRR stimulation of T cells might induce relevant phenotypes to cancer immunotherapy, with for example studies showing that TLR2 stimulation can improve T cell anti-cancer phenotypes^101^, lower antigen density thresholds for T cell activation^102^, and improve CAR T cell responses^103^. The possible impact on synapse formation, lack of accessory molecules, physical dimensions, and potential T cell PRR-engagement by yeast PAMPs on CAR T cell activation are considerable uncertainties to be addressed in future investigations. Nevertheless, we did not encounter any hindrances of using yeast during co-cultivation or activation of T cells.

We have demonstrated that SCASA yeast can effectively simulate CD19+ cancer cells with precise on-demand antigen densities that control the activational state of CAR T cells, allowing for high-resolution interrogation of CAR T cell responses (**Fig. 3-5**). Specifically, we employed yeast to characterize CAR sensitivity to changes in antigen density and cellular ratios (**Fig. 3-5**), to verify that the response of the CD28 CAR is more intense than the 4-1BB CAR, particularly with high-antigen density targets which indicate patient-specific performance, and that these CAR designs differentially affect NFAT, NF-κB, and AP-1 activities (**Fig. 4**), and, finally, we validated functionality of a CAR T cell product made from donor blood using a novel CRISPR-MAD7 method and an alternative CAR design (**Fig. 5**).

The SCASA platform has potential for further advancement. Exchangeable GPCRs can be exploited to link responses to endogenous ligands, to ensure ligand orthogonality to target cells, for biomarker sensing, or to build cell-cell feedback-regulated systems^1,52,67^. The system is compatible with multiplexing through additional effector cassettes, and could allow incorporation of additional layers of orthogonal signaling inputs^75^. Similarly, co-display allows for investigation of multiple signals and closer imitation of immuno-oncological cellular interactions^38,44,56^. Furthermore, the fully DNA-encoded SCASA system eliminates the need for attaching purified proteins and results in a genotype-phenotype bond compatible with evolution platforms and screening of variant libraries^52,61,62^.

With the demonstrated potential and engineerability of yeast platforms, as well as developments and trends in yeast synthetic biology, we anticipate the advancement of yeast-based systems toward targeting immunological challenges and providing novel applications in the future.

## Methods

### Molecular cloning for yeast engineering

#### PCR and DNA handling

Genomic DNA (gDNA) was purified using the Yeast DNA Extraction Kit (Thermo Scientific, Cat.#78870). Custom DNA synthesis was done using gBlocks™ Gene Fragment synthesis service (Integrated DNA Technologies). For amplification of DNA fragments for cloning or genome integration, and inverse PCR plasmid construction, Phusion High-Fidelity (HF) PCR Master Mix with HF Buffer (Thermo Scientific, Cat.#F531L) was used, and, specifically, for amplification of Uracil-Specific Excision Reagent (USER) cloning DNA fragments, Phusion U Hot Start PCR Master Mix (Thermo Scientific, Cat.#F533L) was used with primers with USER-compatible uracil-containing tails. Standard 50 µL PCR reactions were composed of; 25 µL PCR Master Mix (2X), 2 µL of each primer (10 µM), DNA template scaled to 100-200 ng gDNA or 10-20 ng plasmid DNA, and Milli-Q H_2_O to a total reaction volume of 50 µL. For diagnostic and genotyping PCRs, such as colony PCR of *Escherichia coli* or yeast, One*Taq*® Quick-Load® 2X Master Mix with Standard Buffer (New England BioLabs, Cat.#M0486) was used. For each *E. coli* colony PCR, a 10 µL PCR mix was made; a small amount of cell mass was added to a mix of 5 µL PCR Master Mix (2X), 1 µL of each primer (10 µM), and 3 µL Milli-Q H_2_O. For each yeast colony PCR, a 20 µL PCR mix was made; cell mass was added until visible turbidity in a mix of 1 μL of each primer (10 μM) and 8 μL Milli-Q H_2_O, followed by cell lysis by heating at 98°C for 15 min., and after cooling, 10 µL PCR Master Mix (2X) was added. PCRs were conducted according to the manufacturer’s protocols using an S1000 Thermal Cycler (Bio-Rad). All primers were synthesized using a Custom DNA Oligos service (Integrated DNA Technologies) and can be found in **Suppl. Table 12**. Gel electrophoresis was done in 1%w/v agarose gels of 1X Tris-acetate-EDTA buffer at 90v for 30 min., using TriTrack DNA 6X Loading Dye (Thermo Scientific, Cat.#R1161), GeneRuler 1 kb DNA Ladder (Thermo Scientific, Cat.#SM0311), and RedSafe™ Nucleic Acid Staining Solution (iNtRON Biotechnology, Cat.#21141). Gel imaging was conducted on a Gel Doc XR+ System (Bio-Rad). PCR amplicons were purified via gel purification or column purification using the NucleoSpin Gel and PCR Clean-up kit (Macherey-Nagel, Cat.#740609.50). Amplicon sequences were verified by Sanger sequencing using the Overnight Mix2Seq Kit service (Eurofins Genomics). Primers, DNA fragments, amplicons, plasmids, and gDNA were always dissolved in Milli-Q H_2_O and stored at -20°C. An overview of all DNA parts, backbones, and plasmids can be found in **Suppl. Table 13-14.**

#### Construction of plasmids

Plasmid vectors containing genome-integration expression cassettes were based on the EasyClone-MarkerFree system compatible with CRISPR/Cas9 engineering ^104^, and were constructed with USER cloning and assembly techniques ^105^, however, modified by using custom standardized USER-compatible primer tails for DNA parts and inverse PCR for conversion of USER-compatible backbones, denoted MAD-cloning (**Suppl. Fig. 24**). Linearized USER-ready MAD-cloning backbones were generated by purification of inverse PCR-amplified EasyClone-MarkerFree integrative vectors using primers with USER-compatible tails. Similarly, custom yeast expression plasmids were constructed by converting template expression plasmids ^106,107^ into MAD-cloning compatible backbones. Employed parts, such as promoters, terminators, protein-coding sequences (CDSs), or entire genes, were amplified from yeast gDNA, plasmids, or ordered as gBlocks™, and in some cases modified by in-frame fusion using USER-cloning, and coding sequences were modified to contain the AAAACA Kozak sequence. Each cassette assembled by MAD-cloning contained standardized interchangeable parts: I) a promoter, II) a CDS, III) a terminator, IV) a linearized backbone, which for genome integration contained homology arms directed at EasyClone-MarkerFree genome integration sites flanked by *Not*I-sites. USER-assembly reactions contained 20-40 ng linearized USER-ready backbone, 40-80 ng of each USER-compatible part, 0.5 µL USER® Enzyme (New England BioLabs, Cat.#M5505), 1 µL 5X Phusion HF Buffer (Thermo Scientific, Cat.#F518L), and Milli-Q H_2_O to a total reaction volume of 5 μL. USER-assembly mixes were incubated at 37°C for 30 min. and transformed into *E. coli*. Plasmids for integration of GPCR expression cassettes were assembled as previously described ^63^. For CRISPR/Cas9 engineering, all gRNA was expressed from cassettes using the snoRNA *SNR52* promoter and a *SUP4* terminator ^108^. Custom yeast gRNA-expression plasmids were generated by inverse PCR of template gRNA-expression plasmids using a 5’-phosphorylated primer and a primer with a tail for replacing the 20 bp gRNA target sequence. Purified amplicons were then blunt-end ligated in a reaction containing: 1 μL T4 DNA Ligase (Thermo Scientific, Cat.#EL0011), 2 μL 10X T4 DNA Ligase Buffer (Thermo Scientific, Cat.#EL0011), and 17 µL amplicon dissolved in Milli-Q H_2_O, which was incubated for 2 hrs at room temp. Template plasmid was then removed by addition of 1 μL DpnI (New England BioLabs, Cat.#R0176), 5 μL 10X FastDigest Buffer (Thermo Scientific, Cat.#B64), and 24 μL Milli-Q H_2_O, which all was incubated for 1 hr at 37°C, and finally transformed into *E. coli*. All *E. coli* transformants for plasmid assembly were genotyped by colony PCR to screen for successful assembly, and correct assembly was verified by Sanger sequencing of purified plasmids using the Overnight Mix2Seq Kit service (Eurofins Genomics). Plasmids were purified from *E. coli* cultures using the NucleoSpin Plasmid kit (Macherey-Nagel, Cat.#740588.50). All plasmids contained ampicillin resistance (*ampR*) for propagation in *E. coli*, and retention in yeast relied on auxotrophic markers (*HIS3*, *URA3*, *LEU2*, *TRP1*) or nourseothricin antibiotic resistance (*NatR*). Primers, DNA fragments, amplicons, plasmids, and gDNA were always dissolved in Milli-Q H_2_O and stored at -20°C. An overview of parts, backbones, constructed plasmids, and plasmids originating from other studies ^63–65,68,69,79,80,104,106,107^ can be found in **Suppl. Table 13-14**, and primers in **Suppl. Table 12**.

#### Repair and Intentional Primer Dimer (IPD) PCR templates for gene knockout or alteration

For direct genome engineering, such as gene knockouts or minor alterations of genome sequences, DNA repair templates were co-transformed with gRNA-plasmids into Cas9-expressing yeast strains. Repair templates were either two genome-amplified sequences with overlapping homology defined by primer tails or an Intentional Primer Dimer (IPD) PCR amplicon with flanking homology to the cut site. IPD-PCR templates were generated by annealing and PCR-amplifying two complementary primers using each other as templates at their predicted annealing temperatures (50 µL Phusion HF, with 4 µL of each primer at 10 µM). IPD-PCR templates had genome-targeting homology tails and contained a complementary binding region for their amplification, which was also composed of one of several genome-unique CRISPR landing pads previously characterized ^109^, allowing for easy reengineering of the site (**Suppl. Fig. 25**).

#### Bacterial handling for plasmid construction and propagation

Plasmid propagation and USER-assembly were conducted by heat-shock transformation into *E. coli* strain DH5ɑ, which was cultivated in Terrific Broth or Luria-Bertani with ampicillin (100 mg/L) at 37°C in liquid media at 300 r.p.m. or on agar plates for 16-20 hrs. Before transformation, *E. coli* DH5ɑ was made chemically competent and stored at -80°C until needed ^110^. All *E. coli* strains were stored as cryostocks in media with 25% (v/v) glycerol at -80°C.

### Yeast strains, media, and engineering

#### Yeast handling

All yeast strains were engineered from *S. cerevisiae* strain BY4741 (**Suppl. Table 15**). Strains were grown in YPD media with 2% (w/v) glucose except for during experiments or for plasmid retention, in which strains were grown in Synthetic Complete (SC) media with 2% (w/v) glucose using appropriate amino acid dropout mixes (Sigma-Aldrich) and/or with the addition of antibiotic nourseothricin (100 mg/L) (Jena Bioscience, Cat.#AB-101). For experiments requiring pH-control, a SC medium with ammonium sulfate (AS) and urea (SC-AS/Urea) was buffered using citrate-phosphate buffer (CPB) ^111^. Yeast strains were generally grown at 30°C at 250 r.p.m. or on 2% (w/v) agar plates, unless otherwise stated for specific experiments. All strains were stored as cryostocks in media with 25% (v/v) glycerol at -80°C.

#### Yeast transformation and engineering

All transformations of yeast were done using the LiAc/ssDNA/PEG method ^112^. Plasmid-based expression utilized either 2µ- or CEN/ARS-type plasmids. All genome engineering relied on CRISPR/Cas9-based methods, by having cells constitutively expressing Cas9 from pEDJ391 ^106^. Genome integrations relied on co-transformation of I) FastDigest™ *Not*I-linearized (Thermo Fisher, Cat.#FD0593) cassette-containing USER-assembled plasmids with homology arms and II) gRNA helper vectors, both targeting the characterized integration sites described for the EasyClone MarkerFree system ^104^. Gene knockouts relied on co-transformation of I) a gRNA expression vector targeting the site of interest and II) an IPD-PCR repair template with homology flanking the Cas9 cut site, which contained a genome-unique CRISPR landing pad for enabling reengineering of the site ^109^. Similarly, scarless gene alterations were done by using an IPD-PCR template containing the novel sequence of interest. Genomic engineering was confirmed by genotyping via colony PCR and Sanger sequencing of cassettes, as described above.

From the BY4741 background, the strain DIX14 was optimized for GPCR biosensing by overexpression of native ɑ-factor pheromone-sensing mating GPCR Ste2 (*P_CCW12_-STE2-T_CYC1_*), and by balanced expression of native G protein G_ɑ_-subunit (*P_PGK1_-GPA1-T_CYC1_*), as well as by deletion of native PRP-coupled GPCRs (*ste2Δ0* and *ste3Δ0*), native G_ɑ_-subunit (*gpa1Δ0*), a negative feedback regulator (*sst2Δ0*), an ɑ-factor protease (*bar1Δ0*), and a cyclin-dependent kinase inhibitor that induce G1 cell cycle arrest (*far1Δ0*). Furthermore, native *AGA2* was deleted in preparation for YSD optimization (*aga2Δ0*). From DIX14, DIX17-28 were created by integration of yEGFP-expression cassettes. DIX34 was created by knocking out the PRP transcription factor Ste12 (*ste12Δ0*) and integrating a LexA-based synthetic transcription factor, as well as a yEGFP cassette with a LexO-containing promoter for orthogonalized signaling. The strain DIX41 was optimized for YSD by Aga1 overexpression (*P_TDH3_-AGA1-T_ADH1_*). DIX44-56,74 strains were generated from DIX41 for CD19 YSD experiments and co-cultivations, by integration of CD19-expression cassettes. Based on DIX45, genome alterations were made to replace the Gpa1 G_ɑ_-subunit with G_αz_- and G_αi2_-chimeras, to generate DIX57 and DIX58, respectively, for allowing coupling of heterologous GPCRs. From these strains, heterologous GPCRs were integrated to make the GPCR strain library (DIX59-DIX65). The background strain for orthogonal GPCR signaling (DIX33) was equipped with Aga1 overexpression and a CD19 cassette with the LexO-containing promoter for orthogonal GPCR-based control of CD19-display (DIX67). An overview of strains can be found in **Suppl. Table 15**.

### Human cell engineering and cultivation

#### Construction of anti-CD19 CAR Jurkat NFAT-Luc and CAR TPR Jurkat cell lines

To investigate the activation of CAR T cells, a Jurkat cell line equipped with a nuclear factor of activated T cell (NFAT) transcription factor luciferase reporter system (Jurkat NFAT-Luc) (Nordic BioSite, Cat.#BPS-60621) was further modified by insertion of an anti-CD19 CAR (FMC63-CD8ɑ-4-1BB-CD3ζ) ^83^, hence providing a bioluminescent luciferin signal of an intensity proportional to the level of T cell activation. Likewise, a triple parameter reporter Jurkat T cell line (TPR Jurkat cells) was employed to characterize CAR-induced activation of downstream transcription factor activity by fluorescent reporter genes, namely; NF-κB-CFP, NFAT-eGFP, and AP-1-mCherry^84^. Two different CAR designs were inserted into the TPR Jurkat cell line; FMC63-CD8ɑ-4-1BB-CD3ζ (referred to as the ‘4–1BB CAR’) and FMC63-CD28-CD28-CD3ζ (referred to as the ‘CD28 CAR’). Lentiviral particles containing the CD19 CAR elements were produced by lipofectamine-based co-transfection of HEK293 cells (ATCC, 293T Cat.#CRL-3216) with 3^rd^ generation packaging plasmids pMD2.G (Addgene, #12259), pMDLg/pRRE (Addgene, #12251), pRSV-Rev (Addgene, #12253), and the CAR transfer vector (pDTU), a modified version of pLenti-puro (Addgene, #39481), optimized to include a cPPT-CTS sequence, the EF-1ɑ promoter, and a Woodchuck Hepatitis Virus (WHV) Post-transcriptional Regulatory Element (WPRE). 24 hrs after transfection, lentiviral particles were harvested and concentrated using Lenti-X concentrator (Takara Bio) and stored at -80°C. Jurkat cell lines were transduced with lentivirus at an MOI of 5. After transduction, anti-CD19 CAR+ Jurkat NFAT-Luc and TPR Jurkat cells were single-cell sorted and expanded to adequate numbers before cryopreservation in liquid nitrogen. Jurkat cell lines were cultured in RPMI 1640 medium (ATCC modification) (Thermo Fisher, Cat.#A1049101) with 10% fetal bovine serum and 1%pen-strep (RPMI+10%FBS+1%PS) at 37°C with 5% CO_2_.

#### NALM6 cancer cell line

The human CD19+ NALM6 B cell precursor leukemia cell line (DSMZ, no.: ACC 128) was included as a benchmark positive control. NALM6 was cultured in RPMI+10%FBS+1%PS at 37°C with 5% CO_2_. The cell line was further engineered by genomic insertion of a CAG-promoter driven GFP-cassette in safe-harbor site AAVS1_3 using CRISPR-MAD7, as previously described ^86,113^.

#### Isolation of T cells from peripheral blood and CRISPR-MAD7 engineering for CAR insertion

This study was carried out in accordance with the Declaration of Helsinki. Human peripheral blood was obtained from healthy adults after obtaining informed consent (Technical University of Denmark - Rigshospitalet National Hospital approval BC-40). No personal information was gathered, and donors were anonymized for this study. All T cells were derived from the blood of healthy donors collected at the central blood bank at Rigshospitalet (Copenhagen, Denmark). Peripheral blood mononuclear cells (PBMCs) were isolated from buffy coat, by blood filtration using a Falcon 70 µM Cell Strainer (Corning, Cat.#352350), followed by 2X dilution in Dulbecco’s Phosphate Buffered Saline with 2% Fetal Bovine Serum (Stemcell Technologies, Cat.#07905), and finally, density centrifugation using SepMate™-50 (IVD) tubes (Stemcell Technologies, Cat.#85460) and Lymphoprep™ (Stemcell Technologies, Cat.#07811), according to the manufacturer’s protocols. Pan T cells were isolated from PBMCs by negative selection, using the EasySep™ Human T Cell Isolation Kit (Stemcell Technologies, Cat.#17951), with EasySep™ Buffer (Cat.#20144) and EasySep™ Magnet, The "big Easy" magnet (Stemcell Technologies, Cat.#18001), according to manufacturer’s protocols. T cells were then resuspended at 1×10^6^ cells/mL in ImmunoCult™-XF T Cell Expansion Medium (Stemcell Technologies, Cat.#10981) supplemented with Human Recombinant IL-2 (12.5 ng/mL) (Stemcell Technologies, Cat.#78036), IL-7 (5 ng/mL) (Stemcell Technologies, Cat.#78053), IL-15 (5 ng/mL) (Stemcell Technologies, Cat.#78031), and ImmunoCult™ Human CD3/CD28/CD2 T Cell Activator (25 µL/mL) (Stemcell Technologies, Cat.#10990) and cultured for 1.5 days at 37°C with 5% CO_2_. After activation, T cells were engineered using CRISPR-MAD7 according to a recently published method,^86^ to insert the CAR Hu19-CD8ɑ-CD28-CD3ζ (Hu19-CD828Z), containing an anti-CD19 fully-human scFv (Hu19), CD8ɑ hinge and transmembrane domains, a CD28 costimulatory domain, and a CD3ζ activation domain^88^ (Clinical trial: NCT02659943). In addition, the CAR contained a myc-tag for the detection of surface expression. The CAR was expressed using the EF-1ɑ promoter and a bovine growth hormone polyadenylation (bgh-PolyA) signal from the AAVS1_3 safe-harbor site. The CRISPR-MAD7 AAVS1_3 crRNA ribonucleoprotein formulation, Hu19-CD8ɑ-CD28-CD3ζ homology-directed repair (HDR) template generation, and transfection of donor-derived (primary) T cells using a Lonza 4D-Nucleofector with shuttle unit with use of M3814 (Selleckchem) recovery followed the previously described methods ^86,113^. Along with the CAR T cells, a control (CTRL) T cell product was established from the same T cell isolate, undergoing the same transfection protocol as the CAR T cells, however, without using a CAR HDR template. The CAR T cell product was cultivated in RPMI+10%FBS+1%PS at 37°C with 5% CO_2_ for experiments and cryopreserved as 10×10^6^ cells/mL in 1 mL 90% heat-inactivated Human AB Serum (Sigma-Aldrich, Cat.#H4522) +10% DMSO in liquid nitrogen. T cells for investigation of viability during co-culture with yeast were obtained from PBMCs isolated from healthy donor buffy coats, by density centrifugation using LymphoPrep™ Solution (Axis-Shield PoC, Cat.#1858) and cryopreserved at −150°C in FBS+10% DMSO. T cells were isolated from PBMCs by negative selection, using the EasySep™ Human T Cell Isolation Kit (Stemcell Technologies, Cat.#17951).

### Experimental procedures

#### Flow cytometry

Flow cytometric measurements were carried out using either a NovoCyte Quanteon 4025 Flow Cytometer System with a NovoSampler Q System (Agilent) or a CytoFLEX-S (V-B-Y-R) instrument (Beckman Coulter), according to the manufacturers’ protocols. Specifications are provided in the description of individual experiments below. Gating and compensation were done using FlowJo™ v10.8.1 Software (BD Life Sciences). Gating strategies employed fluorescence minus one (FMO) controls, as well as the employment of negative controls, where possible, for determining true positive signals (e.g. yEGFP+, CAR+) (**Suppl. Fig. 27-28**).

#### Processing module promoter characterization

For processing module promoter characterization, a transcriptome analysis was conducted (**Suppl. Fig. 4, Suppl. Table 1**), as described previously ^63^, from which candidate promoters were chosen. Each promoter was defined as the 1,000 bp sequence upstream of the gene CDS, and USER-assembled with yEGFP and T_CPS1_, to form reporter cassettes. Dose-response and phenotype features of strains containing the selected promoters, P_MFA2_, P_FUS1_, P_AGA2_, P_FIG1_, P_MFA1_, P_YCL076W_, P_HPF1_, P_IME4_, P_CSS1_, P_TDH3_, P_PGK1_ and P_6xLexOLEU2_, were then investigated upon GPCR stimulation via yEGFP expression (DIX17-28 and DIX34) (**Fig. 2, Suppl. Fig. 5-9**). The yeast strains were inoculated and pre-cultivated in 0.5 mL SC (24 hrs., 30°C, 250 r.p.m.), then diluted 10-fold and further cultivated in SC (24 hrs., 30°C, 250 r.p.m.), whereafter, for each replicate, cultures were diluted 50-fold by transfer to 200 µL volumes comprising a 10-fold dilution series of ɑ-factor (0-100 µM) (peptide-seq.: WHWLQLKPGQPMY) (GenScript) in SC+1%DMSO, and then stimulated for 5 hrs. (30°C, 250 r.p.m.) (**Suppl. Fig. 26**). Flow cytometric measurement was done using a NovoCyte Quanteon flow cytometer (Agilent), with cells suspended in 1X phosphate-buffered saline (PBS), and collection of 20,000 cells per replicate to quantify yEGFP levels (B525/45-A) and PRP-regulated morphological changes (shmooing) (SSC-A). The gating strategy is described in **Suppl. Fig. 27**. To approximate the effects of PRP activation on expression not directly related to transcriptional upregulation, SSC-normalization was performed by dividing individual fluorescence intensities by the corresponding SSC light scatter intensity. Each condition was examined in three biological replicates. A two-way ANOVA with Tukey’s or Dunnett’s multiple comparisons test was conducted for individual conditions, and to fulfill the assumptions of parametric tests, some data was log_10_-transformed, all specified in **Suppl. Table 2-3**. Linear and non-linear variable slope (four parameter) regressions can be found in **Suppl. Table 3-4**.

#### CD19 display characterization

To investigate the different variants of CD19 modules (**Fig. 2, Suppl. Fig. 3,10-12**), yeast cells were grown and stimulated to present CD19 differentially, whereafter they were stained with antibodies and assessed by flow cytometry. For the investigation of the strain library with constitutive and dynamically regulated expression of CD19 (No YSD, P_PGK1_-Empty, P_MFA2_-CD19, P_FUS1_-CD19, P_FIG1_-CD19, P_MFA1_-CD19, P_HPF1_-CD19, P_TDH3_-CD19, P_PGK1_-CD19), DIX41-56 strains were inoculated and pre-cultivated in 0.5 mL SC (24 hrs., 30°C, 250 r.p.m.), then diluted 10-fold and further cultivated in SC (24 hrs., 30°C, 250 r.p.m.), whereafter, for each replicate, 50,000 cells were transferred to 300 µL volumes comprising a 10-fold dilution series of ɑ-factor (0-100 µM) (GenScript) dissolved in SC+1%DMSO, and stimulated for 20 hrs. (30°C, 250 r.p.m.). For conventional YSD using strain EBY100 and pCT-plasmid-based expression ^107^ (**Suppl. Fig. 1**), strains were pre-cultivated in SC -trp, then transferred to 5 mL SC -trp +2%Gal for induction of YSD or +2%Gluc as a negative control (24 hrs., 30°C, 250 r.p.m.). For the investigation of P_GAL1_-titratability of plasmid-based (DIX41+pMAD58) and genome-integrated CD19 YSD (DIX74) (**Suppl. Fig. 2**), strains were pre-cultivated in SC -ura and SC, respectively, then transferred to SC -ura and SC, respectively, with varying concentrations of galactose (0.0002%-2%Gal), using raffinose as compensating carbon source, and YSD was induced for 24 hrs (30°C, 250 r.p.m.). Strains were similarly grown in 2%Gluc as negative controls. For assessment of the GPCR library for inducing CD19 YSD using different types of ligands (**Fig. 2, Suppl. Fig. 3**), strains were pre-cultivated in 0.5 mL SC (24 hrs., 30°C, 250 r.p.m.), then diluted 10-fold and further cultivated in SC (24 hrs., 30°C, 250 r.p.m.), whereafter, for each replicate, 50,000 cells were transferred to 300 µL volumes comprising a dilution series of respective ligands dissolved in pH-buffered SC-AS/Urea medium^111^, and stimulated for 20 hrs. (30°C, 250 r.p.m.). DIX45 (Ste2) and DIX59 (No GPCR) was stimulated at pH=5 with 0 µM, 0.01 µM, and 1 µM ɑ-factor (GenScript), DIX60 (Mam2) at pH=5 with 0 µM, 1 µM, and 10 µM P-factor (peptide-seq.: TYADFLRAYQSWNTFVNPDRPNL) (GenScript), DIX61 (MTNR1A) at pH=5 with 0 µM, 1 µM, and 10 µM melatonin (Sigma-Aldrich, Cat.#M5250), DIX62 (ADORA2B) at pH=7 with 0 µM, 0.01 µM, and 1 µM adenosine (Sigma-Aldrich, Cat.#01890), DIX63 (ADRA2A) at pH=7 with 0 µM, 1 µM, and 10 µM clonidine hydrochloride (adrenaline agonist) (Sigma-Aldrich, Cat.#C7897), DIX64 (5HT4b) at pH=5 with 0 µM, 1 µM, and 10 µM serotonin hydrochloride (Sigma-Aldrich, Cat.#H9523), and DIX65 (CXCR4a) at pH=7 by 0 µM, 100 µM, and 1000 µM NUCC-390 hydrochloride (CXCL12 agonist) (Cayman Chemical Company, Cat.#30957).

For yeast cell staining, approx. 500,000 cells were pelleted (800g, 3 min., 5°C), washed twice in 150 µL ice-cold PBSA (1X PBS with 1 g/L bovine serum albumin (Sigma-Aldrich, Cat.#A4503)), then resuspended and incubated in 50 µL primary antibody mix (darkness, 30 min., on ice, 200 r.p.m.). Hereafter, the procedure was repeated for the secondary antibody mix. Finally, cells were pelleted and washed twice in 150 µL ice-cold PBSA, whereafter they were resuspended in 150 µL ice-cold PBSA for flow cytometry using a NovoCyte Quanteon flow cytometer (Agilent). The CD19 epitope was stained with 2.5 µg/mL (1:200) mouse anti-CD19 (FMC63) (Absolute Antibody, Cat.#Ab00613) and 40 µg/mL (1:50) goat anti-mouse-AF647 (polyclonal) (Thermo Fisher, Cat.#A-21236), detected in R667/30-H, and the HA-tag was stained with 2.5 µg/mL (1:400) rabbit anti-HA (RM305) (Thermo Fisher, Cat.#MA5-27915) and 10 µg/mL (1:200) goat anti-rabbit-AF488 (polyclonal) (Thermo Fisher, Cat.#A-11008), detected in B525/45-H (**Fig. 2g-j, Suppl. Fig. 10-12**), or alternatively Anti-HA.11-AF647 (BioLegend, Cat.#682404) detected in R667/30-H (**Fig. 2a-b, Suppl. Fig. 3**). NALM6 was stained analogously for direct comparison of CD19 levels. The gating strategy is described in **Suppl. Fig. 27**. Each condition was examined in three biological replicates and 50,000 cells were sampled per replicate. A one-way or two-way ANOVA with Tukey’s, Dunnett’s, or Šídák’s multiple comparisons test was conducted for the individual conditions, and to fulfill the assumptions of parametric tests, some data was log_10_-transformed, all specified in **Suppl. Table 5-6**.

#### Yeast growth assays in mammalian media

For yeast growth assays (**Suppl. Fig. 13**), strains were seeded at 40,000 cells in 200 µL media and grown at 37°C for 24 hrs in yeast media; SC and pH=5-buffered SC-AS/Urea ^111^, as well as in human immune cell media; RPMI+10%FBS+1%PS and ImmunoCult™-XF T Cell Expansion Medium. Yeast cells were grown in a BioTek Epoch 2 (Agilent) microplate reader to monitor growth at OD_630_ in three biological replicates.

#### T cell viability and proliferation in yeast co-cultivations

For assessing T cell viability and proliferation, as well as yeast biosensing during long-term co-cultivation (**Suppl. Fig. 13**), T cells separated from a donor-derived isolate of PBMCs were co-cultured with a yeast strain equipped with Ste2 and a P_FUS1_-yEGFP cassette (B525/45-A). For co-cultivation, a constant cell number of 500,000 T cells was supplied with ratios of 0.5x (250,000), 1.0x (500,000), or 10x (5,000,000) yeast cells and cultivated in 500 µL RPMI+10%FBS, for each replicate, at 37°C with 5% CO_2_ for 4 days (96 hrs). In addition, the 1.0x co-cultures were assessed with supplementation of 20 µM ɑ-factor (GenScript). Yeast was pre-cultivated in 0.5 mL SC (24 hrs., 30°C, 250 r.p.m.), then diluted 10-fold and further cultivated in SC (24 hrs., 30°C, 250 r.p.m.), and then washed two times in 1X PBS before establishing co-cultures (**Suppl. Fig. 26**). T cells were labeled using a CellTrace™ CFSE Cell Proliferation Kit (Thermo Fisher, Cat.#C34554) (B525/45-A) according to the manufacturer’s protocol before co-culture establishment, and after co-cultivation the entire culture was stained with anti-CD3-BV421 (1:50) (BD Biosciences, Cat.#562426) (V445/45-A), and using a Fixable Near-IR Dead Cell Stain Kit (Thermo Fisher, Cat.#L34975) (R780/60-A) according to the manufacturer’s protocol for assessing T cell viability. For staining, the co-cultures were pelleted (300g, 5 min., 5°C), then washed with 100 µL Cell Staining Buffer (BioLegend, Cat.#420201), whereafter each replicate was resuspended in 50 µL staining mix and incubated (darkness, 30 min., on ice). Then cells were pelleted (300g, 5 min., 5°C), washed with 150 µL ice-cold Cell Staining Buffer, and then resuspended in 150 µL ice-cold Cell Staining Buffer for flow cytometry. The co-cultures were examined in three biological replicates and 50,000 cells were sampled per replicate at day 1 (0 hrs), day 2 (24 hrs), and day 5 (96 hrs) by flow cytometry using a NovoCyte Quanteon flow cytometer (Agilent). The gating strategy is described in **Suppl. Fig. 27**. A two-way ANOVA with Tukey’s multiple comparisons test was conducted for the individual conditions, and to fulfill the assumptions of parametric tests, some data was log_10_-transformed, all specified in **Suppl. Table 7**.

#### CAR Jurkat NFAT-Luc activation assay with SCASA yeast cells and NALM6

For the investigation of SCASA yeast cell activation of CAR Jurkat NFAT-Luc cells with comparison to NALM6 (**Fig. 3, Suppl. Fig. 14**), several co-cultures were set up. Yeast strains (DIX44, DIX45, DIX48, DIX47, and DIX49) were inoculated and pre-cultivated in 0.5 mL SC (24 hrs., 30°C, 250 r.p.m.), then diluted 10-fold and further cultivated (24 hrs., 30°C, 250 r.p.m.), whereafter for GPCR stimulation, for each replicate, cultures were diluted 50-fold by transfer to 200 µL volumes comprising a 10-fold dilution series of ɑ-factor (0-1 µM) (GenScript) in SC+1%DMSO, and then stimulated for 20 hrs. to induce differential CD19 YSD (30°C, 250 r.p.m.). Before establishment of co-cultures, SCASA yeast cells were pelleted (800g, 3 min.), washed twice in 150 µL PBS, and resuspended in RPMI+10%FBS+1%PS. For the establishment of 0-100% CAR+ Jurkat NFAT-Luc cultures, 0% CAR+ and 100% CAR+ Jurkat NFAT-Luc cultures were titrated by cell counting to control the amount of CAR+ cells in each culture. Then, for each replicate of co-cultures, 200,000 alive Jurkat NFAT-Luc cells were co-cultured with 40,000 target cells (i.e. a T/E-ratio of 0.2x SCASA yeast cells or NALM6) for 18 hrs in 200 µL RPMI+10%FBS+1%PS at 37°C with 5% CO_2_ (**Suppl. Fig. 26**). After co-cultivation, cells were lysed using Pierce™ Luciferase Cell Lysis Buffer (Thermo Fisher, Cat.#16189) and the luciferase activity was examined using the Luciferase Assay System (Promega, Cat.#E1500) according to the manufacturers’ protocols. Three replicates were made for each tested condition. Relative activation of Jurkat NFAT-Luc cells was calculated by subtraction of background luminescence and normalization to a 100% CAR+ Jurkat cell mono-culture. A one-way or two-way ANOVA with Tukey’s or Dunnett’s multiple comparisons test was conducted for the individual conditions, and to fulfill the assumptions of parametric tests, some data was log_10_-transformed, all specified in **Suppl. Table 8**.

#### CAR design screening assay in TPR Jurkat cells using SCASA yeast cells

To demonstrate the differential impact of target antigen densities and cellular ratios on different CAR designs, a wide variety of co-cultivations were set up (**Fig. 4, Suppl. Fig. 15-19**). Yeast strains DIX44, DIX45, DIX47, DIX48, DIX49, and DIX67 were inoculated and pre-cultivated in 0.5 mL SC (24 hrs., 30°C, 250 r.p.m.), then diluted 10-fold and further cultivated (24 hrs., 30°C, 250 r.p.m.). For the differential target-cell-ratio conditions, yeast cultures were pelleted (1,200g, 90 sec.), washed twice with PBS, and then resuspended in RPMI+10%FBS+1%PS. Yeast cell densities were then normalized to result in 8.0x-0.25x the number of alive CAR TPR Jurkat cells, by establishing the 8.0x condition through equalizing cell counts, and then serially diluting it to produce 4.0x, 2.0x, 1.0x, 0.5x, and 0.25x yeast cultures. The same was done for NALM6. To establish different antigen-density conditions, each yeast strain was stimulated through the controlling GPCR differentially by 20-fold dilutions into a 10-fold dilution series of ɑ-factor (0-1 µM) (GenScript) in full SC. Strains were incubated for 20 hrs. to induce differential CD19 YSD (30°C, 250 r.p.m.). After induction of antigen-densities, yeast cells were pelleted (1,200g, 90 sec.), washed twice with PBS, and then resuspended in RPMI+10%FBS+1%PS. Then each individual yeast culture, composed of a combination of yeast strain and stimulation level, was normalized to the same cell density to ensure that all CAR TPR Jurkat co-cultures would be exactly 1.0x yeast for all antigen-densities examined. All stimulatory conditions were then set up by combining the target cells with the CD28 CAR and 4-1BB CAR TPR Jurkat cell lines individually, always with 50,000 alive CAR TPR Jurkats cells in a total volume of 200 µL RPMI+10%FBS+1%P/S (1.0x). In addition to the co-cultures, negative control monocultures and positive control cultures with 1.0x Human T-Activator CD3/CD28 Dynabeads (Thermo Scientific, Cat.#11161D) (ɑCD3/ɑCD28) were employed. The cells were then co-cultured for 24 hrs at 37°C with 5% CO_2_. Finally, cells were pelleted (300g, 5 min.) and resuspended in 150 µL ice-cold PBSA for flow cytometry using a NovoCyte Quanteon flow cytometer (Agilent); NF-κB-CFP (V525/45-A), NFAT-eGFP (B525/45-A), and AP-1-mCherry (Y615/20-A). Three biological replicates were made for each condition. Responsiveness analysis was done by log_10_-transforming response data and conducting linear regressions (**Suppl. Table 10**). Two-way ANOVA with Šidák or Tukey’s multiple comparisons test was conducted for the individual analyses, and to fulfill the assumptions of parametric tests, data was log_10_-transformed when relevant, all of which is specified in **Suppl. Table 9-10.**

#### Activation assay of donor-derived CAR T cells with SCASA yeast cells and NALM6

SCASA yeast cell activation of donor-derived CAR T cells was done by co-cultivation with comparison to NALM6 (**Fig. 5, Suppl. Fig. 20-23**). Yeast strains P_PGK1_-Empty (DIX44) and P_PGK1_-CD19 (DIX47) were inoculated and pre-cultivated in 0.5 mL SC (24 hrs., 30°C, 250 r.p.m.), then diluted 10-fold and further cultivated (24 hrs., 30°C, 250 r.p.m.). Before the establishment of co-cultures, SCASA yeast cells were pelleted (800g, 3 min.), washed twice in 150 µL PBS, and resuspended in RPMI+10%FBS+1%PS. For co-cultivation, a constant cell number of 100,000 alive CAR+ T cells was supplied with ratios of 0.2x (20,000), 1.0x (100,000), or 5.0x (500,000) SCASA yeast cells or NALM6 cells and cultivated in 200 µL RPMI+10%FBS+1%PS for each replicate. For CTRL T cell co-cultures, the corresponding number of alive T cells was used to maintain the same overall number of T cells. The co-cultures were incubated for 20 hrs at 37°C with 5% CO_2_ (**Suppl. Fig. 26**). After co-cultivation, the cultures were stained for flow cytometry using anti-CD69-PE/Cy7 (1:150) (BioLegend, Cat.#310912) (Y780/60-A), anti-CD3-perCP (1:30) (BioLegend, Cat.#300326) (B690/50-A), anti-c-myc-DL488 (1:40) (Abcam, Cat.#ab117499) (B525/40-A), anti-CD19-BV785 (1:50) (BioLegend, Cat.#302240) (V780/60-A), and the Zombie Violet Fixable Viability Kit (BioLegend, Cat.#423114) (V445/45-A) according to the manufacturer’s protocol. A master mix was made in Cell Staining Buffer (BioLegend, Cat.#420201). For staining, the co-cultures were first pelleted (300g, 5 min., 5°C) to then carefully aspirate the media, whereafter each replicate was resuspended in 50 µL staining master mix and incubated (darkness, 30 min., on ice). Hereafter, the cells were pelleted (300g, 5 min., 5°C), washed twice with 150 µL ice-cold Cell Staining Buffer, and then resuspended in 100 µL ice-cold Cell Staining Buffer for flow cytometry. Three biological replicates were made for each condition, and measurements were conducted using a CytoFLEX-S (V-B-Y-R) instrument (Beckman Coulter) by analyzing 80 µL of each replicate. The gating strategies are described in **Suppl. Fig. 28**. A one-way, two-way, or three-way ANOVA with Tukey’s multiple comparisons test was conducted for the individual conditions, and to fulfill the assumptions of parametric tests, some data was log_10_-transformed, all specified in **Suppl. Table 11**. The performance summary of the investigated conditions (Fig. 5i) shows the mean performance across all T/E ratios (0.2x, 1.0x, 5.0x) for SCASA yeast P_PGK1_-CD19 and NALM6, for each parameter displayed in the individual plots (Fig. 5a-h). This was done by calculating the mean of all measurements normalized to the global maximum value detected, standardized to 100% for each parameter, regardless of target cell type and T/E ratio.

After 5 months of cryopreservation in liquid nitrogen, the CAR T cell product was thawed for reexamination (**Suppl. Fig. 20**). After thawing, the CAR T cells were recovered by two days of cultivation in ImmunoCult™-XF T Cell Expansion Medium (Stemcell Technologies, Cat.#10981) supplemented with Human Recombinant IL-2 (12.5 ng/mL) (Stemcell Technologies, Cat.#78036), with initial seeding at 1×10^6^ alive cells/mL. Yeast strains P_PGK1_-Empty (DIX44), P_MFA2_-CD19 (DIX48), and P_TDH3_-CD19 (DIX49) were pre-cultivated with and without stimulation using 0.1 µM ɑ-factor (20 hrs., 30°C, 250 r.p.m.), as described for CD19 display characterization. SCASA yeast cells were pelleted (800g, 3 min.), washed twice in 150 µL PBS, and resuspended in RPMI+10%FBS+1%PS. Co-cultures were then established by combining 75,000 alive CAR+ T cells with 25,000 yeast cells (0.3x) in 200 µL RPMI+10%FBS+1%PS. A positive activation control was made using 1X Leukocyte Activation Cocktail (LAC), with BD GolgiPlug™ (BD Biosciences, Cat.#550583) as positive control. After 20 hrs of cultivation at 37°C with 5% CO_2_, cultures were stained for flow cytometry with anti-CD69-PE/Cy7 (1:150) (BioLegend, Cat.#310912) (Y780/60-A), anti-CD25-BB700 (BD Biosciences, Cat.#566448) (B695/40-A), anti-CD3-FITC (BD Biosciences, Cat.#349201) (B525/45-A), and the Zombie Violet Fixable Viability Kit (V445/45-A), as described above. The gating strategies are described in **Suppl. Fig. 28**. Two biological replicates were made for each condition, and measurements were conducted using a NovoCyte Quanteon flow cytometer (Agilent).

### Software

For the collection of flow cytometry data, NovoExpress Software v1.6.2 (Agilent) was employed for controlling the NovoCyte Quanteon system (Agilent) and CytExpert Acquisition and Analysis Software v2.5 (Beckman Coulter) for the CytoFLEX-S system (Beckman Coulter). Flow cytometry data was analyzed using FlowJo™ v10.8 Software (BD Life Sciences). Statistical analyses and plotting were done using GraphPad Prism v9.4.1 (GraphPad Software). Schematic figure illustrations for **Fig. 1**, **2c**, **3a**, **4a**, and **Suppl. Fig. 25-26**, were created with BioRender.com.

## Acknowledgements

This work was supported by the Novo Nordisk Foundation, grant number NNF20CC0035580 to MKJ, grant NNF21OC0066562 "Challenge Programme 2021 - Smart Nanomaterials for Applications in Life-Science" to SRH, the U.S. Department of Energy, grant number DE-SC0018368 to RTG, and the Independent Research Fund Denmark, grant number 0129-00005B to MO. Authors would like to thank Dr. Jie Zhang for providing plasmids pESC-URA-gRNA_XI-3 and pESC-URA-gRNA_XII-5, as well as Professor Tom Ellis and Dr. William Shaw for plasmid pWS1776.

## Author contributions

MD, EDJ, SRH, MO, and MKJ conceived the study. MD led the investigation. MD performed yeast characterization experiments with support from NMTK. MD designed and constructed all yeast strains and plasmids of this study, except for DIX32-34 and pMAD26,29-30 constructed by NMTK. MD, KR, and GS performed Jurkat co-cultivation experiments. RUFW, KR, and MO constructed the CAR Jurkat NFAT-Luc cell line and CAR TPR Jurkat cell lines. KZ, MD, and GS constructed donor-derived CAR T cells and performed co-cultivation experiments. MD performed all data analyses. MD, EDJ, and MKJ wrote the manuscript. MKJ, RTG, SRH, and MO acquired the funding enabling this research. All authors approved the manuscript.

## References

1. Cubillos-Ruiz, A. et al. Engineering living therapeutics with synthetic biology. Nat. Rev. Drug Discov. 20, 941–960 (2021).

2. Rohaan, M. W., Wilgenhof, S. & Haanen, J. B. A. G. Adoptive cellular therapies: the current landscape. Virchows Arch. 474, 449–461 (2019).

3. Cappell, K. M. & Kochenderfer, J. N. Long-term outcomes following CAR T cell therapy: what we know so far. Nat. Rev. Clin. Oncol. 20, 359–371 (2023).

4. Lee, D. W. et al. T cells expressing CD19 chimeric antigen receptors for acute lymphoblastic leukaemia in children and young adults: a phase 1 dose-escalation trial. Lancet 385, 517–528 (2015).

5. Park, J. H. et al. Long-Term Follow-up of CD19 CAR Therapy in Acute Lymphoblastic Leukemia. N. Engl. J. Med. 378, 449–459 (2018).

6. Maude, S. L. et al. Chimeric antigen receptor T cells for sustained remissions in leukemia. N. Engl. J. Med. 371, 1507–1517 (2014).

7. Maude, S. L. et al. Tisagenlecleucel in Children and Young Adults with B-Cell Lymphoblastic Leukemia. N. Engl. J. Med. 378, 439–448 (2018).

8. Si Lim, S. J., Grupp, S. A. & DiNofia, A. M. Tisagenlecleucel for treatment of children and young adults with relapsed/refractory B-cell acute lymphoblastic leukemia. Pediatr. Blood Cancer 68, e29123 (2021).

9. Zhang, X. et al. Efficacy and safety of anti-CD19 CAR T-cell therapy in 110 patients with B-cell acute lymphoblastic leukemia with high-risk features. Blood Adv 4, 2325–2338 (2020).

10. Liu, S. et al. Which one is better for refractory/relapsed acute B-cell lymphoblastic leukemia: Single-target (CD19) or dual-target (tandem or sequential CD19/CD22) CAR T-cell therapy? Blood Cancer J. 13, 60 (2023).

11. Pasquini, M. C. et al. Real-world evidence of tisagenlecleucel for pediatric acute lymphoblastic leukemia and non-Hodgkin lymphoma. Blood Adv 4, 5414–5424 (2020).

12. Schultz, L. M. et al. Outcomes After Nonresponse and Relapse Post-Tisagenlecleucel in Children, Adolescents, and Young Adults With B-Cell Acute Lymphoblastic Leukemia. J. Clin. Oncol. 41, 354–363 (2023).

13. Dourthe, M.-E. et al. Determinants of CD19-positive vs CD19-negative relapse after tisagenlecleucel for B-cell acute lymphoblastic leukemia. Leukemia 35, 3383–3393 (2021).

14. Nie, Y. et al. Mechanisms underlying CD19-positive ALL relapse after anti-CD19 CAR T cell therapy and associated strategies. Biomark Res 8, 18 (2020).

15. Xu, X. et al. Mechanisms of Relapse After CD19 CAR T-Cell Therapy for Acute Lymphoblastic Leukemia and Its Prevention and Treatment Strategies. Front. Immunol. 10, 2664 (2019).

16. Bai, Z. et al. Single-cell antigen-specific landscape of CAR T infusion product identifies determinants of CD19-positive relapse in patients with ALL. Sci Adv 8, eabj2820 (2022).

17. Majzner, R. G. & Mackall, C. L. Tumor Antigen Escape from CAR T-cell Therapy. Cancer Discov. 8, 1219–1226 (2018).

18. Sotillo, E. et al. Convergence of Acquired Mutations and Alternative Splicing of CD19 Enables Resistance to CART-19 Immunotherapy. Cancer Discov. 5, 1282–1295 (2015).

19. Orlando, E. J. et al. Genetic mechanisms of target antigen loss in CAR19 therapy of acute lymphoblastic leukemia. Nat. Med. 24, 1504–1506 (2018).

20. Fischer, J. et al. CD19 Isoforms Enabling Resistance to CART-19 Immunotherapy Are Expressed in B-ALL Patients at Initial Diagnosis. J. Immunother. 40, 187–195 (2017).

21. Braig, F. et al. Resistance to anti-CD19/CD3 BiTE in acute lymphoblastic leukemia may be mediated by disrupted CD19 membrane trafficking. Blood 129, 100–104 (2017).

22. Ledererova, A. et al. Hypermethylation of CD19 promoter enables antigen-negative escape to CART-19 in vivo and in vitro. J Immunother Cancer 9, (2021).

23. Gardner, R. et al. Acquisition of a CD19-negative myeloid phenotype allows immune escape of MLL-rearranged B-ALL from CD19 CAR-T-cell therapy. Blood 127, 2406–2410 (2016).

24. Jacoby, E. et al. CD19 CAR immune pressure induces B-precursor acute lymphoblastic leukaemia lineage switch exposing inherent leukaemic plasticity. Nat. Commun. 7, 12320 (2016).

25. Hamieh, M. et al. CAR T cell trogocytosis and cooperative killing regulate tumour antigen escape. Nature 568, 112–116 (2019).

26. Maude, S. L. et al. Sustained remissions with CD19-specific chimeric antigen receptor (CAR)-modified T cells in children with relapsed/refractory ALL. J. Clin. Orthod. 34, 3011–3011 (2016).

27. Lee, D. W., III et al. Long-term outcomes following CD19 CAR T cell therapy for B-ALL are superior in patients receiving a fludarabine/cyclophosphamide preparative regimen and post-CAR hematopoietic stem cell transplantation. Blood 128, 218–218 (2016).

28. Gardner, R. A. et al. Intent-to-treat leukemia remission by CD19 CAR T cells of defined formulation and dose in children and young adults. Blood 129, 3322–3331 (2017).

29. June, C. H., O’Connor, R. S., Kawalekar, O. U., Ghassemi, S. & Milone, M. C. CAR T cell immunotherapy for human cancer. Science 359, 1361–1365 (2018).

30. Fry, T. J. et al. CD22-targeted CAR T cells induce remission in B-ALL that is naive or resistant to CD19-targeted CAR immunotherapy. Nat. Med. 24, 20–28 (2018).

31. Walker, A. J. et al. Tumor Antigen and Receptor Densities Regulate Efficacy of a Chimeric Antigen Receptor Targeting Anaplastic Lymphoma Kinase. Mol. Ther. 25, 2189–2201 (2017).

32. Hombach, A. A. et al. Superior Therapeutic Index in Lymphoma Therapy: CD30(+) CD34(+) Hematopoietic Stem Cells Resist a Chimeric Antigen Receptor T-cell Attack. Mol. Ther. 24, 1423–1434 (2016).

33. Watanabe, K. et al. Target antigen density governs the efficacy of anti-CD20-CD28-CD3 ζ chimeric antigen receptor-modified effector CD8+ T cells. J. Immunol. 194, 911–920 (2015).

34. Majzner, R. G. et al. Low CD19 Antigen Density Diminishes Efficacy of CD19 CAR T Cells and Can be Overcome By Rational Redesign of CAR Signaling Domains. Blood 132, 963 (2018).

35. Majzner, R. G. et al. Tuning the Antigen Density Requirement for CAR T-cell Activity. Cancer Discov. 10, 702–723 (2020).

36. Pillai, V. et al. CAR T-cell therapy is effective for CD19-dim B-lymphoblastic leukemia but is impacted by prior blinatumomab therapy. Blood Adv 3, 3539–3549 (2019).

37. Meléndez, A. V. et al. Novel lectin-based chimeric antigen receptors target Gb3-positive tumour cells. Cell. Mol. Life Sci. 79, 513 (2022).

38. Burton, J. et al. Inefficient exploitation of accessory receptors reduces the sensitivity of chimeric antigen receptors. Proc. Natl. Acad. Sci. U. S. A. 120, e2216352120 (2023).

39. Beppler, C. et al. Hyperstabilization of T cell microvilli contacts by chimeric antigen receptors. J. Cell Biol. 222, (2023).

40. Wang, V., Gauthier, M., Decot, V., Reppel, L. & Bensoussan, D. Systematic Review on CAR-T Cell Clinical Trials Up to 2022: Academic Center Input. Cancers 15, (2023).

41. Cappell, K. M. & Kochenderfer, J. N. A comparison of chimeric antigen receptors containing CD28 versus 4-1BB costimulatory domains. Nat. Rev. Clin. Oncol. 18, 715–727 (2021).

42. Haso, W. et al. Anti-CD22-chimeric antigen receptors targeting B-cell precursor acute lymphoblastic leukemia. Blood 121, 1165–1174 (2013).

43. Salzer, B. et al. Engineering AvidCARs for combinatorial antigen recognition and reversible control of CAR function. Nat. Commun. 11, 4166 (2020).

44. Patel, A. et al. Using CombiCells, a platform for titration and combinatorial display of cell surface ligands, to study T-cell antigen sensitivity modulation by accessory receptors. EMBO J. 43, 132–150 (2024).

45. Hu, Y. et al. Antigen-Multimers: Specific, Sensitive, Precise, and Multifunctional High-Avidity CAR-Staining Reagents. Matter 4, 3917–3940 (2021).

46. Zhang, Y. et al. Extracellular Vesicles Expressing CD19 Antigen Improve Expansion and Efficacy of CD19-Targeted CAR-T Cells. Int. J. Nanomedicine 18, 49–63 (2023).

47. Landgraf, K. E. et al. convertibleCARs: A chimeric antigen receptor system for flexible control of activity and antigen targeting. Commun Biol 3, 296 (2020).

48. Cook, W. J., Choi, Y., Gacerez, A., Bailey-Kellogg, C. & Sentman, C. L. A Chimeric Antigen Receptor That Binds to a Conserved Site on MICA. Immunohorizons 4, 597–607 (2020).

49. Gudipati, V. et al. Inefficient CAR-proximal signaling blunts antigen sensitivity. Nat. Immunol. 21, 848–856 (2020).

50. Cho, B. K., Kieke, M. C., Boder, E. T., Wittrup, K. D. & Kranz, D. M. A yeast surface display system for the discovery of ligands that trigger cell activation. J. Immunol. Methods 220, 179–188 (1998).

51. Boder, E. T. & Wittrup, K. D. Yeast surface display for screening combinatorial polypeptide libraries. Nat. Biotechnol. 15, 553–557 (1997).

52. Deichmann, M., Hansson, F. G. & Jensen, E. D. Yeast-based screening platforms to understand and improve human health. Trends Biotechnol. (2024) doi:10.1016/j.tibtech.2024.04.003.

53. Rappazzo, C. G., Huisman, B. D. & Birnbaum, M. E. Repertoire-scale determination of class II MHC peptide binding via yeast display improves antigen prediction. Nat. Commun. 11, 4414 (2020).

54. Gee, M. H. et al. Antigen Identification for Orphan T Cell Receptors Expressed on Tumor-Infiltrating Lymphocytes. Cell 172, 549–563.e16 (2018).

55. Wen, F., Esteban, O. & Zhao, H. Rapid identification of CD4+ T-cell epitopes using yeast displaying pathogen-derived peptide library. J. Immunol. Methods 336, 37–44 (2008).

56. Smith, M. R., Tolbert, S. V. & Wen, F. Protein-Scaffold Directed Nanoscale Assembly of T Cell Ligands: Artificial Antigen Presentation with Defined Valency, Density, and Ratio. ACS Synth. Biol. 7, 1629–1639 (2018).

57. Brophy, S. E., Holler, P. D. & Kranz, D. M. A yeast display system for engineering functional peptide-MHC complexes. J. Immunol. Methods 272, 235–246 (2003).

58. Wen, F., Sethi, D. K., Wucherpfennig, K. W. & Zhao, H. Cell surface display of functional human MHC class II proteins: yeast display versus insect cell display. Protein Eng. Des. Sel. 24, 701–709 (2011).

59. Wang, L. & Lan, X. Rapid screening of TCR-pMHC interactions by the YAMTAD system. Cell Discov 8, 30 (2022).

60. Boder, E. T., Bill, J. R., Nields, A. W., Marrack, P. C. & Kappler, J. W. Yeast surface display of a noncovalent MHC class II heterodimer complexed with antigenic peptide. Biotechnol. Bioeng. 92, 485–491 (2005).

61. Raeeszadeh-Sarmazdeh, M. & Boder, E. T. Yeast Surface Display: New Opportunities for a Time-Tested Protein Engineering System. in Yeast Surface Display (ed. Traxlmayr, M. W.) 3–25 (Springer US, New York, NY, 2022).

62. Ferreira, R., Limeta, A. & Nielsen, J. Tackling Cancer with Yeast-Based Technologies. Trends Biotechnol. 37, 592–603 (2019).

63. Jensen, E. D. et al. Engineered cell differentiation and sexual reproduction in probiotic and mating yeasts. Nat. Commun. 13, 6201 (2022).

64. Shaw, W. M. et al. Engineering a Model Cell for Rational Tuning of GPCR Signaling. Cell 177, 782–796.e27 (2019).

65. Kapolka, N. J. et al. DCyFIR: a high-throughput CRISPR platform for multiplexed G protein-coupled receptor profiling and ligand discovery. Proc. Natl. Acad. Sci. U. S. A. 117, 13117–13126 (2020).

66. Kapolka, N. J. et al. Proton-gated coincidence detection is a common feature of GPCR signaling. Proc. Natl. Acad. Sci. U. S. A. 118, (2021).

67. Lengger, B. & Jensen, M. K. Engineering G protein-coupled receptor signalling in yeast for biotechnological and medical purposes. FEMS Yeast Res. 20, (2020).

68. Klesmith, J. R. et al. Retargeting CD19 Chimeric Antigen Receptor T Cells via Engineered CD19-Fusion Proteins. Mol. Pharm. 16, 3544–3558 (2019).

69. Klesmith, J. R., Wu, L., Lobb, R. R., Rennert, P. D. & Hackel, B. J. Fine Epitope Mapping of the CD19 Extracellular Domain Promotes Design. Biochemistry 58, 4869–4881 (2019).

70. Younger, D., Berger, S., Baker, D. & Klavins, E. High-throughput characterization of protein–protein interactions by reprogramming yeast mating. Proceedings of the National Academy of Sciences 114, 12166–12171 (2017).

71. Teymennet-Ramírez, K. V., Martínez-Morales, F. & Trejo-Hernández, M. R. Yeast Surface Display System: Strategies for Improvement and Biotechnological Applications. Front Bioeng Biotechnol 9, 794742 (2021).

72. Sun, H. et al. Display of heterologous proteins on the Saccharomyces cerevisiae surface display system using a single constitutive expression vector. Biotechnol. Prog. 30, 443–450 (2014).

73. Hawkins, K. M. & Smolke, C. D. The regulatory roles of the galactose permease and kinase in the induction response of the GAL network in Saccharomyces cerevisiae. J. Biol. Chem. 281, 13485–13492 (2006).

74. Lopez-Morales, J., Vanella, R., Kovacevic, G., Santos, M. S. & Nash, M. A. Titrating Avidity of Yeast-Displayed Proteins Using a Transcriptional Regulator. ACS Synth. Biol. 12, 419–431 (2023).

75. Sanford, A., Kiriakov, S. & Khalil, A. S. A Toolkit for Precise, Multigene Control in Saccharomyces cerevisiae. ACS Synth. Biol. 11, 3912–3920 (2022).

76. Alexander, S. P. H. et al. THE CONCISE GUIDE TO PHARMACOLOGY 2019/20: G protein-coupled receptors. Br. J. Pharmacol. 176 Suppl 1, S21–S141 (2019).

77. Hilger, D., Masureel, M. & Kobilka, B. K. Structure and dynamics of GPCR signaling complexes. Nat. Struct. Mol. Biol. 25, 4–12 (2018).

78. Hagen, D. C., McCaffrey, G. & Sprague, G. F., Jr. Pheromone response elements are necessary and sufficient for basal and pheromone-induced transcription of the FUS1 gene of Saccharomyces cerevisiae. Mol. Cell. Biol. 11, 2952–2961 (1991).

79. Lengger, B. et al. Serotonin G Protein-Coupled Receptor-Based Biosensing Modalities in Yeast. ACS Sens 7, 1323–1335 (2022).

80. Wang, Z.-X., Broach, J. R. & Peiper, S. C. Functional expression of CXCR4 in Saccharomyces cerevisiae in the development of powerful tools for the pharmacological characterization of CXCR4. Methods Mol. Biol. 332, 115–127 (2006).

81. Roberts, C. J. et al. Signaling and circuitry of multiple MAPK pathways revealed by a matrix of global gene expression profiles. Science 287, 873–880 (2000).

82. Reider Apel, A., et al. A Cas9-based toolkit to program gene expression in *Saccharomyces cerevisiae*. Nucleic Acids Res. 45, 496–508 (2017).

83. Ormhøj, M. et al. Chimeric Antigen Receptor T Cells Targeting CD79b Show Efficacy in Lymphoma with or without Cotargeting CD19. Clin. Cancer Res. 25, 7046–7057 (2019).

84. Jutz, S. et al. Assessment of costimulation and coinhibition in a triple parameter T cell reporter line: Simultaneous measurement of NF-κB, NFAT and AP-1. J. Immunol. Methods 430, 10–20 (2016).

85. Huang, W. et al. NFAT and NF-κB dynamically co-regulate TCR and CAR signaling responses in human T cells. Cell Rep. 42, 112663 (2023).

86. Mohr, M. et al. The CRISPR-Cas12a Platform for Accurate Genome Editing, Gene Disruption, and Efficient Transgene Integration in Human Immune Cells. ACS Synth. Biol. 12, 375–389 (2023).

87. Ernst, M. et al. Chimeric antigen receptor (CAR) T-cell therapy for people with relapsed or refractory diffuse large B-cell lymphoma. Cochrane Database Syst. Rev. 9, CD013365 (2021).

88. Brudno, J. N. et al. Safety and feasibility of anti-CD19 CAR T cells with fully human binding domains in patients with B-cell lymphoma. Nat. Med. 26, 270–280 (2020).

89. Blanco, B. et al. Overcoming CAR-Mediated CD19 Downmodulation and Leukemia Relapse with T Lymphocytes Secreting Anti-CD19 T-cell Engagers. Cancer Immunol Res 10, 498–511 (2022).

90. Schneider, D. et al. A tandem CD19/CD20 CAR lentiviral vector drives on-target and off-target antigen modulation in leukemia cell lines. J Immunother Cancer 5, 42 (2017).

91. Fioretti, S., Matson, C. A., Rosenberg, K. M. & Singh, N. J. Host B cells escape CAR-T immunotherapy by reversible downregulation of CD19. Cancer Immunol. Immunother. 72, 257–264 (2023).

92. Eyquem, J. et al. Targeting a CAR to the TRAC locus with CRISPR/Cas9 enhances tumour rejection. Nature 543, 113–117 (2017).

93. Cliff, E. R. S. et al. High Cost of Chimeric Antigen Receptor T-Cells: Challenges and Solutions. Am Soc Clin Oncol Educ Book 43, e397912 (2023).

94. Sterner, R. C. & Sterner, R. M. CAR-T cell therapy: current limitations and potential strategies. Blood Cancer J. 11, 69 (2021).

95. Xiong, Y., Libby, K. A. & Su, X. The physical landscape of CAR-T synapse. Biophys. J. (2023) doi:10.1016/j.bpj.2023.09.004.

96. Xiao, Q., et al. Size-dependent activation of CAR-T cells. Sci Immunol 7, eabl3995 (2022).

97. Vera, J. et al. The CD5 ectodomain interacts with conserved fungal cell wall components and protects from zymosan-induced septic shock-like syndrome. Proc. Natl. Acad. Sci. U. S. A. 106, 1506–1511 (2009).

98. Sato, M. et al. Direct binding of Toll-like receptor 2 to zymosan, and zymosan-induced NF-kappa B activation and TNF-alpha secretion are down-regulated by lung collectin surfactant protein A. J. Immunol. 171, 417–425 (2003).

99. Dunne, M. R., Wagener, J., Loeffler, J., Doherty, D. G. & Rogers, T. R. Unconventional T cells - New players in antifungal immunity. Clin. Immunol. 227, 108734 (2021).

100. Sharma, J., Mudalagiriyappa, S. & Nanjappa, S. G. T cell responses to control fungal infection in an immunological memory lens. Front. Immunol. 13, 905867 (2022).

101. Nouri, Y., Weinkove, R. & Perret, R. T-cell intrinsic Toll-like receptor signaling: implications for cancer immunotherapy and CAR T-cells. J Immunother Cancer 9, (2021).

102. Mercier, B. C., Cottalorda, A., Coupet, C.-A., Marvel, J. & Bonnefoy-Bérard, N. TLR2 engagement on CD8 T cells enables generation of functional memory cells in response to a suboptimal TCR signal. J. Immunol. 182, 1860–1867 (2009).

103. Lai, Y. et al. Toll-like receptor 2 costimulation potentiates the antitumor efficacy of CAR T Cells. Leukemia 32, 801–808 (2018).

104. Jessop-Fabre, M. M. et al. EasyClone-MarkerFree: A vector toolkit for marker-less integration of genes into Saccharomyces cerevisiae via CRISPR-Cas9. Biotechnol. J. 11, 1110–1117 (2016).

105. Nour-Eldin, H. H., Hansen, B. G., Nørholm, M. H. H., Jensen, J. K. & Halkier, B. A. Advancing uracil-excision based cloning towards an ideal technique for cloning PCR fragments. Nucleic Acids Res. 34, e122 (2006).

106. Jensen, E. D. et al. A synthetic RNA-mediated evolution system in yeast. Nucleic Acids Res. 49, e88 (2021).

107. Chao, G. et al. Isolating and engineering human antibodies using yeast surface display. Nat. Protoc. 1, 755–768 (2006).

108. DiCarlo, J. E. et al. Genome engineering in Saccharomyces cerevisiae using CRISPR-Cas systems. Nucleic Acids Res. 41, 4336–4343 (2013).

109. D’Ambrosio, V. et al. A FAIR-compliant parts catalogue for genome engineering and expression control in Saccharomyces cerevisiae. Synth Syst Biotechnol 7, 657–663 (2022).

110. Inoue, H., Nojima, H. & Okayama, H. High efficiency transformation of Escherichia coli with plasmids. Gene 96, 23–28 (1990).

111. Prins, R. C. & Billerbeck, S. A buffered media system for yeast batch culture growth. BMC Microbiol. 21, 127 (2021).

112. Gietz, R. D. & Schiestl, R. H. Quick and easy yeast transformation using the LiAc/SS carrier DNA/PEG method. Nat. Protoc. 2, 35–37 (2007).

113. Vlassis, A., et al. CRISPR-Cas12a-integrated transgenes in genomic safe-harbors retain high expression in human hematopoietic iPSC-derived lineages and primary cells. iScience 108287 (2023).

