## Supplementary Information for "Engineered yeast cells simulating CD19+ cancers to control CAR T cell activation"

### Suppl. Fig. 1

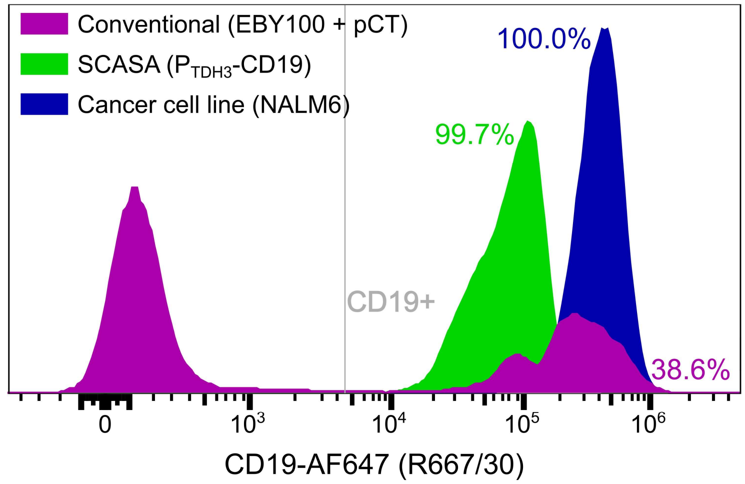


**Suppl. Fig. 1: Comparison of SCASA system performance to cancer cell expression of CD19 and conventional YSD systems.** Conventional YSD of CD19 was assessed using the traditional EBY100 strain containing a pCT-type plasmid with galactose-inducible ($P_{GAL1}$ ) CD19 expression (2%Gal, 24 hrs.). The SCASA yeast cell design expressed CD19 constitutively via $P_{TDH3}$ from a genome-integrated cassette. The human CD19+ NALM6 B cell leukemia cell line was included as a benchmark control. The percentage of alive CD19+ cells is provided. Histograms are normalized to the unit area and are representative replicates.

### Suppl. Fig. 2

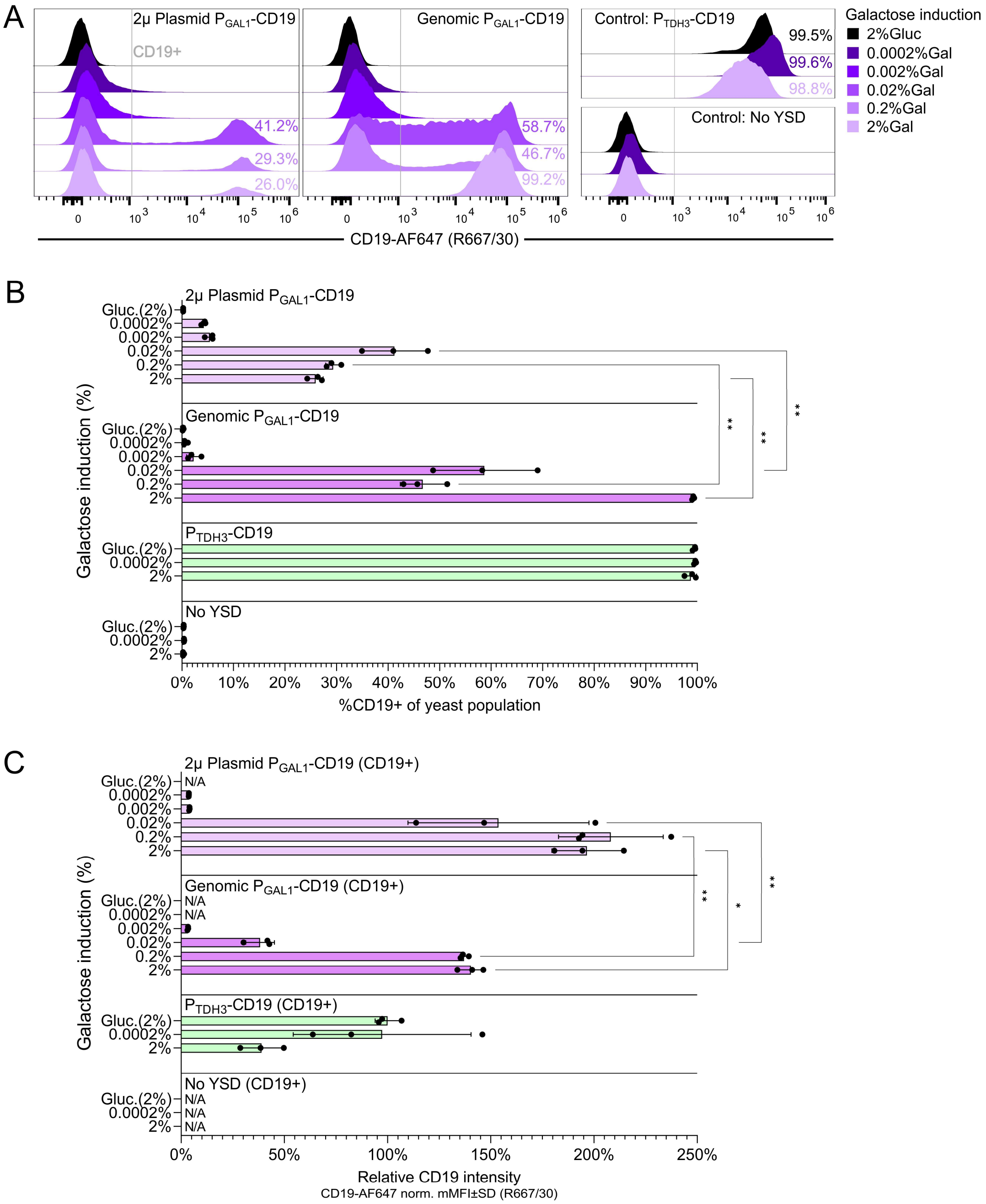


**Suppl. Fig. 2 - CD19 YSD using galactose induction from plasmid and genome-integrated cassettes.** Investigation of $P_{GAL1}$-based control of YSD from a 2µ multicopy expression plasmid (pMAD58) and a genome-integrated site. Cells were stained and analyzed by flow cytometry after 24 hrs of galactose induction (0.0002%-2%Gal), using raffinose as the compensating carbon source. The non-displaying background strain was employed as negative control ('No YSD') and $P_{TDH3}$-CD19 as positive control. A negative control condition was included at which cells were cultured in glucose (2%Gluc). **A**. Comparison of populational behavior in relation to CD19 YSD levels upon stimulation with different concentrations of galactose (*purple*) or glucose (*black*). Percentages on histograms designate the percentage of yeast that express CD19 (%CD19+) based on the shown CD19+ gate. **B**. Comparison of the %CD19+ population sizes of different designs across stimulation levels. **C**. Comparison of CD19 intensities of gated CD19+ cells of different designs across stimulation levels. Data represents means of cell counts or median fluorescence intensities (mMFI) for three biological replicates (n=3) and standard deviations hereof. Histograms are normalized to the mode and are representative replicates. Significance levels: ✱: P≤0.05, ✱✱: P≤0.001. Not all pairwise comparisons are shown. Statistics in: **Suppl. Table 5**.

### Suppl. Fig. 3

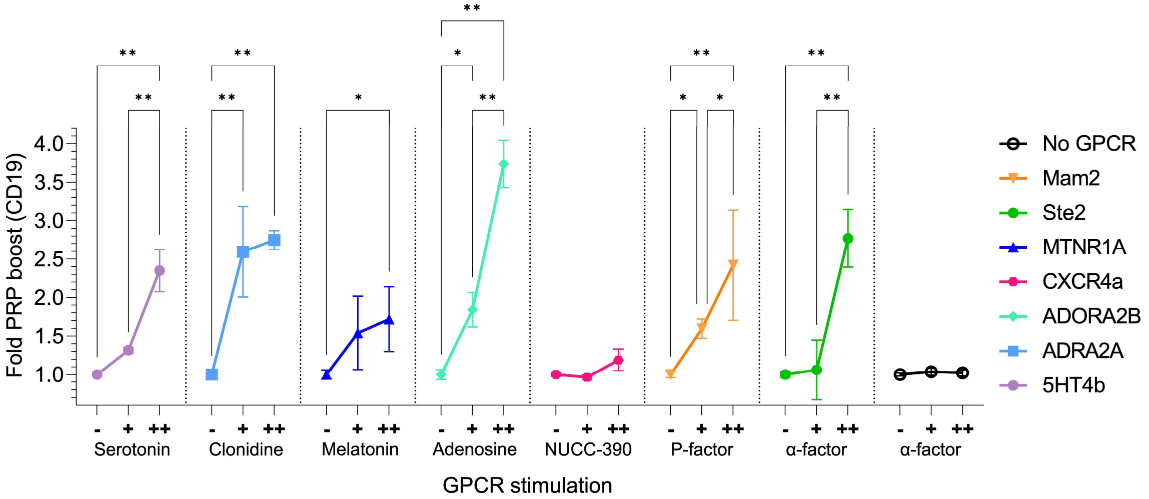


**Suppl. Fig. 3 - PRP activation and approximated PRP boost for GPCR library strains**. Characterization of PRP activation for strains employing heterologously expressed human and fungal GPCRs used for controlling CD19 densities. All strains employ the $P_{FUS1}$ processing module and only vary in the type of GPCR and $G_{\alpha}$-subunit. The fold change in quantified PRP boost in response to GPCR stimulation is provided for each strain in relation to the unstimulated condition. Strains were stimulated with different concentrations of cognate ligands, from none (**-**) to medium (**+**) and high levels (**++**). Data represents means of median fluorescence intensity (mMFI) for three biological replicates (n=3) and standard deviations hereof. Significance levels: ✱: P≤0.05, ✱✱: P≤0.001. Statistics in: **Suppl. Table 6**.

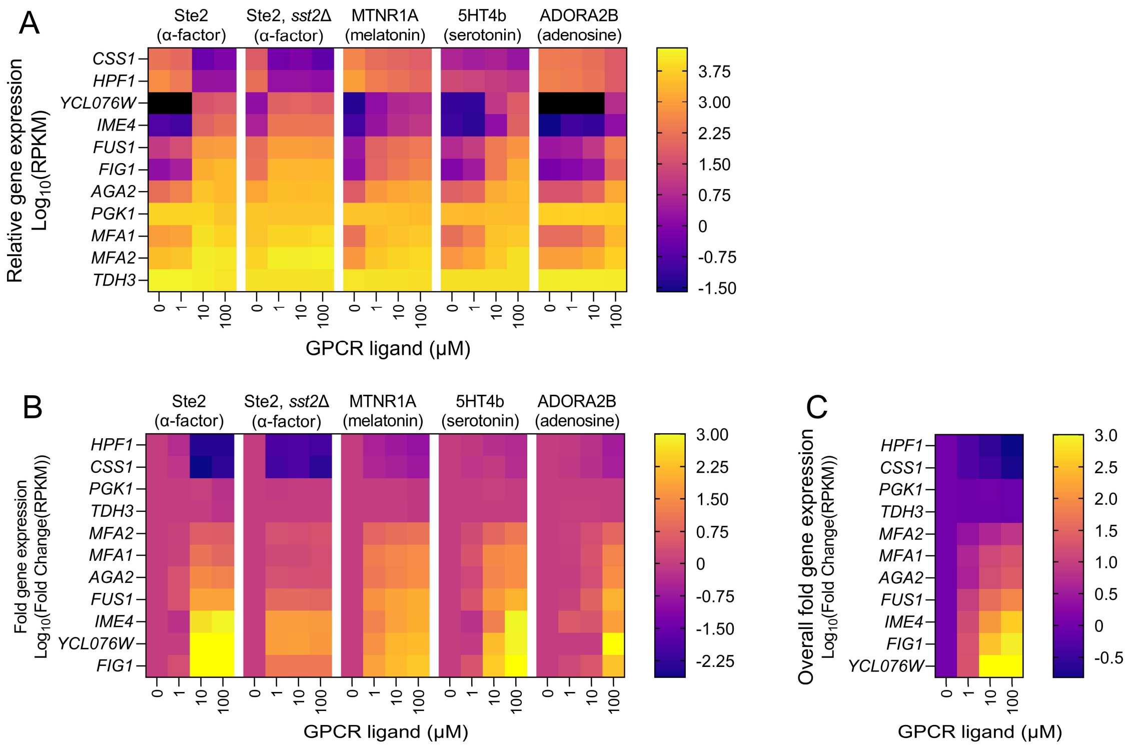


**Suppl. Fig. 4**: **Transcriptome analysis of candidate genes for supplying promoter parts for the SCASA processing module design**. Gene expression patterns were analyzed in five different yeast strains throughout stimulation of their GPCRs using 0-100 µM of their cognate ligands: Ste2 in a wild-type strain, Ste2 in an *sst2Δ* strain, as well as MTNR1A, 5HT4b, and ADORA2B in optimized biosensor strains. Based on the gene expression patterns during GPCR stimulation, the 11 shown genes were chosen as sources for promoter parts. The results are a re-analysis of data from our previous study[1]. **A**. Relative gene expression quantified as $log_{10}$-transformed trimmed mean of M values (TMM) normalized reads per kilobase of transcript per million mapped reads (RPKM). Genes are sorted based on maximal gene expression level from low to high across strains and stimulation levels. **B**. Fold change gene expression upon GPCR stimulation relative to the unstimulated condition for each gene and strain individually ($log_{10}$-transformed values). **C**. Overall fold change gene expression across all strain designs and different GPCRs ($log_{10}$-transformed values). See **Suppl. Table 1** for raw values and fold change values.

### Suppl. Fig. 5

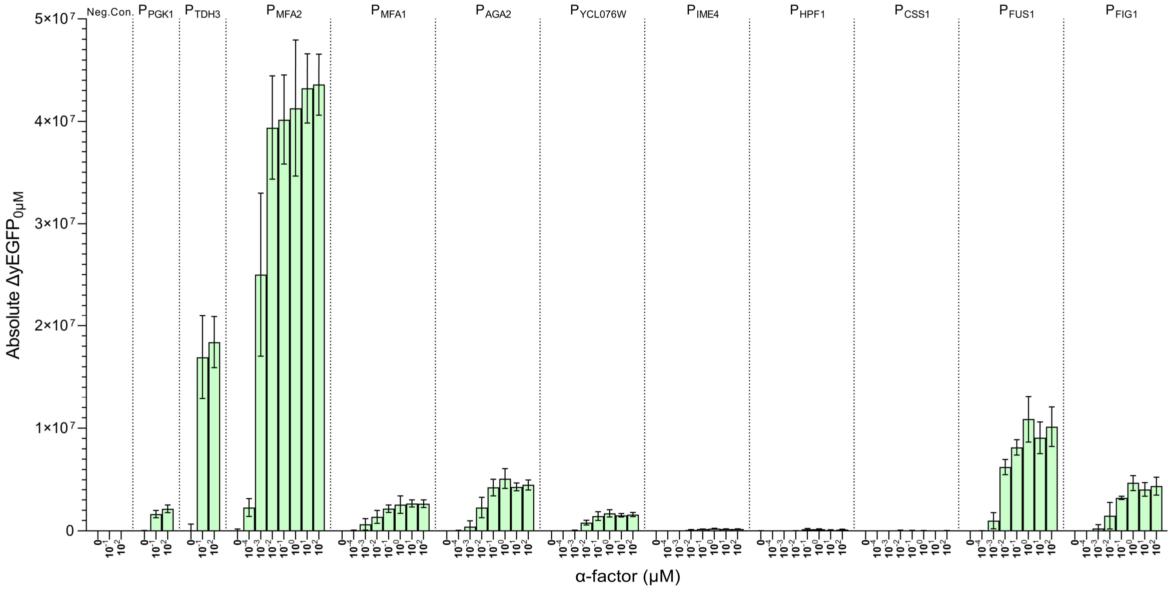


**Suppl. Fig. 5: Differences in absolute yEGFP levels upon GPCR stimulation for the processing module promoter library.** Absolute changes in yEGFP levels are calculated relative to the background level at 0 µM ($\Delta yEGFP_{0\mu M} = yEGFP_X - yEGFP_{0\mu M}$) for each strain design across different concentrations of α-factor (0-100 µM). Data represents means of median fluorescence intensity (mMFI) for three biological replicates (n=3) and standard deviations hereof. Statistics in: **Suppl. Table 2** (not shown).

**Suppl. Fig. 6**

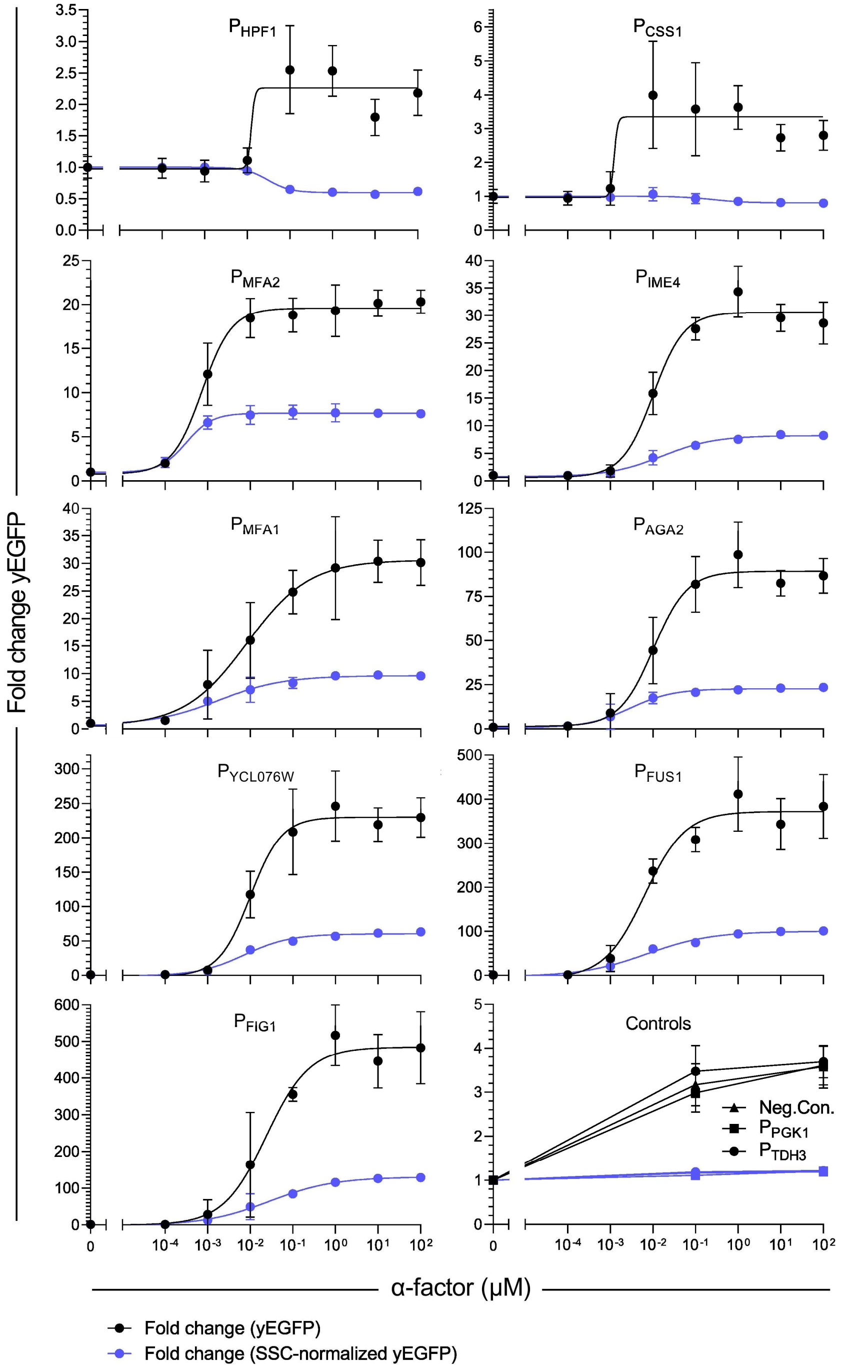


**Suppl. Fig. 6 - Comparison of fold change absolute yEGFP dose-response curves with and without PRP boost.** Fold change yEGFP from the individual processing module promoters, including PRP boost (*black*) and excluding PRP boost via SSC-normalization (*blue*), with curve fitting using variable slope (four-parameter) nonlinear regression. Strains employ the GPCR Ste2 to sense α-factor (0-100 µM). Data represents means of median fluorescence intensity (mMFI) for three biological replicates (n=3) and standard deviations hereof. Statistics and curve fits: **Suppl. Table 4.**

**Suppl. Fig. 7**

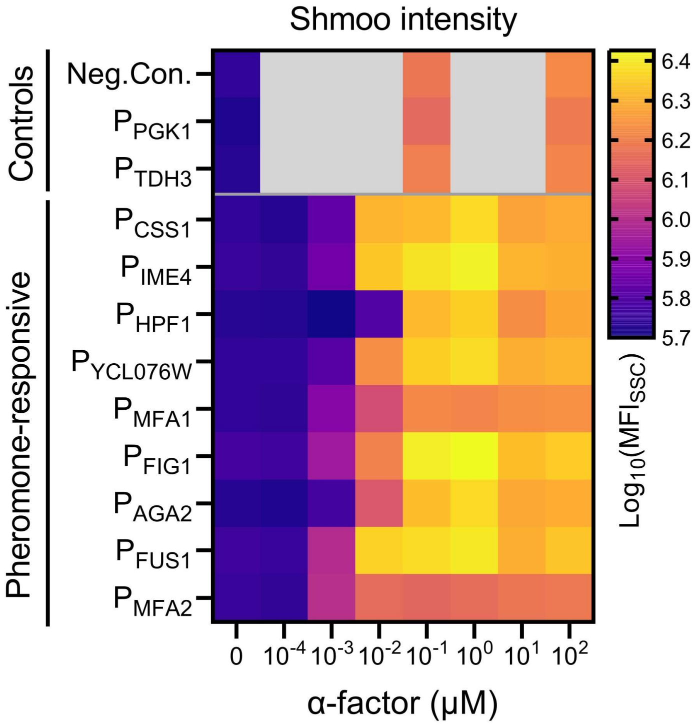


**Suppl. Fig. 7 – Quantification of PRP activation by morphological changes (shmoo intensity).** Intensity of PRP-regulated morphological changes (shmooing) of cells as a measure of PRP activation ($log_{10}$-scale), quantified by flow cytometry (SSC). The promoters are sorted on maximum absolute yEGFP intensity, from low to high (as in **Fig. 2d**). Data represents means of median fluorescence intensity (mMFI) for biological replicates (n=3). All statistics and extended analyses in **Suppl. Table 2.**

### Suppl. Fig. 8

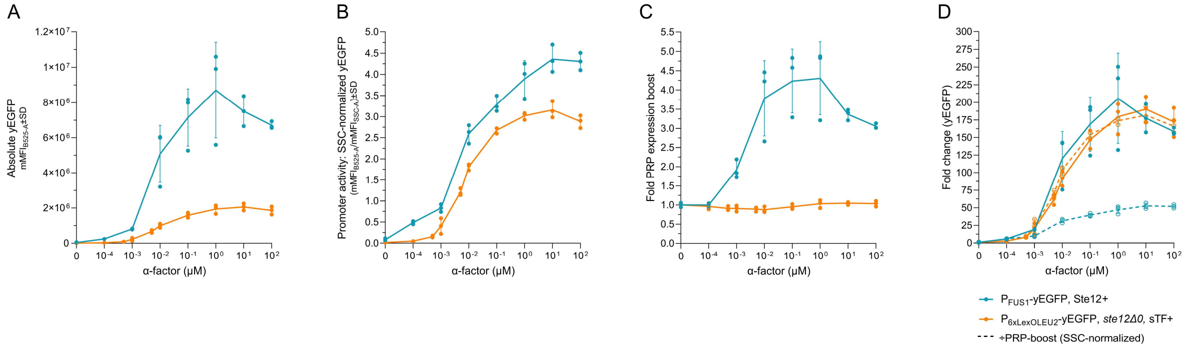


**Suppl. Fig. 8: Comparison of GPCR signaling using Ste12 to an orthogonal synthetic transcription factor.** The dynamics of two biosensor strains employing the GPCR Ste2 to sense α-factor were compared by flow cytometry measurements of yEGFP mean fluorescence intensity (MFI) and morphological changes (SSC). Specifically, we compared the $P_{FUS1}$ processing unit design (*blue*) to orthogonal GPCR signaling that employs an orthogonal signal pathway relying on a LexA-based synthetic transcription factor (sTF) and a LexO-containing promoter ($P_{6xLexOLEU2}$), and importantly has the PRP transcription factor Ste12 deleted (*orange*) [2]. **A**. Absolute yEGFP MFI. **B**. SSC-normalized yEGFP levels. **C**. Quantification of the PRP expression boost. **D**. Fold change yEGFP, with and without SSC-normalization. SSC-normalized fold changes are illustrated with dashed lines. Data represents means of median fluorescence intensity (mMFI) for three biological replicates (n=3) and standard deviations hereof. Statistics in: **Suppl. Table 2** (not shown).

**Suppl. Fig. 9**

### Additional logical analysis of the PRP boost effect

Absolute changes in yEGFP (ΔyEGFP) between $P_{PGK1}$ and $P_{TDH3}$ were significantly different, indicating that the PRP boost is not an additive effect of constant value - i.e. not a general effect that applies equally across promoters, but instead is dependent on the individual promoter (**Suppl. Fig. 9A**). Interestingly, fold changes in yEGFP were identical between $P_{PGK1}$ and $P_{TDH3}$, indicating that the PRP boost effect is a multiplicative effect, which applies as a constant to all promoters with an intensity correlated to the basal expression level of the individual promoter (**Suppl. Fig. 9B**). Additionally, the yEGFP fold change patterns for $P_{PGK1}$ and $P_{TDH3}$ resembled those of PRP-induced morphological changes (shmooing), measured as SSC, and these were also identical between the two strains (**Suppl. Fig. 9B**). Positive linear correlation between yEGFP intensity and PRP activation for constitutive promoter constructs $P_{TDH3}$- and $P_{PGK1}$-yEGFP indicated that it was in fact the PRP activation that caused increased expression of yEGFP (**Suppl. Fig. 9C**), despite constant mRNA levels of the cognate genes of the promoters (**Suppl. Fig. 4**). This also indicate that the effect is post-transcriptional. Additionally, the intensity of the PRP boost is dependent on the GPCR stimulation level, as seen for yEGFP output, CD19 YSD output, and CAR T cell responses (**Fig. 2-4**), and is dependent on the Ste12 transcription factor (**Suppl. Fig. 8**). As of the corresponding fold changes and linear relationship between yEGFP levels and PRP-activation/shmooing (SSC) in strains with constitutive promoters, we deduced that approximated quantification of the PRP-boost can be done in an unbiased manner via SSC (i.e. independent of e.g. yEGFP or CD19 levels).

With these hypotheses of the effect, we next investigated strains with non-constitutive PRP-regulated promoters ($P_{FUS1}$ and $P_{FIG1}$) and applied the assumption that total expression level is the product of two separate sequential effects; I) an individual transcriptional level determined by the specific promoter, which may be PRP-regulated, and II) a general PRP-induced post-transcriptional expression boost. In the case of PRP-regulated promoters, the changes in expression level are a product of both effects, meaning that there is less linear correlation between yEGFP and SSC (**Suppl. Fig. 9D**), compared to the constitutive promoters (**Suppl. Fig. 9C**). Similarly, this means that the dose-response curve for the yEGFP output, as a function of the concentration of GPCR ligand, must be a product of both effects (**Suppl. Fig. 9E**). Hence, applying the logic that SSC is an approximated quantifier of the PRP boost and that the PRP boost is a multiplicative effect, the normalization of output using SSC-values should isolate the expression pattern attributed to the promoters. Indeed, SSC-normalization of yEGFP levels decreases the linear correlation between yEGFP and SSC for $P_{FUS1}$ and $P_{FIG1}$ (**Suppl. Fig. 9F**) and enhances the dose-response model fit of the commonly applied variable slope (four-parameter) nonlinear regression (**Suppl. Fig. 9G**).

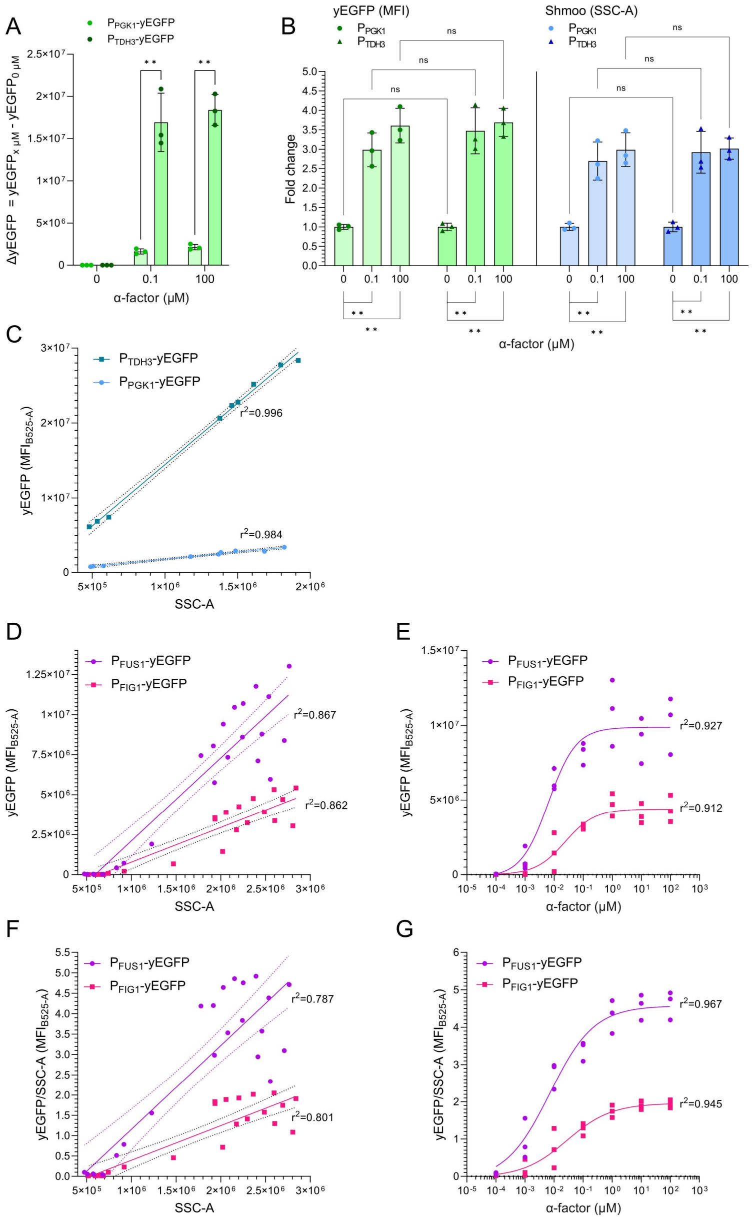


**Suppl. Fig. 9: Analysis of the PRP-induced expression boost.** The PRP boost analysis was based on flow cytometry measurements of yEGFP mean fluorescence intensity (MFI) and morphological changes (SSC) from two strains employing constitutive promoters, $P_{PGK1}$ and $P_{TDH3}$, and two strains containing PRP-regulated promoters, $P_{FUS1}$ and $P_{FIG1}$, with GPCR (Ste2) stimulation (0-100 μM α-factor). **A.** Absolute changes in yEGFP (ΔyEGFP) between $P_{PGK1}$ and $P_{TDH3}$ (constitutive promoters) relative to the background level at 0 μM (ΔyEGFP$_{0\mu M}$ = yEGFP$_X$ - yEGFP$_{0\mu M}$). **B.** Fold changes in yEGFP and SSC levels for strains with constitutive promoters. **C.** Linear relationship between yEGFP MFI and SSC for strains with constitutive promoters. **D**. Relationship between yEGFP MFI and SSC for strains with PRP-regulated promoters. **E**. yEGFP output dose-response relationship across GPCR stimulation levels with curve fitting using variable slope (four-parameter) nonlinear regression. **F**. Relationship between SSC-normalized yEGFP MFI and SSC for strains with PRP-regulated promoters. **G**. SSC-normalized yEGFP output dose-response relationship across GPCR stimulation levels with curve fitting using variable slope (four-parameter) nonlinear regression. Data represents means of median fluorescence intensity (mMFI) for three biological replicates (n=3) and standard deviations hereof. Significance levels: ✱: P≤0.05, ✱✱: P≤0.001. Not all pairwise comparisons are shown. Statistics in: **Suppl. Table 3**.

**Suppl. Fig. 10**

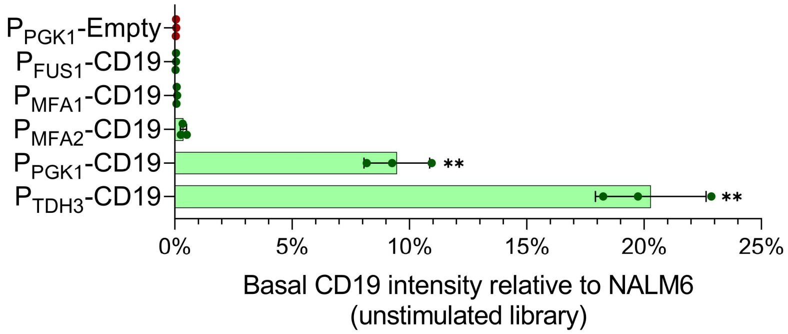


**Suppl. Fig. 10: Comparison of CD19 levels of the unstimulated SCASA yeast library to NALM6.** The basal CD19 density of SCASA yeast strains from the processing module library relative to NALM6 (100%) without stimulation of their GPCRs. Data is based on means of median fluorescence intensities (mMFI) for biological replicates (n=3). One-way ANOVA with Tukey's multiple comparisons statistical test in relation to $P_{PGK1}$-Empty is shown. Significance levels: ✱: P≤0.05, ✱✱: P≤0.001. All statistics in **Suppl. Table 5.**

### Suppl. Fig. 11

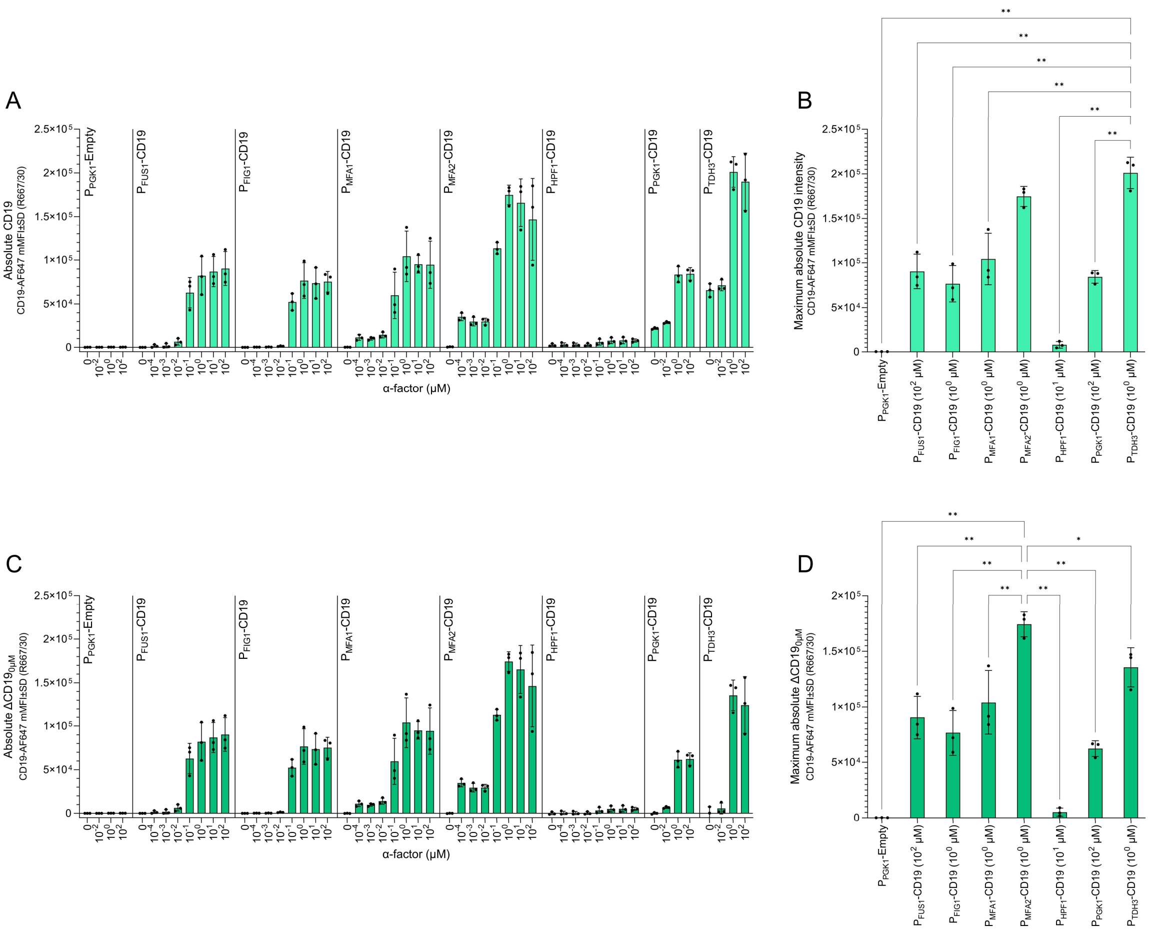


**Suppl. Fig. 11: Absolute CD19 YSD intensities upon GPCR stimulation and absolute differences herein**. **A**. absolute CD19 YSD levels for each design of the yeast strain library across different concentrations of α-factor (0-100 µM) as quantified by flow cytometry. **B**. Comparison of maximal levels of CD19 YSD between all strains for any GPCR stimulation level. **C**. Absolute changes in CD19 YSD levels calculated relative to the background level at 0 µM ($\Delta CD19_{0\mu M} = CD19_X - CD19_{0\mu M}$) for each design of the yeast strain library across different concentrations of α-factor (0-100 µM). **D**. Comparison of maximal changes in CD19 YSD levels ($\Delta CD19_{0\mu M}$) between all strains for any GPCR stimulation level. Data represents means of median fluorescence intensity (mMFI) for three biological replicates (n=3) and standard deviations hereof. Significance levels: ✱: P≤0.05, ✱✱: P≤0.001. Not all pairwise comparisons are shown. Statistics in: **Suppl. Table 5**.

### Suppl. Fig. 12

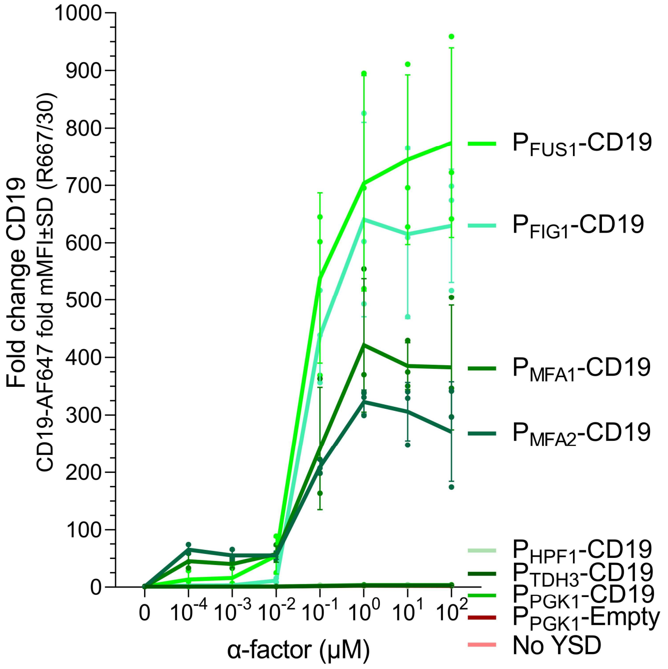


**Suppl. Fig. 12: Fold change CD19 level in SCASA yeast strains upon stimulation.** Comparison of fold changes of CD19 display levels during GPCR stimulation of the SCASA yeast processing module library. Data represents means of median fluorescence intensity (mMFI) for three biological replicates (n=3) and standard deviations hereof. Statistics in: **Suppl. Table 5** (not shown).

### Suppl. Fig. 13

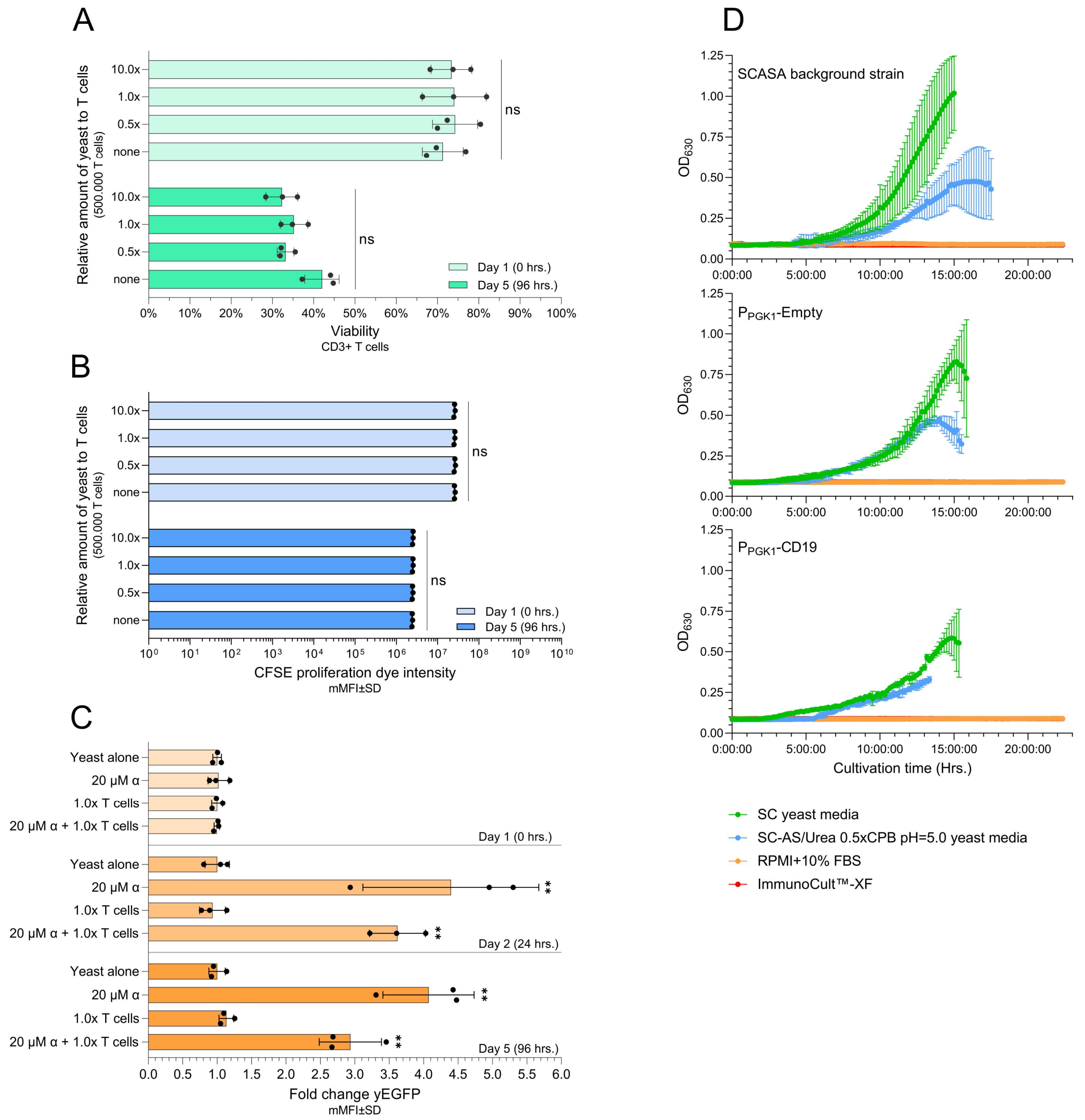


**Suppl. Fig. 13 - Examined co-cultivation conditions for human T cells and yeast.** Investigation on co-cultivation conditions for T cells isolated from a healthy donor and non-displaying SCASA yeast cells for 96 hrs. (37°C, 5%$CO_2$) in RPMI1640 glutamax + 10%FBS + 1% pen-strep without any addition of interleukins (RPMI+10%FBS). 500.000 alive CD3+ T cells were co-cultivated with yeast at three different cellular ratios; 0.5x, 1.0x, and 10.0x amounts of yeast per T cell. **A**. No significant effects were found on the viability of CD3+ T cells after co-cultivation with any number of SCASA yeast after 96 hrs., as determined by Near-IR Live/Dead staining and flow cytometry. **B**. Proliferation of CD3+ T cells was indifferent between mono-cultures and co-cultivations with any number of SCASA yeast cells after 96 hrs., as determined by staining with CFSE cell proliferation dye and flow cytometry. **C**. Fold change yEGFP output from yeast biosensors co-cultivated with T cells indicating that the yeast is alive and functional after 96 hrs. in cultivation conditions that satisfy human T cells. The biosensors are Ste2-based and sense α-factor and yEGFP was measured by flow cytometry. **D**. Three different strains of SCASA yeast were examined for growth in human cell media; the common background strain (DIX41), the negative control $P_{PGK1}$-Empty, and the CD19-displaying $P_{PGK1}$-CD19. The strains did not display the ability to grow in human cell media, RPMI+10%FBS or ImmunoCult™-XF T Cell Expansion Medium, within at least 22 hrs of cultivation at 37°C. They all displayed exponential growth within 5 hrs. in yeast media; standard synthetic complete (SC) and pH-buffered SC using ammonium sulfate and urea (SC-AS/Urea). Data is based on means of median fluorescence intensities (mMFI) (A, B, C) or $OD_{630}$ measurements (D) for three biological replicates (n=3) and standard deviations hereof. Significance levels: ✱: P≤0.05, ✱✱: P≤0.001. Statistics in: **Suppl. Table 7**

### Suppl. Fig. 14

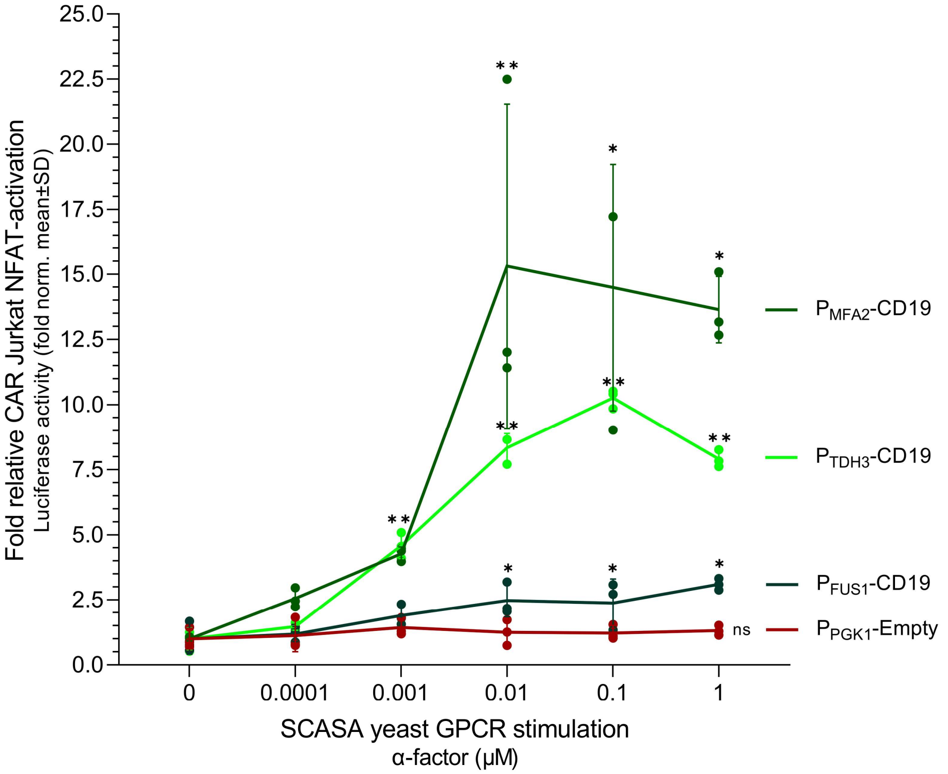


**Suppl. Fig. 14 - Fold activation of CAR Jurkat NFAT-Luc cells using GPCR-stimulated CD19+ SCASA yeast cells.** The fold change CAR Jurkat NFAT activation relative to the unstimulated condition (0 μM) for each SCASA yeast strain in co-cultures of 100% CAR+ Jurkat NFAT-Luc cells with three different SCASA yeast designs ($P_{FUS1}$-CD19, $P_{MFA2}$-CD19, and $P_{TDH3}$-CD19, *green lines*), across a concentration range of α-factor stimulation (0-1 μM) of the SCASA yeast cell GPCR, Ste2. As negative control, we used yeast lacking the CD19 CDS in the display construct ($P_{PGK1}$-Empty, *red line*). Before co-cultivation, the individual SCASA yeast cells were stimulated with α-factor for 20 hrs to induce a controlled increase in the CD19 surface display. Data is based on means of three biological replicates (n=3) and standard deviations hereof. Significance levels: ✱: P≤0.05, ✱✱: P≤0.001. Not all pairwise comparisons are shown. Statistics: **Suppl. Table 8.**

### Suppl. Fig. 15

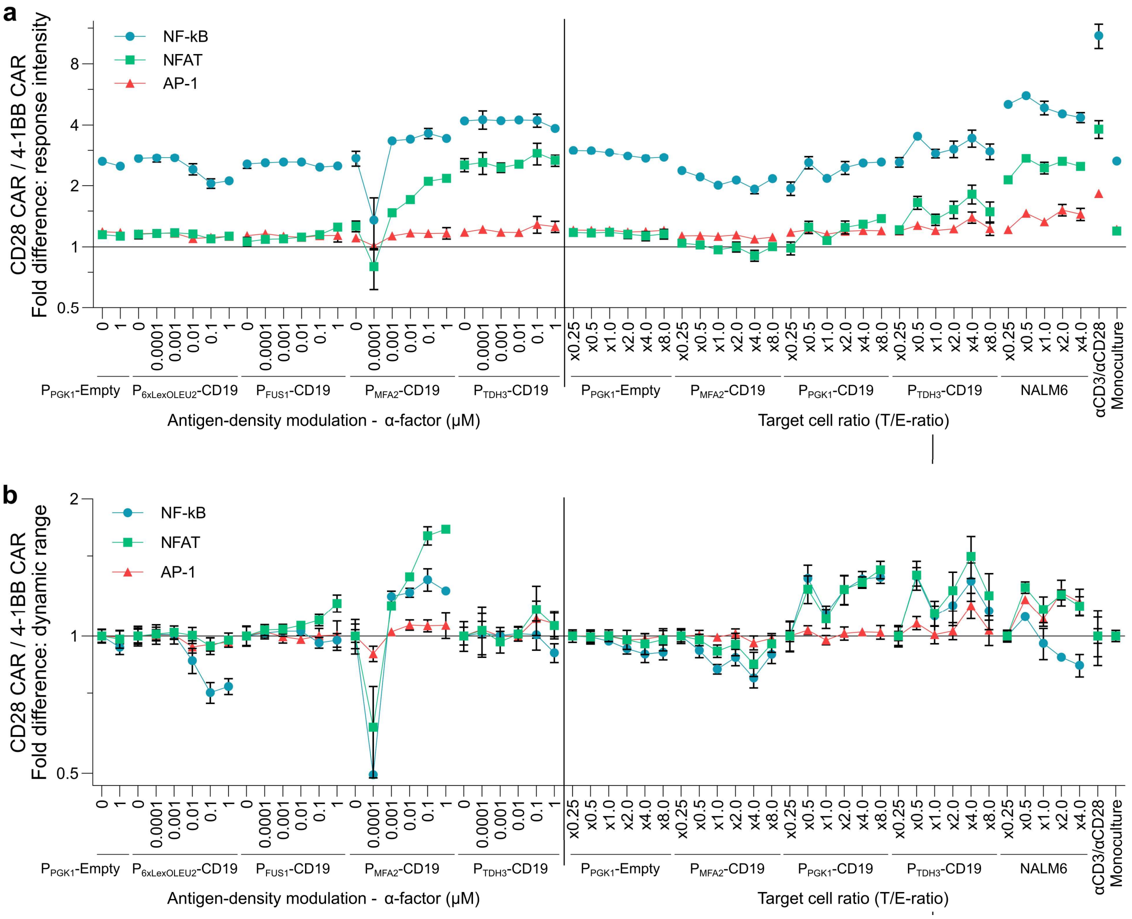


**Suppl. Fig. 15 – Direct comparison of CAR designs on NF-κB, NFAT, and AP-1 activity**. Quantification of fluorescent outputs of NF-κB-CFP, NFAT-eGFP, and AP-1-mCherry, as displayed in **Fig. 4**. Here a direct comparison of the responses of the two CAR designs, by quantification of the fold difference of the CD28 CAR over the 4-1BB CAR. **a**. Fold difference in response intensity between CARs. **b**. Fold difference in dynamic range between CARs. Data represents means of median fluorescence intensities (mMFI) for three biological replicates (n=3) and standard deviations hereof. All statistics in: **Suppl. Table 9**.

### Suppl. Fig. 16

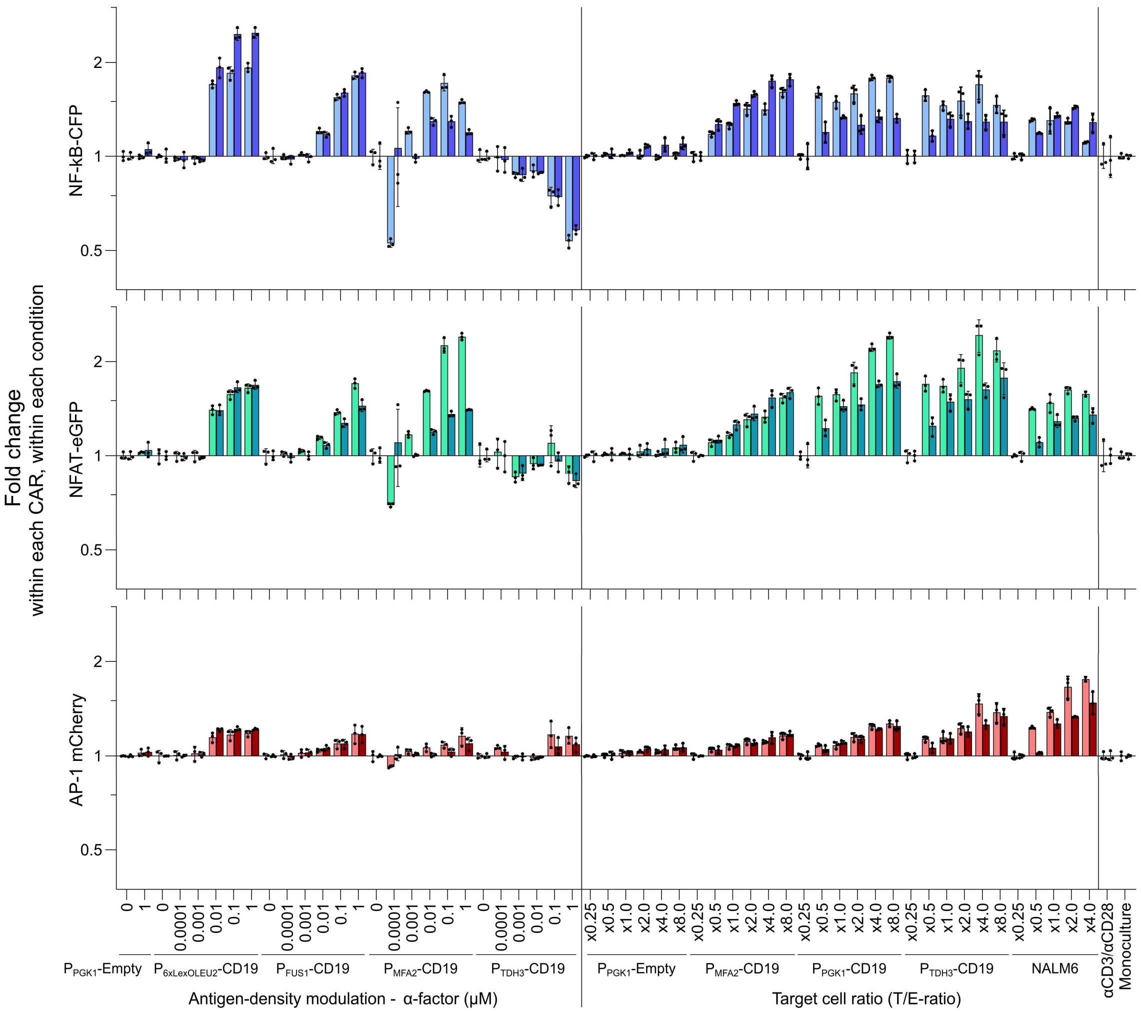


**Suppl. Fig. 16 – Fold difference of NF-κB, NFAT, and AP-1 within each condition (*separated in transcription factors*)** - Quantification of fluorescent outputs of NF-κB-CFP, NFAT-eGFP, and AP-1-mCherry, as displayed in **Fig. 4**. Here, the fold change in activity within changes in each condition relative to the lowest stimulation level (i.e. 0 µM or x0.25) is shown for each reporter (e.g. changes caused by individual strains or NALM6). This is plotted for each CAR individually; CD28 CAR (*light colors*) and 4-1BB CAR (*dark colors*) – the data is also presented in **Suppl. Fig. 17** with separation into CARs for an easier overview in that dimension. Data represents means of median fluorescence intensities (mMFI) for three biological replicates (n=3) and standard deviations hereof. All statistics in: **Suppl. Table 9**.

**Suppl. Fig. 17**

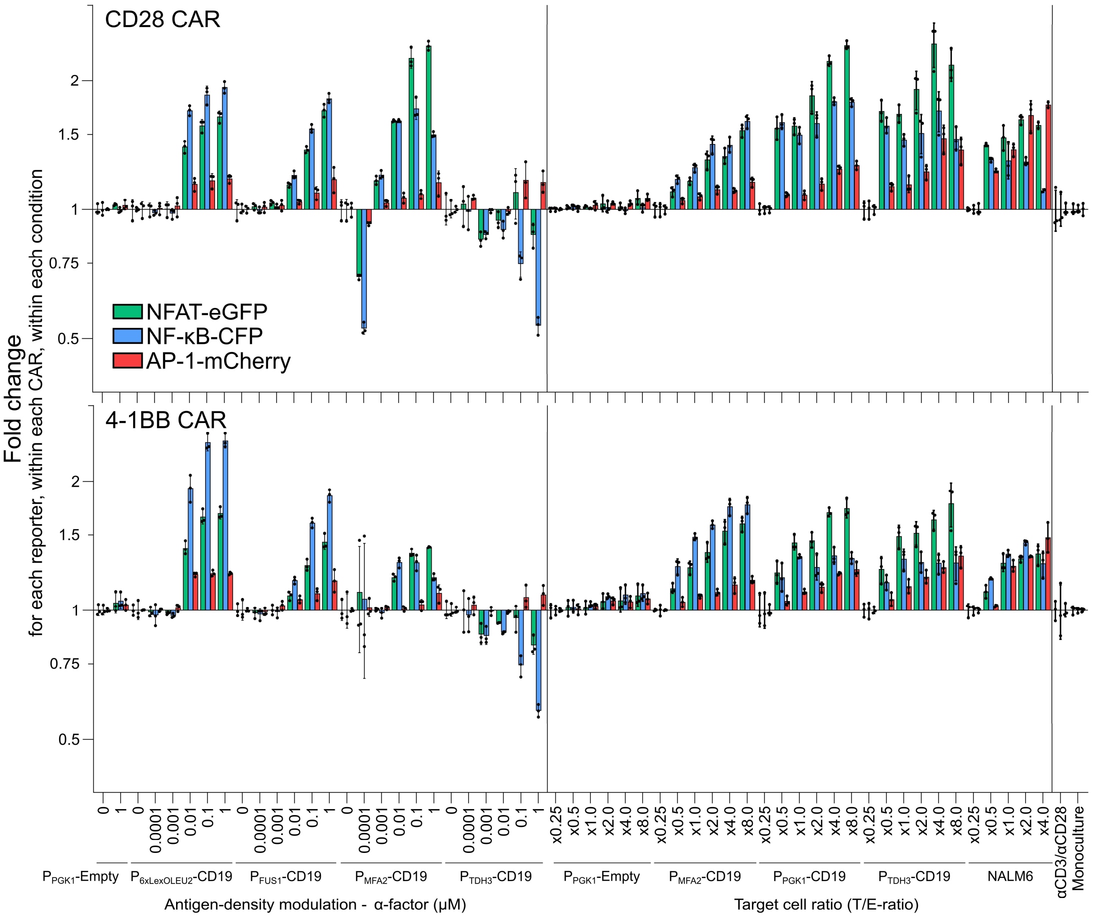


**Suppl. Fig. 17 - Fold difference of NF-κB, NFAT, and AP-1 within each condition (*separated in CAR designs*)** - Quantification of fluorescent outputs of NF-κB-CFP, NFAT-eGFP, and AP-1-mCherry, as displayed in **Fig. 4**. Here, the fold change in activity within changes in each condition relative to the lowest stimulation level (i.e. 0 μM or x0.25) is shown for each reporter (e.g. changes caused by individual strains or NALM6). The data is also presented in **Suppl. Fig. 16** with separation into transcription factors for an easier overview in that dimension. Data represents means of median fluorescence intensities (mMFI) for three biological replicates (n=3) and standard deviations hereof. All statistics in: **Suppl. Table 9**.

### Suppl. Fig. 18

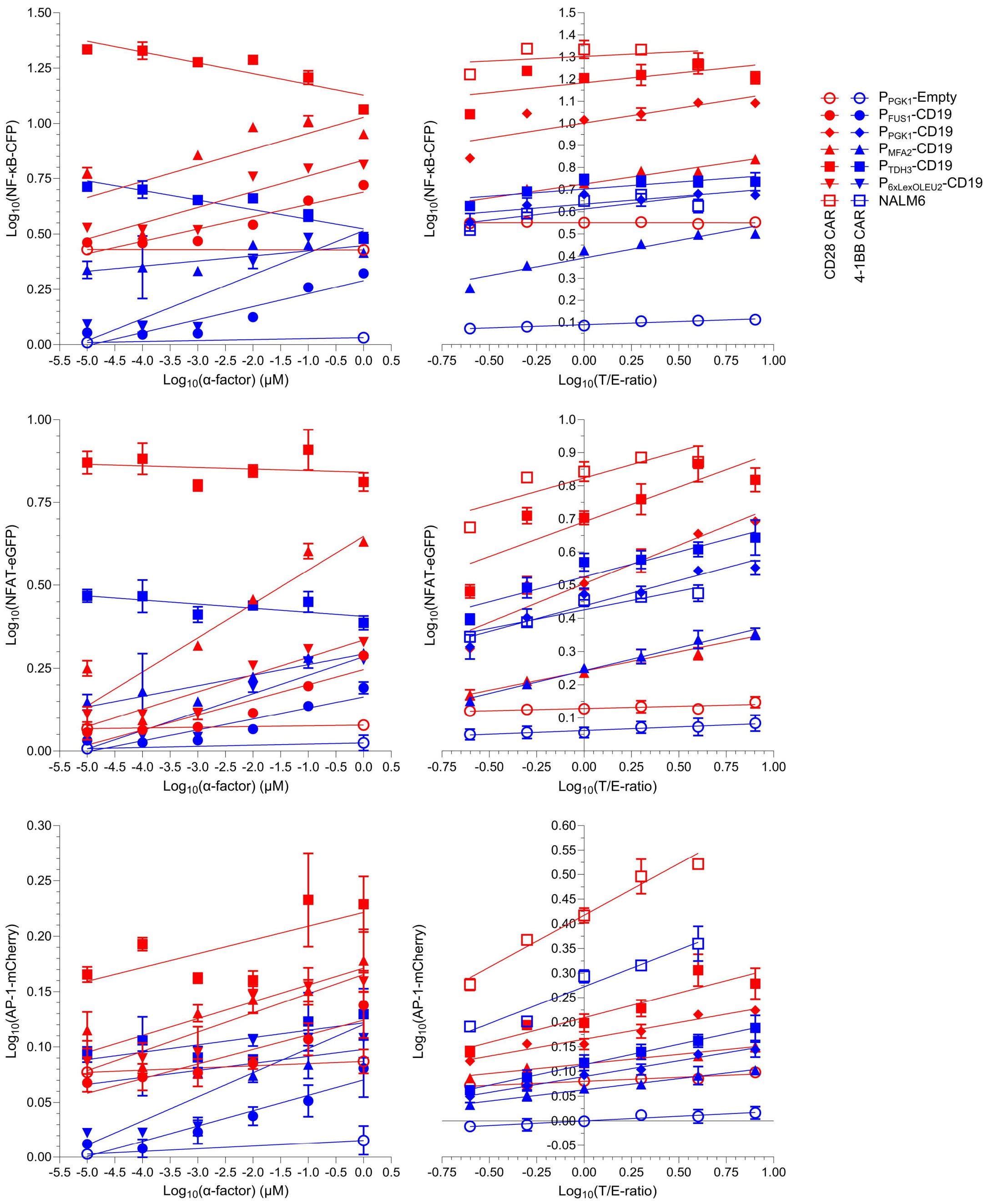


**Suppl. Fig. 18 – Linear regressions to determine responsiveness coefficients (slopes).** Data is based on the quantification of fluorescent outputs of NF-κB-CFP, NFAT-eGFP, and AP-1-mCherry, as displayed in **Fig. 4**. Linear regressions were performed for each individual combination of CAR, transcription factor, and condition (i.e. strain and stimulatory change; antigen density variation or target-to-effector (T/E) ratio). This was done to acquire the slope, which represents the responsiveness coefficients. Data represents means of median fluorescence intensities (mMFI) for three biological replicates (n=3) and standard deviations hereof. Entire analysis and statistics in: **Suppl. Table 10**.

### Suppl. Fig. 19

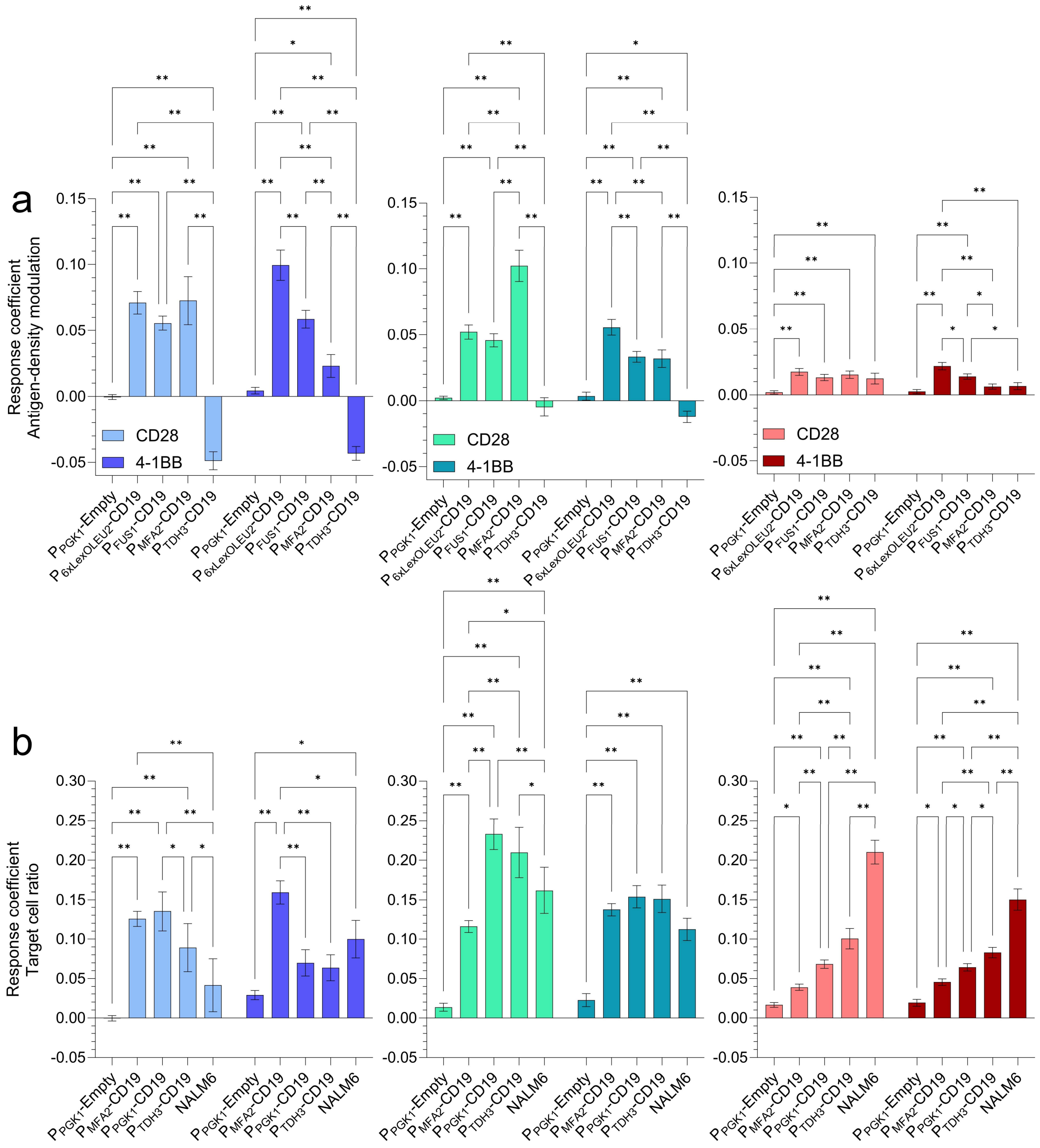


**Suppl. Fig. 19 – Responsiveness coefficients of CAR T cell designs for changes in antigen density and cellular ratios.** Data is based on the quantification of fluorescent outputs of NF-κB-CFP, NFAT-eGFP, and AP-1-mCherry, as displayed in **Fig. 4.** Responsiveness coefficients for each reporter gene and CAR design, describing the degree of response change as a function of changes in antigen density (**a**) and target-to-effector (T/E) cell ratios (**b**) for individual SCASA yeast strains and NALM6 - i.e. the response sensitivity to changes. Data represents means of median fluorescence intensities (mMFI) for three biological replicates (n=3) and standard deviations hereof. Selected comparisons from two-way ANOVAs with Šídák's multiple comparisons tests are shown. Significance levels: ✱: P≤0.05, ✱✱: P≤0.001. All statistics and extended analyses in: **Suppl. Table 10**.

### Suppl. Fig. 20

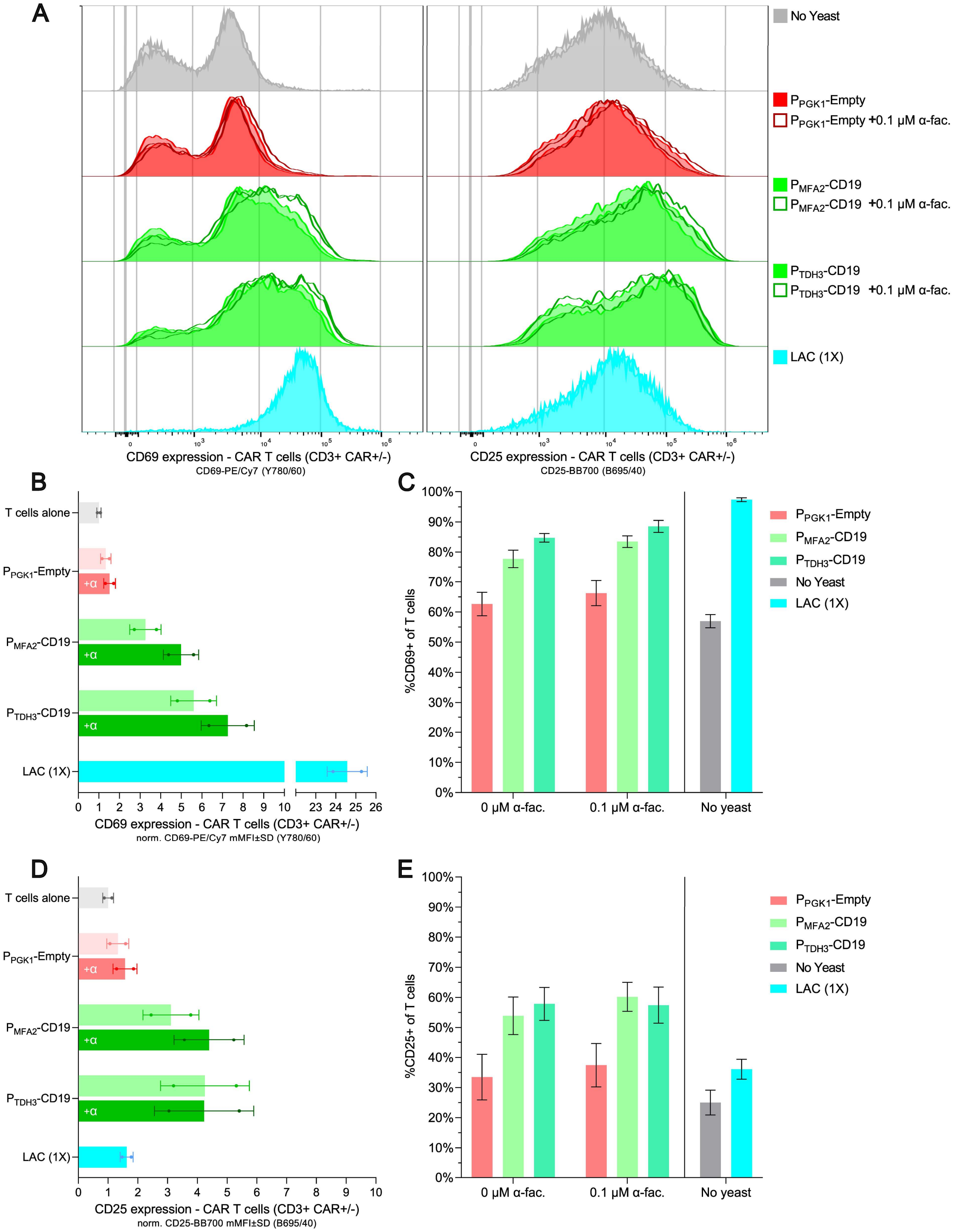


**Suppl. Fig. 20 - Activation of CAR T cells from a healthy donor after 5 months of cryopreservation using engineered SCASA yeast cells.** The Hu19-CD8α-CD28-CD3ζ CAR T cell product was manufactured from healthy donor blood using CRISPR-MAD7, whereafter the functionality was verified using SCASA yeast cells and NALM6 (**Fig. 5**). A proportion of the CAR T cell product was cryopreserved for 5 months in liquid nitrogen, and the activational profile was then re-assessed using SCASA yeast cells. Yeast cells were examined with and without GPCR pre-stimulation for 20 hrs. using 0.1 µM α-factor for increasing CD19 levels. Alive CAR T cells were cultivated in excess compared to yeast cells (0.3x yeast cell per CAR T cell) in co-cultures for 20 hrs., and subsequently, the activational profile was assessed by flow cytometric detection of CD69 and CD25 expression on

T cells using anti-CD69-PE-Cy7 (Y780/60) and anti-CD25-BB700 (B695/40), respectively. Note: the CAR T cell culture is not gated on CAR+/-. **A**. Expression of activation marker CD69 (*left*) and CD25 (*right*) in the CAR T cell culture (Alive, CD3+) after mono-cultivation without yeast (*grey*), or co-cultivation with negative control SCASA yeast $P_{PGK1}$-Empty lacking CD19 (*red*), SCASA yeast $P_{MFA2}$-CD19 (*green*), or SCASA yeast $P_{TDH3}$-CD19 (*green*), with GPCR pre-stimulation (*unfilled histograms*) or without stimulation (*filled histograms*). Additionally, a monoculture of CAR T cells was activated with lymphocyte activation cocktail (LAC) (*blue*). **B**. CD69 expression intensities of alive CD3+ CAR T cells after co-cultivation with the various SCASA yeast cells with (*+α*) and without GPCR stimulation. Intensities have been normalized to CAR T cell monocultures ('T cells alone'). **C**. Percentage of alive CD3+ CAR T cell population expressing CD69 (%CD69+) at any intensity for the co-cultivations. **D**. CD25 expression intensities of alive CD3+ CAR T cells after co-cultivation with the various SCASA yeast cells with and without GPCR stimulation, normalized to CAR T cell monocultures. **E**. Percentage of alive CD3+ CAR T cell population expressing CD25 (%CD25+) at any intensity for the co-cultivations. Data represents means of cell counts or median fluorescence intensities (mMFI) for two biological replicates (n=2) and standard deviations hereof. Histograms are normalized to the mode and all replicates are shown.

### Suppl. Fig. 21

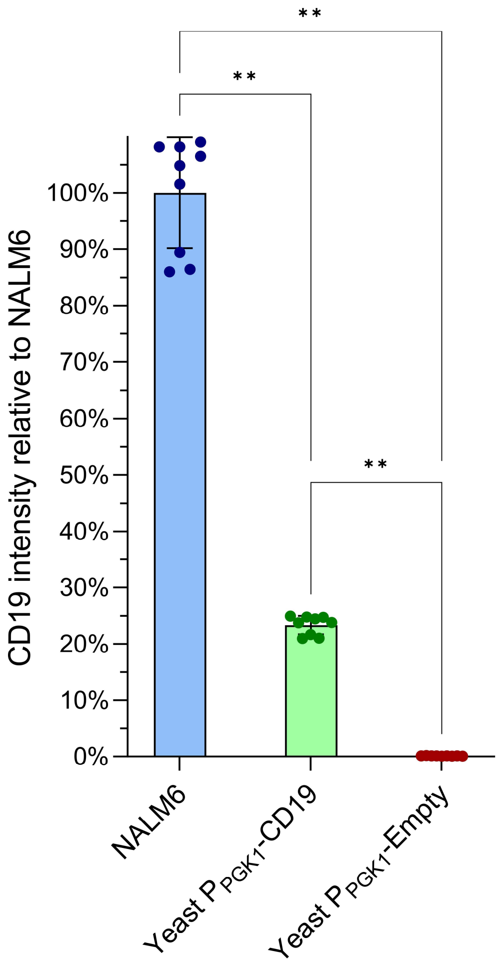


**Suppl. Fig. 21 - SCASA yeast cell $P_{PGK1}$-CD19 relative intensity of CD19 compared to NALM6 in CTRL T cell co-cultures.** The CD19 expression intensity was measured for all target cells during co-cultivation with the Hu19-CD8α-CD28-CD3ζ CAR T cell product and accompanying controls. The relative CD19 levels were determined in control (CTRL) T cell cultures via staining with anti-CD19-BV785 (V780/60) and flow cytometry. CD19 intensities per cell were normalized to NALM6. Data is based on means of 9 biological replicates (n=9) and standard deviations hereof. Significance levels: ✱: P≤0.05, ✱✱: P≤0.001. Statistics: **Suppl. Table 11**.

### Suppl. Fig. 22

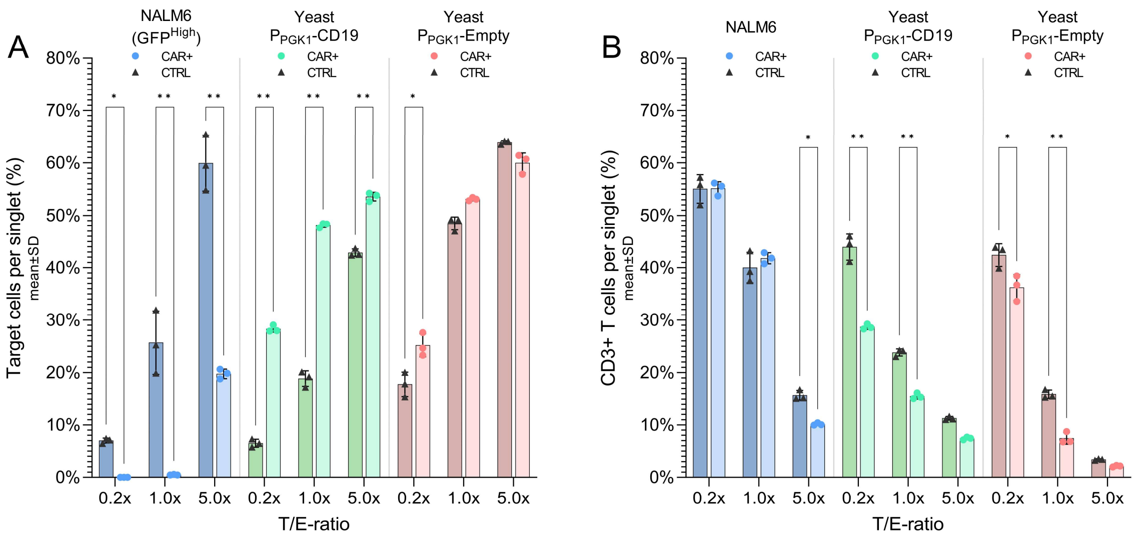


**Suppl. Fig. 22 - Cell counts after NALM6 and SCASA yeast cell co-cultivations with the Hu19-CD8α-CD28-CD3ζ CAR T cell product.** Cells were counted using flow cytometry and the number of classified cells per detected single cell (singlet) of the entire cell co-culture was calculated. **A**. Percentage of live target cells detected per singlet; NALM6 (*blue*), SCASA yeast $P_{PGK1}$-CD19 (*green*), and SCASA yeast $P_{PGK1}$-Empty (*red*) in control (CTRL) (*triangle*) and CAR (*circle*) T cell co-cultures for each target-to-effector cell ratio (T/E-ratio). **B**. Percentage of live CD3+ T cells detected per singlet in CTRL and CAR T cell co-cultures. Data is based on means of three biological replicates (n=3) and standard deviations hereof. Significance levels: ✱: P≤0.05, ✱✱: P≤0.001. Not all pairwise comparisons are shown. Statistics: **Suppl. Table 11**.

**Suppl. Fig. 23**

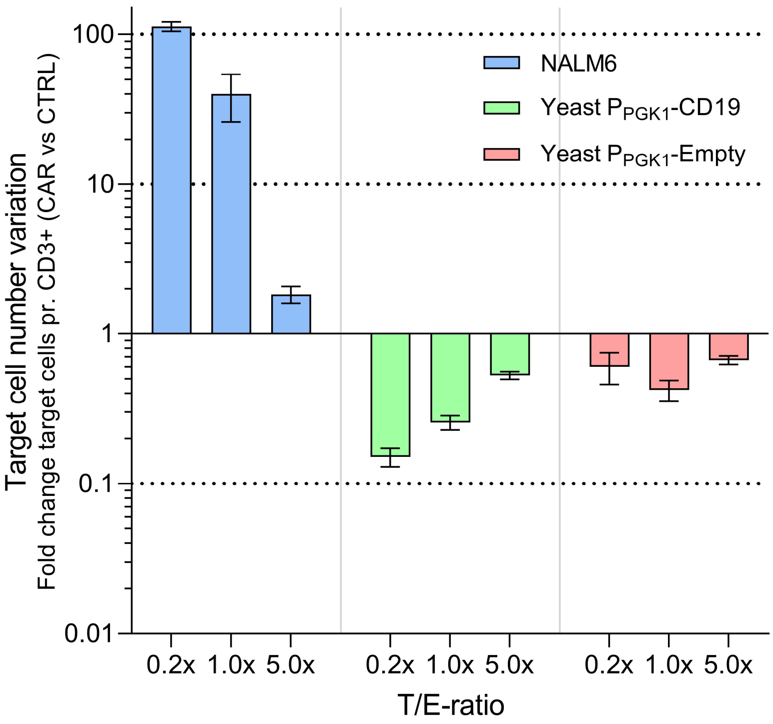


**Suppl. Fig. 23 - Characterization of target cell number stability for NALM6 and SCASA yeast cells.** The variability in target cell numbers, i.e. for NALM6 (*blue*), SCASA yeast $P_{PGK1}$-CD19 (*green*), and SCASA yeast $P_{PGK1}$-Empty (*red*), was quantified by assessing fold change in the number of detected target cells per alive CD3+ T cell between the control (CTRL) and CAR T cell cultures, for each initially seeded target-to-effector cell ratio (T/E-ratio). That is, the change in target-to-effector cell ratio caused by CAR expression in the CAR+ CD3+ T cell subpopulation for each co-cultivation setup. Here is shown the fold change variation in target-to-effector cell ratio between CTRL and CAR T cell cultures ($log_{10}$-scale). See **Fig. 5f** for the absolute fold change variation in target-to-effector cell ratio between CTRL and CAR T cell cultures, disregarding whether the variation causes increased or diminished levels of target-to-effector cells. The absolute fold change was calculated by acquiring the reciprocal value of fold changes <1. Data is based on means of three biological replicates (n=3) and standard deviations hereof.

### Suppl. Fig. 24

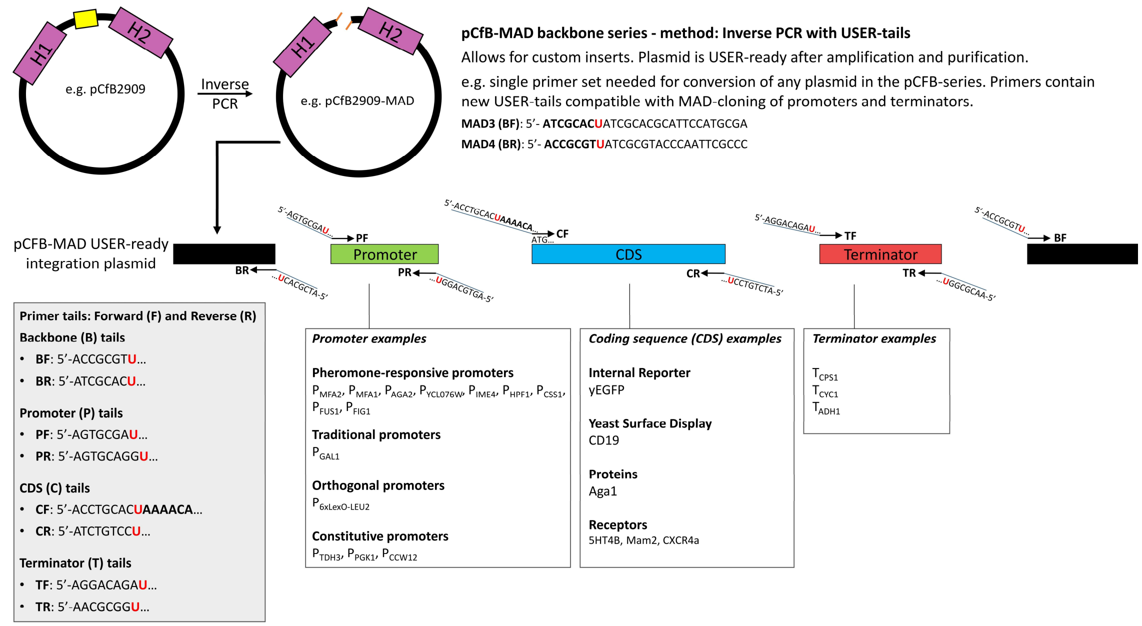


**Suppl. Fig. 24 - MAD-cloning procedure example.** Genome-integration plasmids were based on the EasyClone-MarkerFree integrative vectors (e.g. pCfB2909 for genome integration site XII-5)[3], which were inverse amplified for generating a linearized USER-ready backbone and for the removal of the EasyClone-MarkerFree cloning site structure, using the backbone primers (BF+BR) with USER-tails compatible with promoter and terminator parts. The regions for upstream homology (H1) and downstream homology (H2) are flanked by *Not*I-sites (not shown) for the release of the integration cassette. All parts were amplified using the same primer tails for the individual USER-cloning assembly points, allowing for easy exchange of different parts and library generation: promoters (PF+PR), coding sequences (CF+CR), and terminators (TF+TR). Forward primers for coding sequences were equipped with the AAAACA Kozak sequence.

### Suppl. Fig. 25

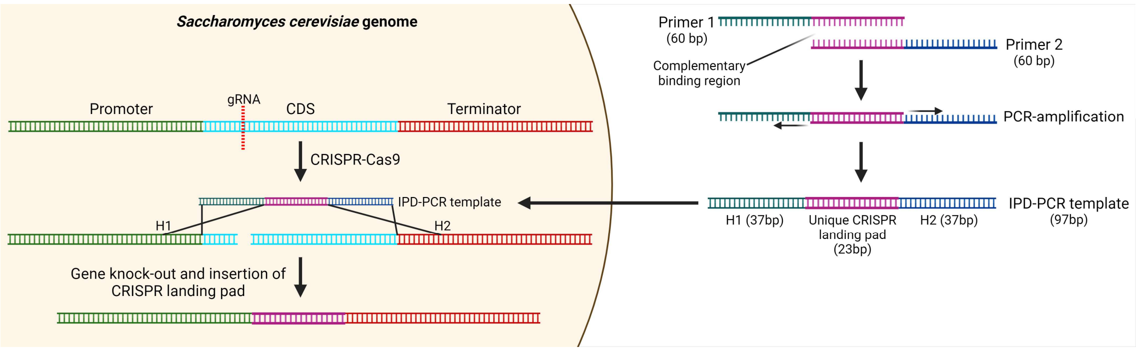


**Suppl. Fig. 25 - Intentional Primer Dimer (IPD) PCR templates for gene knockouts.** IPD-PCR templates were generated by employing two primers with a complementary binding region, which can employ each other as PCR templates for amplification. The primer tails define the genomic homology regions for targeted repair (H1 + H2) and hence flank a gRNA-targeted CRISPR-Cas9 cut site in the genome, such as for removal of exactly the CDS of a gene. In addition to allowing amplification and assembly of the IPD-PCR template, the complementary binding region also contains a 23 bp sequence comprising a unique CRISPR landing pad, which is not found in the *S. cerevisiae* genome[4], and that allows re-engineering of the site. The IPD-PCR knockout template is co-transformed with a CRISPR-Cas9 system to efficiently knockout the targeted gene.

### Suppl. Fig. 26

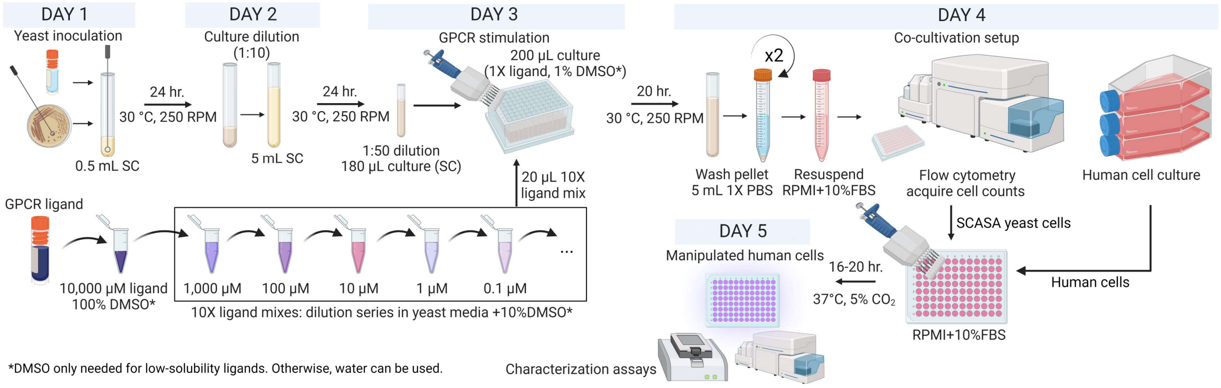


**Suppl. Fig. 26 - Example procedure for preparing SCASA yeast cells with varied GPCR stimulation for co-cultivation with human cells.** An example of a SCASA-based protocol for co-cultivation of human cells with SCASA yeast cells presenting signal molecules at different levels caused by stimulation with different amounts of GPCR ligand. Initially, yeast is inoculated and pre-cultured before GPCR stimulation. The SCASA yeast cells can be systematically exposed to a dilution series of GPCR ligands and stimulated for 20 hrs., which allows for the establishment of stable yeast surface display levels. Alternatively, built-in differences in the designs of SCASA yeast cells are used to vary the output intensity without the use of stimulation. After stimulation, the yeast cells are washed in PBS twice to prepare them for co-cultivation in human cell media, such as RPMI+10%FBS. In the case of this study, flow cytometry was employed to verify GPCR stimulation and count cells for the establishment of the co-cultures. The SCASA yeast cells and human cells can be combined in different cell-to-cell ratios to further diversify the intensity of interactions between the cells. After co-cultivation, the response of the human cells can be characterized e.g. via immunoassays or immunophenotyping. The duration of GPCR stimulation can vary between types of SCASA yeast cell inputs and outputs, and co-cultivation duration should be adapted for different cell types and response types.

### Suppl. Fig. 27

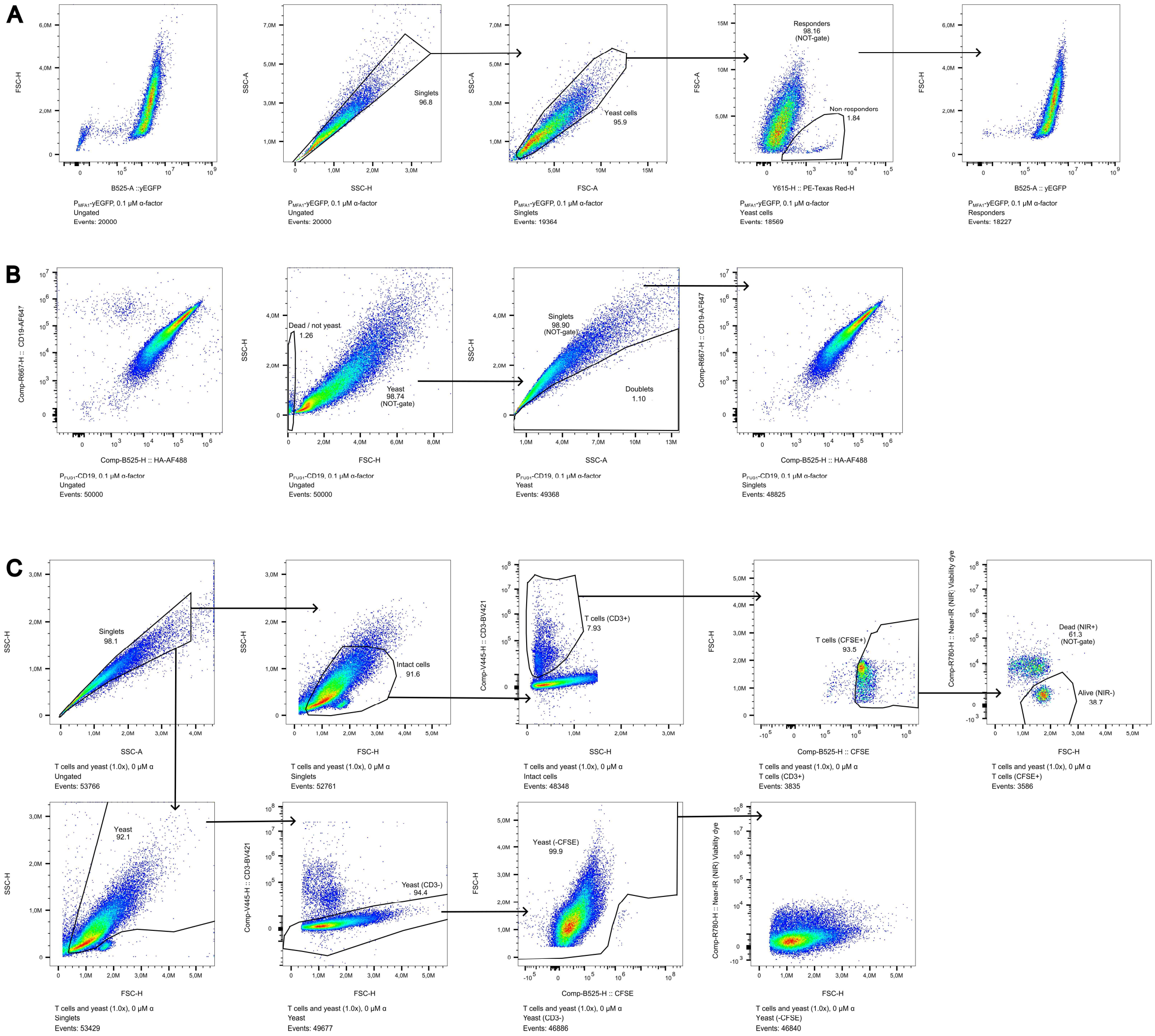


**Suppl. Fig. 27 - Gating strategies for SCASA yeast characterization and co-culture conditions.** Gating and compensation were done using FlowJo™ v10.8.1 Software (BD Life Sciences). Gating strategies employed fluorescence minus one (FMO) controls, as well as the employment of negative controls where possible for determining true positive signals (e.g. yEGFP+, CFSE+). **A**. Gating strategy for yEGFP experiments (**Fig. 2**). First singlets were gated ('Singlets'), then yeast cells were gated for removal of cellular debris and non-cellular events ('Yeast cells'), then dying or non-responding yeast cells were removed through a NOT-gate, via their distinct morphology and increased red autofluorescence ('Non-responders'), leaving responsive yeast cells for detection of yEGFP and morphological changes ('Responders'). The example is $P_{MFA1}$-yEGFP stimulated with 0.1 µM α-factor. **B**. Gating strategy for CD19 YSD (**Fig. 2**). First yeast cells were gated by a NOT-gate removing cellular debris and non-cellular events ('Yeast'), then singlets were gated by a NOT-gate ('Singlets'). The example is $P_{FUS1}$-CD19 stimulated with 0.1 µM α-factor. **C**. Gating for examination of viability and proliferation of T cells in yeast co-cultures (**Suppl. Fig. 13**). First singlets were gated ('Singlets'), then for the gating of T cells, intact cells were gated for removal of cellular debris and non-cellular events ('Intact cells'), which were then gated for their expression of CD3 ('T cells (CD3+)'). T cells were then gated for CFSE proliferation dye staining ('T cells (CFSE+)'), and dead and alive cells were identified based on Near-IR (NIR) viability dye ('Dead (NIR+)' / 'Alive (NIR-)'). For yeast cells, yeast was first identified based on morphological traits ('Yeast'), then on their lack of CD3 expression ('Yeast (CD3-)'), and finally their lack of CFSE staining ('Yeast (-CFSE)'). The example is co-cultivation of T cells with 1.0x yeast, at 0 µM alpha-factor, on Day 5 (96 hrs).

### Suppl. Fig. 28

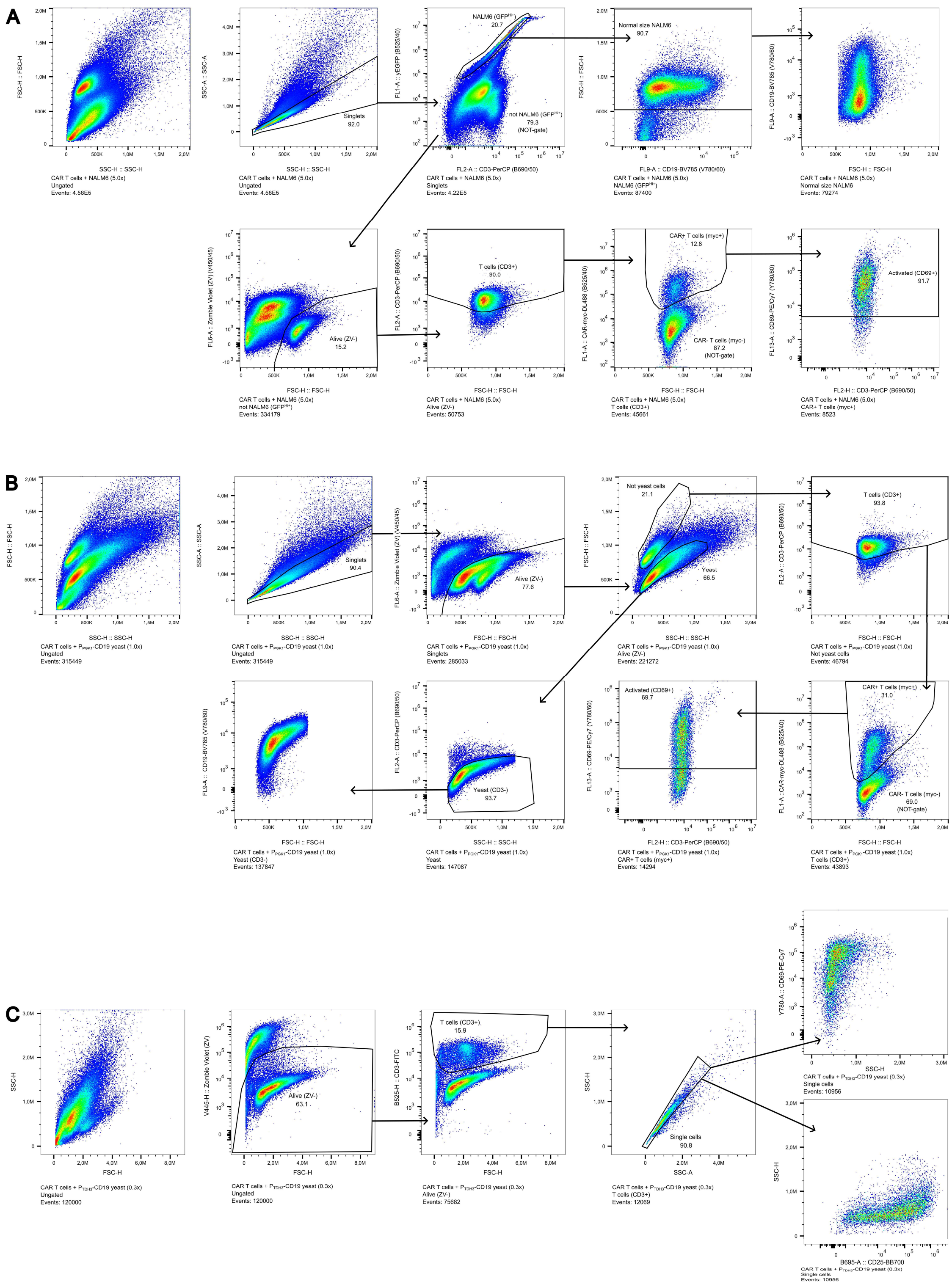


**Suppl. Fig. 28 - Gating strategies for CAR T cell activation assays.** Gating and compensation were done using FlowJo™ v10.8.1 Software (BD Life Sciences). Gating strategies employed fluorescence minus one (FMO) controls, as well as the employment of negative controls where possible for determining true positive signals (e.g. yEGFP+, CAR+). **A**. Gating strategy for activation of donor-derived CAR T cells using NALM6 (**Fig. 5**). First singlets were gated ('Singlets'), then for separation of NALM6, the GFP expressing population was gated ('NALM6 ($GFP^{Hi+}$)') and further gated by size ('Normal size NALM6'). For the separation of T cells, a NOT-gate was employed for the NALM6 population ('not NALM6 ($GFP^{Hi+}$)'). Then the alive population was gated by a Zombie Violet (ZV) stain ('Alive (ZV-)'). T cells were then identified by CD3 expression ('T cells (CD3+)'), and from here CAR-expressing cells were identified through myc-tag staining ('CAR+ T cells (myc+)'), and non-expressing T cells through a NOT-gate ('CAR- T cells (myc-)'). Activated cells were identified by a CD69 stain ('Activated (CD69+)'). The same strategy was employed for the CTRL T cell culture. The example is a co-cultivation of CAR T cells with 5.0x NALM6. **B**. Gating strategy for activation of donor-derived CAR T cells using SCASA yeast (**Fig. 5**). First singlets were gated ('Singlets'), then the alive population was gated by a Zombie Violet (ZV) stain ('Alive (ZV-)'). Hereafter, yeast cells ('Yeast') could be separated from human cells ('Not yeast cells') through morphological traits. T cells were then identified via CD3 expression ('T cells (CD3+)'), and from here CAR-expressing cells were identified through myc-tag staining ('CAR+ T cells (myc+)'), and non-expressing T cells through a NOT-gate ('CAR- T cells (myc-)'). Activated cells were identified by a CD69 stain ('Activated (CD69+)'). Yeast cells were further identified from the lack of CD3 expression ('Yeast (CD3-)') for the investigation of CD19 levels. The same strategy was employed for the CTRL T cell culture and the negative control yeast ($P_{PGK1}$-Empty). The example is a co-cultivation of CAR T cells with 1.0x $P_{PGK1}$-CD19 SCASA yeast. **C**. Gating strategy for post-cryopreservation activation of donor-derived CAR T cells using SCASA yeast (**Suppl. Fig. 20**). First the alive population was gated by a Zombie Violet (ZV) stain ('Alive (ZV-)'), then T cells were then identified by CD3 expression ('T cells (CD3+)'), and lastly singlets were gated ('Single cells') for the examination of CD69 and CD25 expression. The same strategy was employed for the other SCASA yeast strains and LAC control. The example is a co-cultivation of CAR T cells with $P_{TDH3}$-CD19 SCASA yeast (0.3x) without α-factor.

### References


1. Jensen, E. D. *et al.* Engineered cell differentiation and sexual reproduction in probiotic and mating yeasts. *Nat. Commun.* **13**, 6201 (2022).
2. Shaw, W. M. *et al.* Engineering a Model Cell for Rational Tuning of GPCR Signaling. *Cell* **177**, 782–796.e27 (2019).
3. Jessop-Fabre, M. M. *et al.* EasyClone-MarkerFree: A vector toolkit for marker-less integration of genes into Saccharomyces cerevisiae via CRISPR-Cas9. *Biotechnol. J.* **11**, 1110–1117 (2016).
4. D'Ambrosio, V. *et al.* A FAIR-compliant parts catalogue for genome engineering and expression control in Saccharomyces cerevisiae. *Synth Syst Biotechnol* **7**, 657–663 (2022).
